# PDE3A-SLFN12 Molecular Glues Target Multiple KIT D816V Cell Types in Preclinical Models of Mast Cell Malignancies

**DOI:** 10.64898/2026.08.13.744624

**Authors:** Xuhuang Fu, Anne Kaiser, Payal Chawla, Axel Choidas, Peter Habenberger, Tiago Maié, Alessia Piergentili, Venkatakrishnan Hariharan, Paul Wanek, Susanne Schmitz, Maximilian Ackermann, Deborah Christen, Jens Panse, Hubert Schorle, Michel Arock, Heidi Greulich, Giulia Rossetti, Steffen Koschmieder, Ivan G Costa, Tim H Brümmendorf, Marcelo A S Toledo, Bert M Klebl, Martin Zenke

## Abstract

A drug discovery approach was used to specifically target malignant cells with *KIT* D816V mutation, which is the predominant disease-causing mutation in clonal mast cell malignancies. To this end, KIT D816V cells derived from induced pluripotent stem cells (iPS cells) of KIT D816V patients were employed to screen a library of FDA approved and experimental drugs for specific killing of KIT D816V cells. We discovered the novel compound LDC 3416, which targets multiple malignant KIT D816V cell types, including hematopoietic stem/progenitor cells and mast cells. Importantly, by exploring the LDC 3416 targeting profile, we identified the phosphodiesterase 3A-*Schlafen* 12 (PDE3A-SLFN12) molecular glue pathway as a novel approach for specific targeting of malignant KIT D816V cells. We found that the KIT D816V mutant protein leads to increased expression of PDE3A and SLFN12 and thus confers a selective molecular vulnerability to PDE3A-SLFN12 molecular glues. Primary malignant mast cells of KIT D816V patients with indolent and advanced systemic mastocytosis also exhibit increased expression of PDE3A and SLFN12. We extended our study to include additional PDE3A-SLFN12 molecular glues and demonstrate their synergistic action with KIT D816V selective tyrosine kinase inhibitors (TKIs) in killing KIT D816V cells. Furthermore, the PDE3A-SLFN12 molecular glues also target KIT D816V megakaryocytes, a cell type that has been underestimated in malignant mast cell pathophysiology and molecular targeting. The identified molecular glues, along with their synergy with TKIs and their simultaneous targeting of multiple KIT D816V cell types, open novel treatment options for KIT D816V mast cell malignancies and other KIT D816V associated diseases.

## Introduction

Malignant mast cell disorders are devastating, often chronic progressive diseases that are mostly caused by acquired mutations in the *KIT* gene. The most common mutation found in malignant mast cell disorders is *KIT* D816V, which results in constitutive activation and signaling of the stem cell factor (SCF) receptor tyrosine kinase KIT (Arock et al., 2018; Akin et al., 2025). Indeed, the *KIT* D816V mutation is found in >85% of patients with systemic mastocytosis (SM) and causes neoplastic mast cell expansion and accumulation in affected tissues and organs (Akin et al., 2025; Hoermann et al., 2026). *KIT* mutations, including *KIT* D816V, are also driver mutations in other malignancies, such as gastrointestinal stromal tumors (GIST) and acute myeloid leukemia (AML; Craig et al., 2020; Klug et al., 2022).

Frequently, drugs such as tyrosine kinase inhibitors (TKIs), directly target the aberrantly activated signaling molecules in a well-defined pocket. Accordingly, several broad and selective TKIs effectively inhibit KIT D816V tyrosine kinase signaling (Arock et al., 2018; Valent et al., 2023; Akin et al., 2025). The KIT D816V selective TKI Avapritinib is now used for SM therapy, including indolent SM (ISM) and advanced forms of SM (advSM), such as aggressive SM (ASM), SM with an associated hematological neoplasm (SM-AHN) and mast cell leukemia (MCL) (DeAngelo et al., 2021; Gotlib et al., 2021; Reiter et al., 2022; Gotlib et al., 2023; 2026). Avapritinib shows an overall response rate of 75%, including several patients achieving complete remissions, some of which persisting long-term, yet data on overall long-term outcome are limited. Despite the central role of KIT D816V, disease persistence or relapse are influenced by multilineage involvement, cooperating mutations (e.g. *TET2*, *SRSF2*, *ASXL1* and *RUNX1*), inflammatory signaling and stem cell persistence, highlighting the need for further therapeutic strategies. So far, the only proven curative SM therapy, especially for TKI refractory patients and patients suffering from side effects under TKI therapy, is allogeneic hematopoietic stem cell transplantation. Accordingly, allogeneic stem cell transplantation is recommended in eligible advSM patients, yet can be associated with significant morbidity and transplant-related mortality, and the relapse-related mortality is 15-23%, depending on the SM subtype (Valent et al., 2023; Lübke et al., 2024; McLornan et al., 2024).

Thus, there is a clear need to explore further targets and/or synergistic treatment strategies to control and potentially cure SM. Here we embarked on the hypothesis to target the KIT D816V cancer cell phenotype in an unbiased drug discovery approach and to identify non-TKI compounds that selectively kill KIT D816V cells and potentially synergize with TKIs.

Human-centric new approach methodologies (NAMs) are currently receiving major attention in drug development for assessing drug efficacy, safety and quality, and to overcome the high failure rates observed in traditional drug discovery (Liu et al., 2026). In this context, disease-specific induced pluripotent stem cells (iPS cells) derived from patient cells represent a particularly appealing approach for disease modelling and *in vitro* drug discovery (Yamanaka, 2026). iPS cells and the cells derived thereof capture the mutational landscape and genomic profile of the patient and can be expanded as clonal and homogenous cell populations to high cell numbers. Here we employed KIT D816V iPS cells and isogenic controls without KIT D816V mutation from patients with ASM and MCL as a patient-specific SM disease model for drug discovery. Drug screening identified several novel compounds that selectively killed KIT D816V cells, including phosphodiesterase 3A (PDE3A) modulators that can act as molecular glues.

Molecular glues represent a new class of cell selective small molecules that induce proximity between two or more proteins and thereby generate enhanced and/or new functionalities (King et al., 2026). For example, the molecular glues studied here bind to PDE3A homodimers and thereby recruit two Schlafen 12 (SLFN12) monomers to form PDE3A-SLFN12 heterotetramers (Chen et al., 2021; Garvie et al., 2021; Yan et al., 2022; Lee et al., 2023; Greulich, 2024). This leads to activation of SLFN12 RNase activity, tRNA Leu (TAA) cleavage and apoptosis. Molecular glues offer advantages over traditional inhibitors, which often require continuous high dosing and are more prone to resistance (Buntz, 2025). In addition, compared to bivalent degraders, such as PROTACs, molecular glues are small molecules, allowing better permeability into cells and easier synthesis and large-scale manufacturing (Mullard, 2024; King et al., 2026).

Here we discovered a novel PDE3A modulator, LDC 3416, which acts as a molecular glue and induces selective killing of iPS cell-derived KIT D816V hematopoietic stem/progenitor cells, mast cells and megakaryocytes of SM and MCL patients. By exploring the LDC 3416 targeting profile, we identified the PDE3A-SLFN12 molecular glue pathway as a novel approach for effective targeting malignant KIT D816V cells. We found that the PDE3A-SLFN12 molecular glues Anagrelide, BRD 9500 and BAY 2666605 also induced selective killing of KIT D816V cells. Additionally, PDE3A-SLFN12 molecular glues acted in synergy with the KIT D816V selective TKI Avapritinib and Nintedanib and enhanced KIT D816V cell killing.

## Materials and Methods

### Culture of iPS Cells and their Differentiation into Hematopoietic Stem/Progenitor Cells, Mast Cells and Megakaryocytes

Patient-specific iPS cells with KIT D816V mutation and without mutation, referred to as KIT D816V iPS cells and control iPS cells, respectively, were from ASM patient 1 and MCL patient 3 as described (Toledo et al., 2021) here referred to as patient 1 and 2. Details on iPS cell culture and differentiation into hematopoietic stem/progenitor cells, mast cells and megakaryocytes (Supplementary Figure 1A-C), culture of ROSA and HMC-1 cells, flow cytometry analysis and magnetic activated cell sorting are in the Supplementary Material and Methods section.

### Compound Library Screening and Testing by CellTiter-Glo and MTT Assay

A compound library of 4330 FDA-approved and experimental drugs, containing also 80 compounds of the Merck Biopharma Mini Library, was screened at 10 µM final concentration with KIT D816V hematopoietic stem/progenitor cells derived from KIT D816V patient iPS cells and KIT D816V human ES cells (Toledo et al., 2021; here referred to as KIT D816V cells) employing CellTiter-Glo Luminescent Cell Viability Assay (Promega, Madison, WI) in 384- and 1536-well microtiter plates (Supplementary Figure 1D-E). Cells derived from isogenic iPS cells and ES cells without KIT D816V mutation were used as control (KIT control). Details on hit validation and dose-response curves are described in the Supplementary Material and Methods section.

### scRNA-Seq Data Analysis and Gene Expression Analysis by RT-qPCR

scRNA-seq data are from ISM and SM-AHN patients (Söderlund et al., 2023; Huang et al., 2024; GEO data base accession numbers GSE222830 and GSE249445, respectively) and were processed as described in the Supplementary Material and Methods section. RNA was isolated with NucleoSpin RNA kit (Macherey-Nagel, Düren, Germany), reverse transcribed into cDNA using random primers and MultiScribe Reverse Transcriptase (Invitrogen, Waltham, MA, USA) and subjected to quantitative PCR (qPCR) analysis as described in the Supplementary Material and Methods section.

### Molecular Docking

Molecular docking studies for ligands LDC 3416, BRD 9500 and BAY 2666605 and for Trequinsin, as a negative control, were performed using the Schrödinger 2023-1 suite (Glide, Schrödinger, 2023). For details we refer to the Supplementary Material and Methods section.

### Western Blotting, Immunofluorescence Analysis, Apoptosis and CFU Assays

For details on Western blotting, immunofluorescence analysis, apoptosis and CFU assays see the Supplementary Material and Methods section.

### Animal Experiments

The SCL-GFP-KIT D816V mouse model (Pelusi et al., 2017) and xenotransplantation of human ROSA^KIT^ ^D816V^ cells into non-obese diabetic severe combined immunodeficiency gamma (NSG) mice are described in the Supplementary Material and Methods section.

### Statistical Analysis

All data are expressed as the mean ± SD of at least three independent experiments or displayed according to specific criteria as described. Statistical analysis was performed using the Welch’s t-test or Welch ANOVA using GraphPad Prism 11 software. p values ≤0.05 considered statistically significant are indicated by *; ** p <0.01; *** p <0.001; **** p <0.0001. IC50 values were calculated by analyzing nonlinear regression with GraphPad Prism 11. For the animal experiments: Brown–Forsythe and Welch one-way ANOVA followed by Dunnett T3 post-hoc tests; for non-normally distributed data: Mann–Whitney test and Kruskal–Wallis test with Dunn’s multiple comparisons: not significant, all non-indicated statistics; * p <0.05; ** p <0.01; *** p <0.001; **** p <0.0001 using GraphPad Prism 11 software.

## Results

### LDC 3416 Targets KIT D816V Cells

A compound library comprising 4330 FDA-approved and experimental drugs was screened with KIT D816V hematopoietic stem/progenitor cells from patient KIT D816V iPS cells (in the following referred to as KIT D816V cells; Supplementary Figure 1A-E) with the objective to identify non-TKI compounds targeting KIT D816V cells. We employed multiple replicates of KIT D816V iPS cell lines from two ASM and MCL patients (referred to as patient 1 and 2) and CRISPR engineered KIT D816V human ES cells along with their respective isogenic controls (referred to as KIT control; Toledo et al., 2021). The initial screen with patient 1 KIT D816V cells identified 335 compounds, which reduced viability of KIT D816V cells by more than 50% of vehicle control (Supplementary Figure 1D and E). Compounds were further processed to select for prototypic compounds. In the end 234 compounds were subjected to hit validation in dose response assays employing KIT D816V cells from iPS cells of patient 1 and 2 and from KIT D816V human ES cells (Figure 1A, Supplementary Figure 1F, Supplementary Table 1).

**Figure 1.**
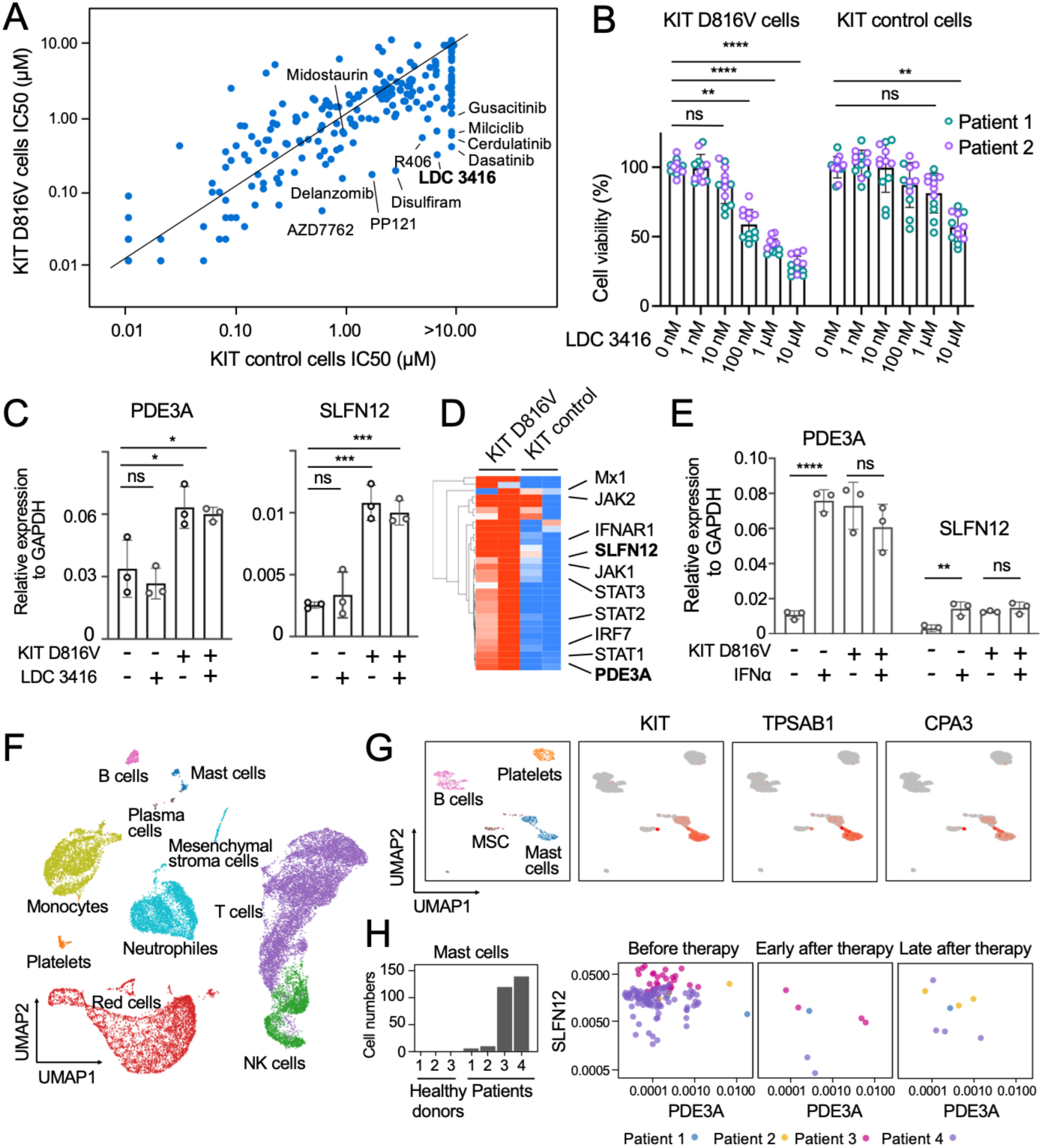
LDC 3416 compound targets KIT D816V cells and PDE3A/SLFN12 expression in KIT D816V cells from iPS cells and in KIT D816V mastocytosis patient samples. (A) KIT D816V cells and KIT control cells derived from iPS cells (patient 1) were subjected to compound screening and IC50 values upon drug treatment are shown. IC50 values above 10 µM were not resolved (>10.00). LDC 3416 is highlighted. Some kinase inhibitors, the proteasome inhibitor Delanzomib, and the aldehyde dehydrogenase inhibitor Disulfiram are shown as references. (B) Dose response curves for LDC 3416 on KIT D816V cells of patient 1 and 2 (n=6, 3 independent experiments). KIT control cells, isogenic cells of patient 1 and 2 without KIT D816V mutation. DMSO, vehicle control (0 nM). Data are represented as mean ± SD of triplicates. Two-way ANOVA, ** p<0.01, *** p<0.001, and **** p<0.0001 in comparison to no compound treatment (0 nM); ns, not significant. (C) KIT D816V cells show elevated PDE3A and SLFN12 mRNA expression compared to KIT control cells, which is unaffected by LDC 3416 treatment. Data are by quantitative RT-PCR. Cells with and without KIT D816V mutation, and with and without LDC 3416 treatment are shown as indicated (n=3). DMSO, vehicle control. One-way ANOVA, PDE3A, ** p=0.007 and SLFN12, *** p=0.0008; ns, not significant. (D) IFN signature genes expressed in KIT D816V mast cells and KIT control mast cells are subjected to bidirectional hierarchical clustering and depicted in heat map format (red, high expression; blue, low expression). Expression of PDE3A and SLFN12 and selected genes is indicated (n=2 for both KIT D816V cells and KIT control). Data extracted from Toledo et al., 2023; GSE223883. (E) IFNα treatment (2 days, 1000 U/ml) increases PDE3A and SLFN12 mRNA expression in KIT control cells to levels which are similar to levels in KIT D816V cells (n=3). *KIT* D816V mutation and IFNα treatment are indicated. Welch’s t-test, PDE3A, **** p<0.0001 and SLFN12, ** p=0.008; ns, not significant. (F) UMAP of scRNA-seq data of four KIT D816V patients with advanced SM with associated hematologic neoplasm (SM-AHN) and three healthy donors. Data extracted from Huang et al., 2024; GSE249445; annotated cell populations are indicated. (G) UMAP of scRNA-seq data of (F) representing mast cells, mesenchymal stroma cells (MSC), platelets and B cells. Mast cell marker expression of CD117/KIT, mast cell tryptase (TPSAB1) and carboxypeptidase A3 (CPA3) is shown in red. (H) Frequency of mast cells in white blood cells of SM-AHN patients (Patients 1-4) and healthy donors (Donors 1-3) from scRNA-seq data of (F) (left panel). PDE3A and SLFN12 expression in mast cells of patients 1-4 before Avapritinib therapy and early (20-120 days) and late (200-400 days) after Avapritinib therapy (GSE249445) (right panels).

The compound LDC 3416 sparked our interest, since it is positioned in the effector zone together with multiple kinase inhibitors (Figure 1A), including Dasatinib that targets KIT D816V cells (Shah et al., 2006; Gleixner et al., 2007; Arock et al., 2018). LDC 3416 was effective in selectively killing KIT D816V cells from both patient 1 and 2 iPS cells, and from KIT D816V human ES cells. Additionally, LDC 3416 had little effect on control cells without mutation (Figure 1A and B, Supplementary Figure 1F, Supplementary Table 1).

LDC 3416 is classified as a PDE3A inhibitor and surprisingly we found that conventional PDE inhibitors, such as Luteolin, Pimobendan and Cilomilast, did not impact cell viability of KIT D816V cells (Supplementary Figure 2A and B). Some PDE3A inhibitors act as molecular glues via formation of a PDE3A-SLFN12 complex to induce SLFN12 RNase activity, depletion of tRNA Leu (TAA), ribosome pausing and apoptosis (Chen et al., 2021; Garvie et al., 2021; Yan et al., 2022; Lee et al., 2023; Greulich, 2024). Thus, to extend the notion on LDC 3416 acting through PDE3A-SLFN12 complex, we subjected KIT D816V cells to a second round of hit validation by employing an additional panel of 40 compounds with PDE3, PDE4, PDE5, PDE7, PDE9 and PDE10 inhibitor activity and included kinase inhibitors as controls. All conventional PDE inhibitors were essentially inactive and Anagrelide, Zardaverine and BRD 9500, which all target the PDE3A-SLFN12 complex (Chen et al., 2021; Yan et al., 2022; Greulich, 2024; Lewis et al., 2024), were found to impair KIT D816V cell viability to a similar extent as LDC 3416 (Supplementary Figure 2B). Cytostatic compounds, like Cladribine, were not selective and effectively killed cells with and without KIT D816V mutation. Thus, we proceeded to explore the potential of LDC 3416 acting as a molecular glue through the PDE3A-SLFN12 complex.

### KIT D816V Cells Express PDE3A and SLFN12

First, we showed that KIT D816V cells express PDE3A and SLFN12 mRNA both in the presence and absence of LDC 3416 (Figure 1C). Second, KIT D816V cells also expressed PDE3A and SLFN12 at the protein level (Supplementary Figure 2C and D). Third, quantitative examination of Western blotting data indicated that LDC 3416 increased PDE3A and SLFN12 protein levels, probably through the reported protein stabilization in the PDE3A-SLFN12 complex (Supplementary Figure 2D). Isogenic iPS cell-derived progenitors without *KIT* D816V mutation (KIT control) also expressed PDE3A and SLFN12 mRNA and protein, albeit at lower levels than KIT D816V cells (Figure 1C; Supplementary Figure 2C). Thus, KIT D816V iPS cells express components of the PDE3A-SLFN12 complex and LDC 3416 left KIT D816V phosphorylation unaffected. Therefore LDC 3416 appears to act through a pathway different from inhibiting KIT D816V tyrosine kinase signaling and rather through the PDE3A-SLFN12 pathway.

We proceeded to demonstrate PDE3A and SLFN12 expression in KIT D816V mast cells derived from iPS cells using our previously generated RNA-seq data (Toledo et al., 2023; Figure 1D). KIT D816V mast cells express multiple members of the PDE and SLFN gene families (Supplementary Figure 2E). Interestingly, members of the SLFN12 cluster, such as SLFN11, SLFN12L, SLFN13 and SLFN14, but not SLFN5, were more abundantly expressed in KIT D816V mast cells compared to KIT control mast cells (Figure 1D and Supplementary Figure 2F). KIT D816V mast cells also express aryl hydrocarbon receptor interacting protein (AIP; Supplementary Figure 2E), which has been implicated in PDE3A-SLFN12 complex formation (Wu et al., 2020).

SLFN genes play a role in antiviral immune responses and some SLFN genes are upregulated by interferon α (IFNα; Al-Marsoummi et al., 2021; Jo & Pommier, 2022; Puck et al., 2015). In our previous work we found that KIT D816V mast cells display an inflammatory phenotype, including an interferon gene signature (Toledo et al., 2023). In addition, proteomic analysis of KIT D816V cells showed an increase of infection-related pathways (Rajan et al., 2023). Thus, KIT D816V might cause PDE3A and SLFN12 upregulation due to the proinflammatory phenotype of KIT D816V cells. To test this hypothesis, KIT control cells without KIT D816V mutation were treated with IFNα, and PDE3A and SLFN12 expression was found to be induced to levels similar as those in cells with KIT D816V mutation (Figure 1E). Thus, it appears that the proinflammatory environment associated by KIT D816V is consistent with elevated PDE3A and SLFN12 expression conferring sensitivity to LDC 3416.

To assess the clinical relevance of the PDE3A-SLFN12 pathway we interrogated PDE3A and SLFN12 expression in SM patient scRNA-seq data sets. In the first data set (Huang et al., 2024; GSE249445), malignant mast cells were found in peripheral blood of SM-AHN patients before TKI Avapritinib treatment and were effectively reduced after Avapritinib treatment (Figure 1F-H and Supplementary Figure 3). Importantly, malignant mast cells expressed PDE3A and SLFN12 prior to Avapritinib treatment and frequencies of PDE3A/SLFN12 expressing mast cells were effectively reduced after Avapritinib treatment (Figure 1H). In the second data set (Söderlund et al., 2023; GSE222830), BM mast cells from ISM patients enriched for KIT/CD117 and FcεRI expression were analyzed, and mast cells that were negative, single and double positive for PDE3A and SLFN12 expression were found (Supplementary Figure 4A-C) consistent with ISM representing an early, indolent stage of the disease. Interestingly, PDE3A+SLFN12+ mast cells exhibited a distinct transcriptional signature with high expression of *KIT*, *BTK* and *FCER1A*, indicative of an activated mast cell state poised to rapidly respond to degranulation stimuli (Supplementary Figure 4D and E).

To determine the impact of LDC 3416 on KIT D816V mast cells, KIT D816V iPS cells and isogenic iPS cells without *KIT* mutation were differentiated into mast cells (Supplementary Figure 1A). LDC 3416 was effective in selectively killing KIT D816V mast cells and had little effect on KIT control cells without mutation (Supplementary Figure 5A).

To further extend this observation we analyzed ROSA^KIT^ ^D816V^ cells, a human mast cell line, where *KIT* D816V was introduced by lentivirus to achieve high KIT D816V levels (Saleh et al., 2014). ROSA^KIT^ ^D816V^ cells showed high expression of PDE3A and SLFN12 mRNA and protein and acquired high sensitivity to growth inhibition by LDC 3416 (Supplementary Figure 5B-F). In parental ROSA^KIT^ ^WT^ cells without *KIT* D816V, PDE3A and SLFN12 expression was absent or very low, respectively, and cells were hardly affected by LDC 3416. The effect of LDC 3416 on further increasing PDE3A and SLFN12 protein levels was not apparent in ROSA^KIT^ ^D816V^ cells (Supplementary Figure 5C), which might be related to the already high expression levels of KIT D816V due to ectopic overexpression. Similarly, the TKI Nintedanib decreased KIT D816V phosphorylation, but not completely even at high TKI concentrations (Supplementary Figure 5D), again probably due to high KIT D816V levels.

Interestingly, the human mast cell leukemia lines HMC-1.2^KIT^ ^V560G^ ^D816V^ and HMC-1.1^KIT^ ^V560G^, containing KIT D816V mutation or not, respectively, did not express PDE3A and expressed only low levels of SLFN12 (Supplementary Figure 5B). Accordingly, HMC-1.2^KIT^ ^V560G^ ^D816V^ and HMC-1.1^KIT^ ^V560G^ cells were essentially unaffected by LDC 3416 (Supplementary Figure 5E and F). This finding underscores the importance of employing relevant cell systems for compound testing, such as patient iPS cell-derived cells, which more faithfully recapitulate the KIT D816V phenotype than established cell lines.

### PDE3A–SLFN12 Molecular Glues Target KIT D816V Cells

Given the similar activity of LDC 3416, Anagrelide and BRD 9500 in selectively killing KIT D816V cells (Supplementary Figure 2B) and their similarity in structure (Figure 2A), we determined by molecular docking the interactions of LDC 3416 within the PDE3A-SLFN12 complex and compared it to the respective interactions of PDE3A–SLFN12 molecular glues studied before (Figure 2B, Supplementary Figures 6A-D). BAY 2666605, which represents a further developed variant of BRD 9500 (Lewis et al., 2024), was also included in this study.

**Figure 2.**
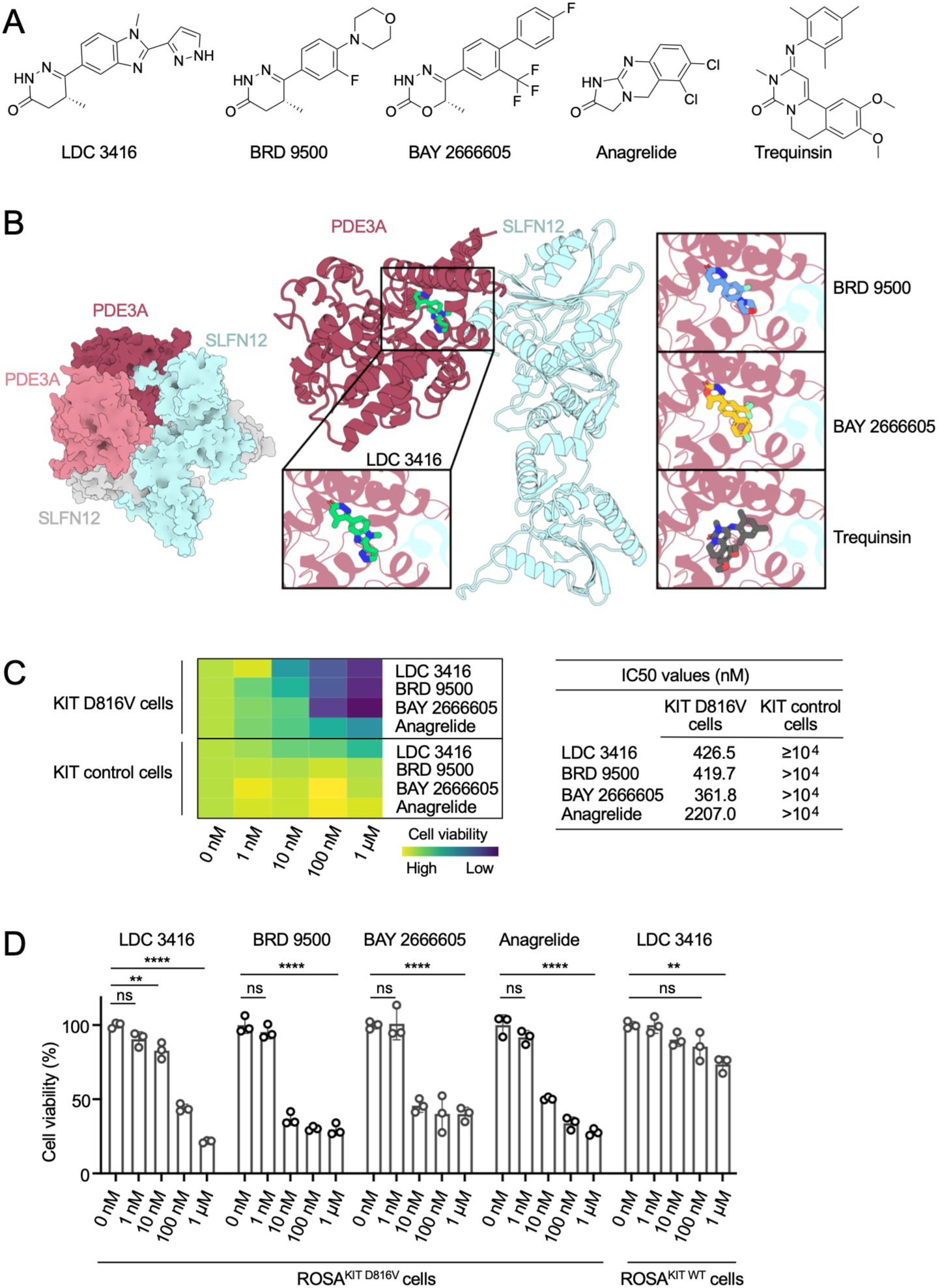
Molecular docking of LDC 3416 in PDE3A-SLFN12 complex and activity of PDE3A-SLFN12 molecular glues BRD 9500, BAY 2666605 and Anagrelide on KIT D816V cells. (A) Chemical structure of LDC 3416, BRD 9500, BAY 2666605, Anagrelide and Trequinsin. (B) Heterotetrameric PDE3A-SLFN12 assembly in surface representation (left). Ribbon representation of PDE3A-SLFN12 complex in the docking binding pose with LDC 3416 (R)-isomer (for LDC 3416 (S)-isomer see Supplementary Figure 6C and D). PDE3A is shown in raspberry cartoon derived from the X-ray crystal structure (PDB ID 7KWE). SLFN12 is depicted in cyan cartoon obtained from the cryo-EM structure (PDB ID 7LRD). Ligands are represented in licorice with a zoomed-in view of their binding poses: LDC 3416 (green), BRD 9500 (blue), BAY 2666605 (yellow), and Trequinsin (grey). (C) LDC 3416 shows a cell killing activity on KIT D816V cells similarly to the PDE3A-SLFN12 molecular glues BRD 9500, BAY 2666605 and Anagrelide (0-1 μM; n=3). Cell viability is depicted in heatmap format (yellow, high cell viability; blue, low cell viability). DMSO, vehicle control (0 nM). IC50 values of compounds on KIT D816V cells and isogenic KIT control cells without *KIT* D816V mutation are also shown. (D) LDC 3416 shows an activity profile on ROSA^KIT^ ^D816V^ cells, which is similar as the activity profile of the PDE3A-SLFN12 molecular glues BRD 9500, BAY 2666605 and Anagrelide (0-1 μM; n=3). DMSO, vehicle control (0 nM). Two-way ANOVA, * p≤0.05, ** p<0.01, *** p<0.001, and **** p<0.0001; ns, not significant.

LDC 3416 has major interactions with amino acids His961, Gln1001 and Phe1004 of the PDE3A catalytic domain, which provides a hydrophobic interface for interaction with the C-terminal alpha helix of SLFN12 (Leu553, Ile556 and Ile557) and thus PDE3A-SLFN12 complex formation, similar as the interactions of BRD 9500 and BAY 2666605. This is the same binding pocket occupied by DNMDP ligand in the cryo-EM of the PDE3A-SLFN12 complex (Garvie et al., 2021; Supplementary Figure 6D left). The PDE inhibitor Trequinsin does not maintain the same key interactions. Hydrogen bonds with His961 and Gln1001 are lost and the Trequinsin docking score indicates a lower binding affinity compared to the other compounds (Supplementary Figure 6A and B).

We then proceeded to test all three compounds, LDC 3416, BRD 9500 and BAY 2666605, in parallel on KIT D816V cells and on ROSA^KIT^ ^D816V^ cells. All compounds selectively killed KIT D816V cells and ROSA^KIT^ ^D816V^ cells and had little effect on control cells without mutation (Figure 2C and D). IC50 values of LDC 3416, BRD 9500 and BAY 2666605 were very similar. Anagrelide, a potent inhibitor of blood platelet aggregation and in clinical use for many years, with some molecular glue activity (Chen et al., 2021; Meanwell, 2023), was also active yet less potent (Figure 2C and D).

To obtain mechanistic insights into the cell killing activity of LDC 3416, we examined apoptosis as previously reported for PDE3A-SLFN12 complex formation (Li et al., 2019; Chen et al., 2021). LDC 3416 effectively induced apoptosis in KIT D816V cells and there was almost no apoptosis induced in controls without mutation (Supplementary Figure 7A and B). The same result was obtained with ROSA^KIT^ ^D816V^ cells and LDC 3416 effectively induced apoptosis in a time and concentration dependent manner (Supplementary Figure 7C and D).

SLFN12 is an RNase and molecular glue-catalyzed PDE3A binding increases SLFN12 RNase activity (Garvie et al., 2021; Yan et al., 2022). KIT D816V cells are actively proliferating and contain elevated levels of components of the ribosome assembly complex (Rajan et al., 2023) and more rRNA than isogenic controls, similar as many other cancer cells (Pelletier et al., 2018; Supplementary Figure 7E). Importantly, LDC 3416 reduced 18S rRNA levels in KIT D816V cells to levels similar in controls without mutation (Supplementary Figure 7E).

To determine the most effective PDE3A–SLFN12 molecular glue in selective killing KIT D816V cells, we performed all following studies with all three compounds LDC 3416, BRD 9500 and BAY 2666605. The conventional PDE3A inhibitor Trequinsin antagonizes molecular glues in PDE3A-SLFN12 complex formation (Wu et al., 2020; Chen et al., 2021; Garvie et al., 2021). Accordingly, Trequinsin effectively antagonized LDC 3416, BRD 9500 and BAY 2666605 in killing KIT D816V cells and ROSA^KIT^ ^D816V^ cells and had no killing effect on its own (Figure 3A; Supplementary Figure 8A and B). Trequinsin antagonism was less pronounced for ROSA^KIT^ ^D816V^ cells, probably because of the high KIT D816V levels, which made these cells particularly sensitive to drug-induced killing. Importantly, Trequinsin effectively antagonized LDC 3416 and BAY 2666605 also in killing KIT D816V mast cells (Figure 3C).

**Figure 3.**
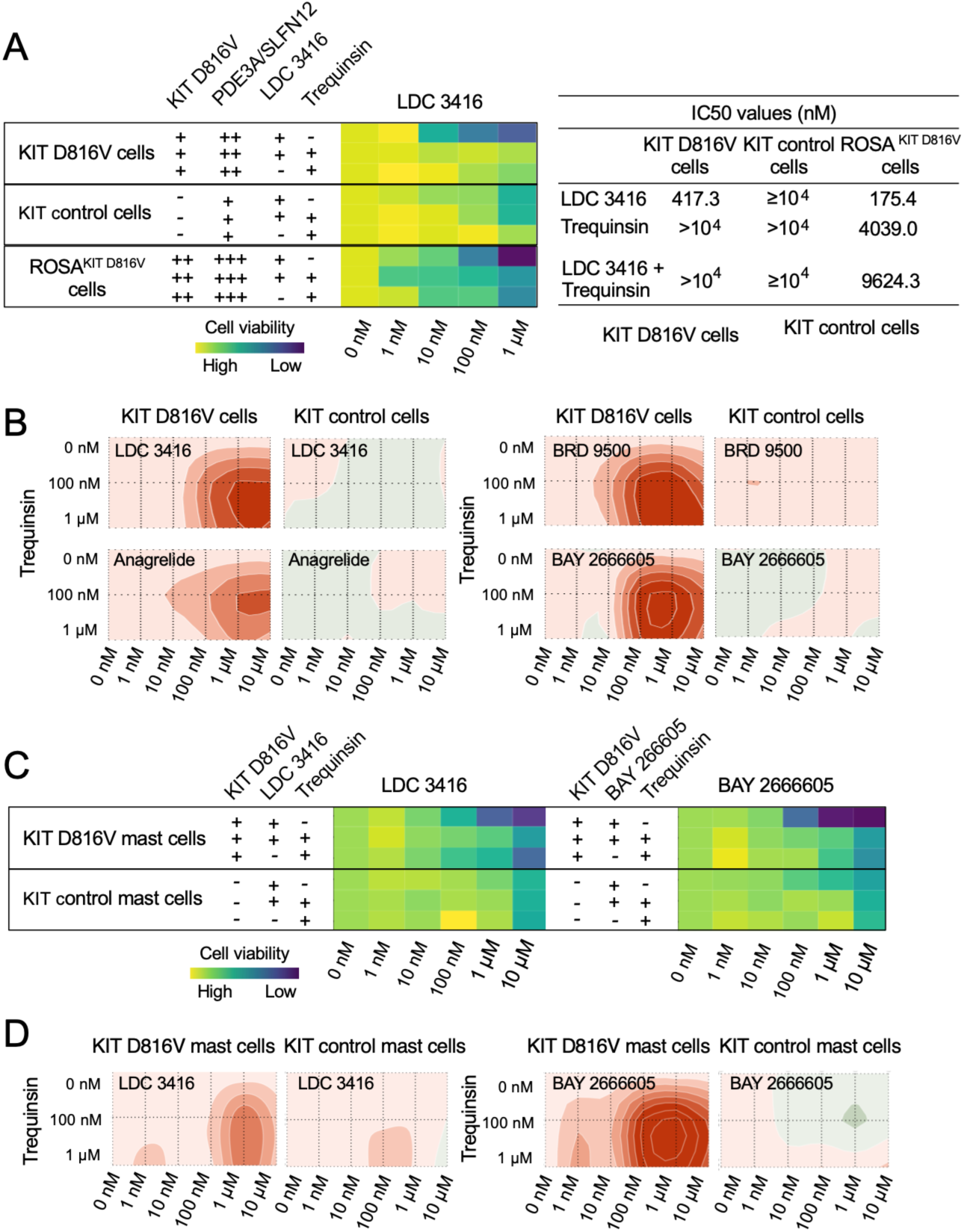
PDE3A inhibitor Trequinsin antagonizes PDE3A-SLFN12 molecular glue killing activity in KIT D816V cells. (A) PDE3A inhibitor Trequinsin antagonizes LDC 3416 killing activity in KIT D816V cells and ROSA^KIT^ ^D816V^ cells as depicted in heatmap format (yellow, high cell viability; blue, low cell viability). The *KIT* D816V mutation, levels of PDE3A-SLFN12 expression and compound treatments are indicated. Cells were treated with LDC 3416 (0-1 μM) and/or Trequinsin (1 μM) for 66 hours and cell viability was determined. DMSO, vehicle control (0 nM). IC50 values of Trequinsin antagonism on LDC 3416 in KIT D816V cells and ROSA^KIT^ ^D816V^ cells are shown. (B) Representation of Trequinsin antagonism of LDC 3416, BRD 9500, BAY 2666605, and Anagrelide in blunting killing of KIT D816V cells determined by Combenefit analysis. The red color code shows antagonism (n=3). (C) PDE3A inhibitor Trequinsin antagonizes LDC 3416 and BAY 2666605 killing activity in KIT D816V mast cells depicted as in (A). (D) Representation of Trequinsin antagonism of LDC 3416 and BAY 2666605 in KIT D816V mast cells of (C) as determined by Combenefit analysis. The red color code shows antagonism.

To further visualize our results, we employed Combenefit analysis and graphical representation of antagonism. The robust Trequinsin antagonism was evident for all three PDE3A–SLFN12 molecular glues, LDC 3416, BRD 9500 and BAY 2666605, at concentrations higher than 100 nM (Figure 3B and D, Supplementary Figure 8C). No Trequinsin antagonism was observed on KIT control cells. LDC 3416, BAY 2666605, BRD 9500 and Anagrelide differ in IC50 values (Figure 2C) and this is also seen for the Trequinsin antagonism: The stronger the PDE3A–SLFN12 molecular glue, the more effective is the competition with Trequinsin (Figure 3B and C). The weak PDE3A–SLFN12 molecular glue Anagrelide antagonized cytotoxic responses of LDC 3416, and this was less for the stronger molecular glue BRD 9500 (Supplementary Figure 8D-F).

In summary, LDC 3416 acts, similar to BRD 9500 and BAY 2666605, as molecular glue targeting the PDE3A-SLFN12 complex in KIT D816V cells to induce apoptosis, 18S rRNA degradation and finally cell death.

### PDE3A-SLFN12 Molecular Glues Combined with TKI Act in Synergy in Killing KIT D816V Cells

TKIs, such as Avapritinib and Nintedanib, target KIT D816V phosphorylation and thus KIT receptor signaling (DeAngelo et al., 2021; Gotlib et al., 2021; Reiter et al., 2022; Toledo et al., 2021). The PDE3A-SLFN12 molecular glues studied here activate SLFN12 RNase activity and thus act through a completely different pathway. Therefore, we set out to investigate whether combined treatment of TKI and PDE3A-SLFN12 molecular glues would be synergistic and therefore more effective than single treatments.

TKI Nintedanib (100 nM) in combination with 100 nM or 1 μM LDC 3416, BRD 9500 and BAY 2666605 killed KIT D816V cells more effectively than single compounds, yielding IC50 values of 6.9-39.9 nM (Figure 4A and B). Combenefit analysis revealed synergy between molecular glues and TKI. The high cytotoxicity of simultaneous molecular glue and TKI treatment was due to more efficient induction of apoptosis and KIT control cells were essentially left unaffected (Figure 4C and Supplementary Figure 9A).

**Figure 4.**
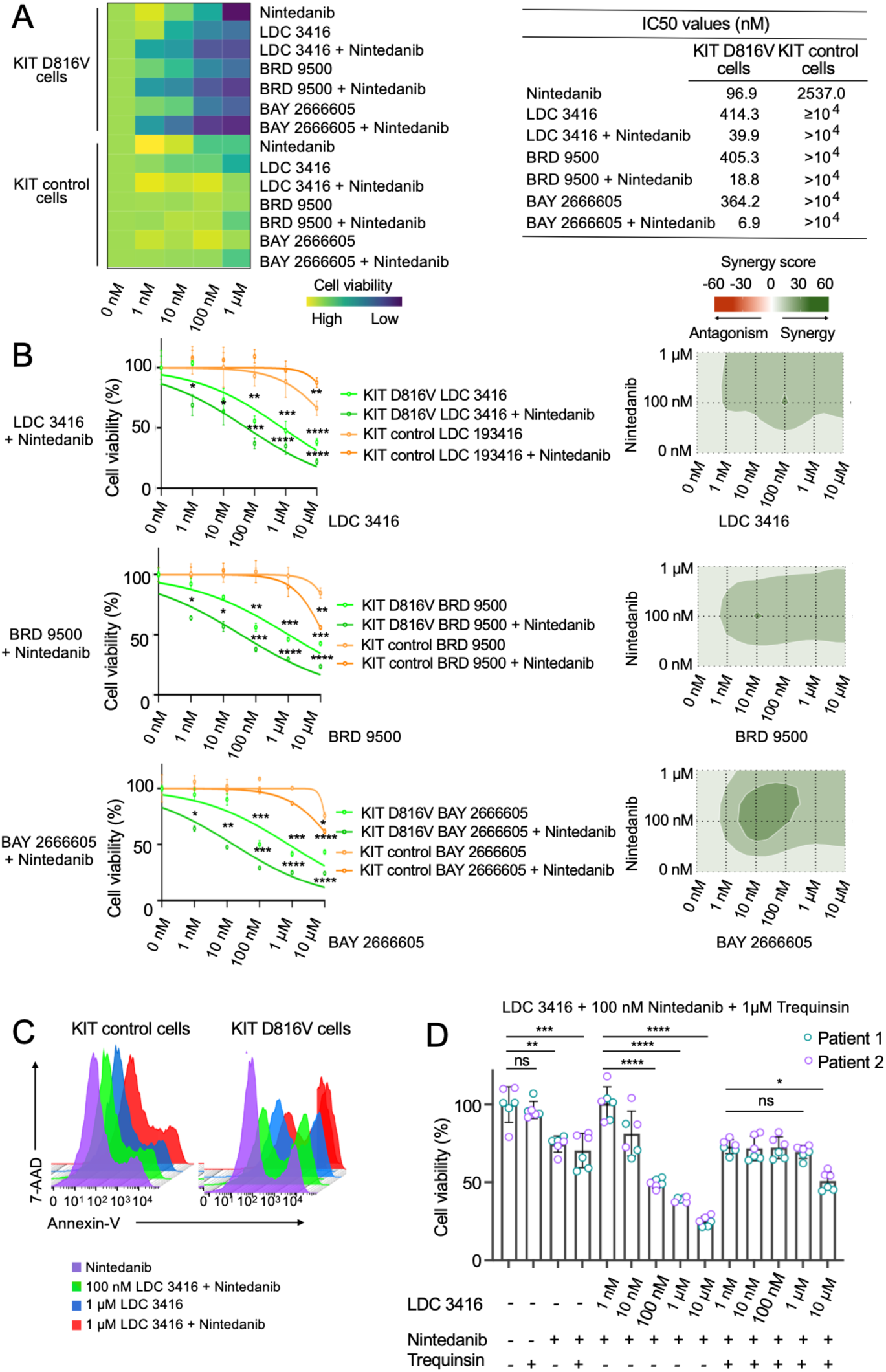
Synergy of PDE3A-SLFN12 molecular glues LDC 3416, BRD 9500 and BAY 2666605 with TKI Nintedanib in killing KIT D816V cells. (A) The PDE3A-SLFN12 molecular glues LDC 3416, BRD 9500 and BAY 2666605 synergize with TKI Nintedanib (100 nM) in killing KIT D816V cells. Growth inhibition is depicted in heatmap format (yellow, high cell viability; blue, low cell viability). IC50 values of Nintedanib synergy with molecular glues in KIT D816V cells are shown (n=3). (B) The synergy of the PDE3A-SLFN12 molecular glues LDC 3416, BRD 9500 and BAY 2666605 with TKI Nintedanib of (A) is shown in dose response curves (left) and in synergy representation by Combenefit analysis in green (right). Nintedanib concentrations were 100 nM and 1 µM, and green color code shows synergy (n=3). Data are represented as mean ± SD of triplicate wells. Two-way ANOVA, * p≤0.05, ** p<0.01, *** p<0.001, and ****p<0.0001. (C) Synergy of LDC 3416 (100 nM and 1 µM) with TKI Nintedanib (100 nM) in inducing apoptosis in KIT D816V cells assessed by Annexin V/7-AAD staining and flow cytometry (patient 1; right; n=3). Isogenic KIT control cells without mutation (left). (D) Trequinsin antagonism of LDC 3416 blocks LDC 3416 - Nintedanib synergy and rescues growth of KIT D816V cells (patient 1 and 2; n=3). Data are represented as mean ± SD of triplicate wells; Two-way ANOVA, * p≤0.05, ** p<0.01, *** p<0.001, and **** p<0.0001 for comparisons of compound treatments; ns, not significant.

A similar result was obtained for ROSA^KIT^ ^D816V^ cells (Supplementary Figure 9B-D). Co-treatment of TKI Nintedanib (100 nM) with LDC 3416, BRD 9500 and BAY 2666605 led to reduced viability and increased apoptosis, which was however rather moderate, probably due to the high KIT D816V level in these cells. Accordingly, Combenefit analysis revealed synergy only at high compound concentrations (Supplementary Figure 9B).

Finally, we performed a rescue experiment to investigate whether Trequinsin could block LDC 3416-Nintedanib synergy by inhibiting LDC 3416 (Figure 4D). KIT D816V cells were treated with increasing concentrations of LDC 3416 and 100 nM Nintedanib to show LDC 3416-Nintedanib synergy. Importantly, co-treatment with 1 µM Trequinsin effectively blocked this synergy, while retaining Nintedanib only activity. As expected, LDC 3416-Nintedanib synergy was less effective in cells without KIT D816V mutation, as was the blocking effect of Trequinsin (Supplementary Figure 9E). Essentially the same result was obtained with ROSA^KIT^ ^D816V^ cells (Supplementary Figure 9F).

We extended our analysis on PDE3A-SLFN12 molecular glue-TKI synergy to Avapritinib, since Avapritinib is approved for ISM and advSM therapy (DeAngelo et al., 2021; Gotlib et al., 2021; 2023; 2026). We demonstrated that in KIT D816V cells, the combination of LDC 3416, BRD 9500 and BAY 2666605 with TKI Avapritinib was much more effective in killing than either compound alone (Supplementary Figure 10A-D). Accordingly, Combenefit analysis identified molecular glue-TKI synergy, which was most prominent for BAY 2666605 and Avapritinib combination. KIT control cells without KIT D816V mutation were virtually unaffected. Synergy of LDC 3416, BRD 9500 and BAY 2666605 with Avapritinib was also seen in killing ROSA^KIT^ ^D816V^ cells (Supplementary Figure 10E).

We next investigated the impact of PDE3A-SLFN12 molecular glues and TKIs on primary KIT D816V MCL BM samples (Supplementary Figure 11A-C, Supplementary Table 2). BM represents a heterogeneous cell population, which contains diseased KIT D816V cells and normal healthy cells of different lineages. Thus, we set out to treat BM samples of MCL patients in a clonogenic assay. BM samples of individuals without MCL or other hematological malignancies were used as controls. Individual treatments with TKIs Avapritinib and Nintedanib or in combination with LDC 3416 or BAY 2666605 caused a reduction in colony numbers in all patient samples as well as in patient control samples (Supplementary Figure 11B).

We thus proceeded to harvest colonies and subjected them to flow cytometry analysis using lineage markers for monocyte/macrophages, granulocytes, mast cells and red cells. Patient control samples behaved similarly under all treatment conditions tested and all lineage surface markers analyzed (Supplementary Figure 11C). The impact of compounds on patient samples was heterogeneous, mirroring the patients’ disease history and therapy status. Interestingly, BM sample of MCL Patient 50 with high KIT D816V allele burden and no AHN (Supplementary Table 2) showed an effective reduction in KIT/CD117^+^ cells with both BAY 2666605 and Avapritinib (Supplementary Figure 11A and C). The result provides evidence for BAY 2666605 acting on primary MCL patient cells. PDE3A-SLFN12 molecular glue and TKI synergy was not apparent, probably because of the heterogeneity of the cell populations analyzed. This underscores the need for human-centric NAMs that yield clonal and homogeneous KIT D816V cell populations, as obtained with the iPS cell approach, to overcome this limitation.

In summary, the PDE3A-SLFN12 molecular glues LDC 3416, BRD 9500 and BAY 2666605 combined with TKI Avapritinib and Nintedanib act in synergy in killing KIT D816V cells and thus this drug combination should provide a particularly effective means for targeting malignant KIT D816V cells.

### PDE3A-SLFN12 Molecular Glues Target KIT D816V Megakaryocytes

Advanced KIT D816V SM patients, especially SM patients with associated hematological neoplasms (SM-AHN), often show elevated numbers of dysplastic megakaryocytes in BM, which is associated with myelofibrosis (Figure 5A; Sotlar et al., 2008; Chiu et al., 2009; Xu and Xu, 2018; Naumann et al., 2021; Huang et al., 2024). In these patients, platelet counts in peripheral blood can be elevated as also seen in scRNA-seq data (Figure 5B; Naumann et al., 2021; Huang et al., 2024). This is interesting, since pathognomonic skewed megakaryocytes are implicated in myelofibrosis of myeloproliferative neoplasm (MPN; Psaila et al., 2020; Flosdorf et al., 2024) and SM disease progression to SM-AHN can lead to myelofibrosis. Further to this, TKI therapy of SM with Avapritinib reduces platelet counts in peripheral blood concomitantly with reducing mast cell burden (Figure 1H and Figure 5C).

**Figure 5.**
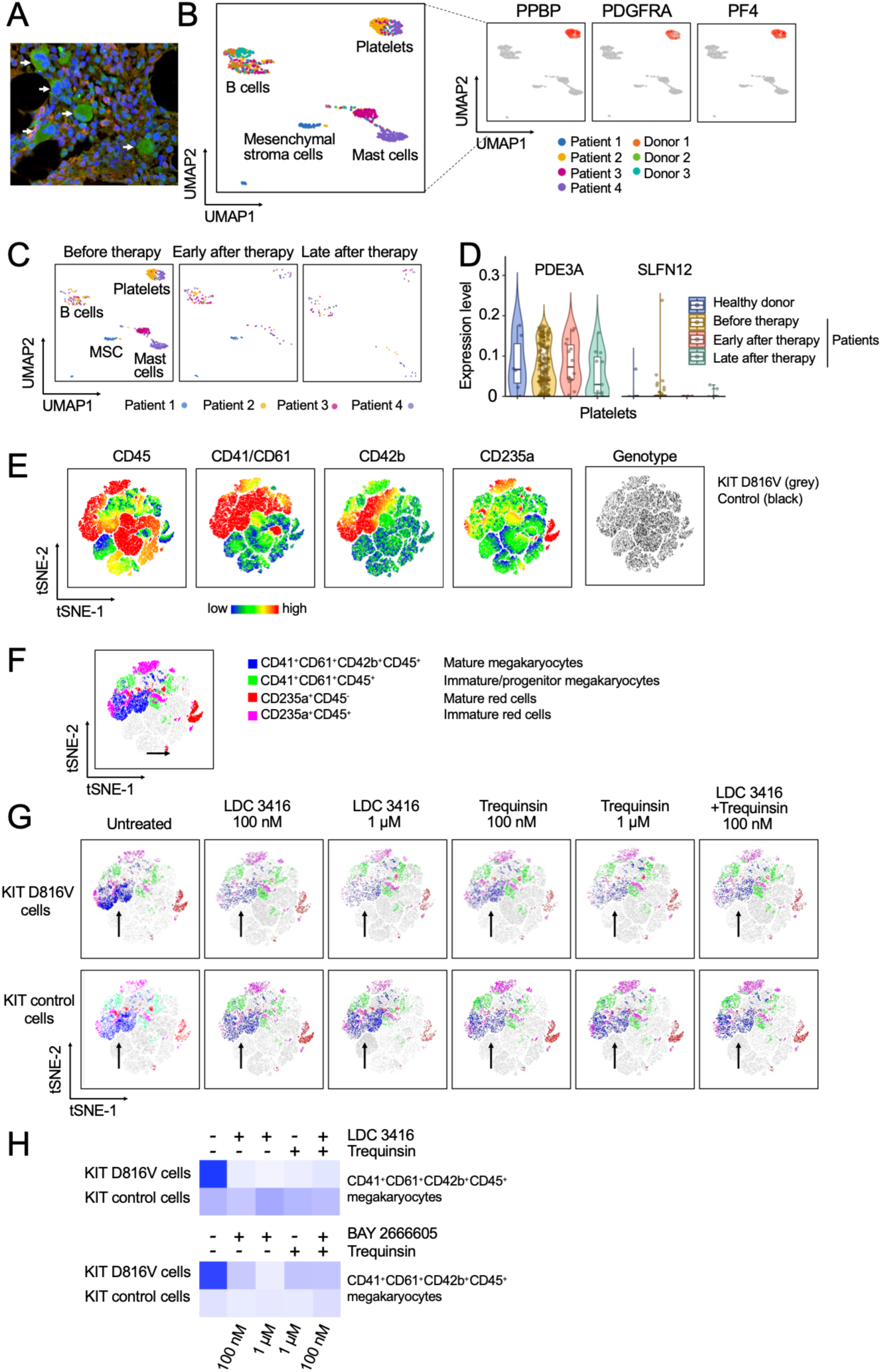
LDC 3416 targets KIT D816V megakaryocytes. (A) Megakaryocytes stained for PDE3A (green) in bone marrow of KIT D816V advSM patient (white arrows). Mast cells by tryptase staining, orange; nuclei, blue. (B) UMAP of scRNA-seq data of four KIT D816V SM-AHN patients and three healthy donors with mast cells, platelets, mesenchymal stroma cells and B cells as in Figure 1F (GSE249445). Patient samples 1-4 and healthy donor samples 1-3 are color coded as indicated. Platelet marker expression of pro-platelet basic protein (PPBP), platelet-derived growth factor receptor A (PDGFRA) and platelet factor 4 (PF4) is shown in red (right panels). (C) UMAP of platelet frequency in white blood cells of KIT D816V SM-AHN patients (Patients 1-4) before Avapritinib therapy and early (20-120 days) and late (200-400 days) after Avapritinib therapy as in (B). (D) PDE3A and SLFN12 expression in platelets in healthy donor and KIT D816V SM-AHN patients before Avapritinib therapy, early and late after Avapritinib therapy as in (C). (E) t-SNE plots of KIT D816V and KIT control cells upon megakaryocyte differentiation depicting expression of surface markers CD45, CD41/CD61, CD42b, and CD235a by flow cytometry. Color scale indicates relative expression levels (green, low and red, high). Genotype distribution is shown for KIT D816V cells and KIT control cells (grey and black, respectively; right panel). (F) t-SNE clustering and annotation of cell populations: CD41⁺CD61⁺CD42b⁺CD45⁺ mature megakaryocytes (blue), CD41⁺CD61⁺CD45⁺ immature/progenitor megakaryocytes (green), CD235a⁺CD45⁻ mature red cells (red), and CD235a⁺CD45⁺ immature erythroid cells (magenta). (G) t-SNE plots of KIT D816V cells and KIT control cells (top and bottom rows, respectively) upon treatment with LDC 3416 (100 nM, 1 µM), Trequinsin (100 nM, 1 µM) for 4-6 days, and a combination thereof (100 nM), and untreated control. Arrows indicate shifts in the mature megakaryocyte population (blue). (H) LDC 3416, BAY 2666605 and Trequinsin impact differentiation of CD41^+^CD61^+^CD42b^+^CD45^+^ megakaryocytes. Data are from panel (G) and Supplementary Figure 12D (n=1-2). Megakaryocyte frequencies are color coded (dark and light blue, high and low frequencies, respectively).

Megakaryocytes express PDE3A abundantly and SLFN12 at low levels (Figure 5A and D) and thus we investigated whether LDC 3416 and BAY 2666605 target KIT D816V megakaryocytes. KIT D816V iPS cells and isogenic control without *KIT* mutation were differentiated into KIT D816V and KIT control megakaryocytes (Flosdorf et al., 2024; Chawla et al., 2026) and subjected to compound treatment and analyzed by flow cytometry (Figure 5E-H; Supplementary Figure 12A-D). LDC 3416 and BAY 2666605 reduced frequencies of KIT D816V CD41^+^CD42b^+^CD61^+^ megakaryocytes in a dose dependent fashion and this effect was less pronounced in unmutated control (Figure 5G and H and Supplementary Figure 12D). Interestingly, Trequinsin also reduced frequencies of KIT D816V CD41^+^CD42b^+^CD61^+^ megakaryocytes, although less effectively, and did not antagonize LDC 3416 and BAY 2666605 activity. This indicates that LDC 3416 and BAY 2666605 action on KIT D816V megakaryocytes appears to occur mainly via conventional PDE3A inhibition rather than via the PDE3A-SLFN12 molecular glue pathway.

### The Impact of PDE3A-SLFN12 Molecular Glue BAY 2666605 on KIT D816V Cells *in vivo*

To explore the activity of PDE3A-SLFN12 molecular glues on KIT D816V cells *in vivo* we used two mouse models: (i) the transgenic SCL-GFP-KIT D816V model containing a humanized KIT D816V *knockin* (Pelusi et al., 2017) and (ii) the xenograft model of transplanting human ROSA^KIT^ ^D816V^ mast cells into immunodeficient NSG mice (Bibi et al., 2016; Kaiser et al., 2026; Figure 6A-F). The SCL-GFP-KIT D816V mouse model allows tamoxifen-induced expression in hematopoietic stem cells of a chimeric KIT D816V molecule containing extracellular mouse KIT sequences and intracellular human KIT sequences with the *KIT* D816V mutation (Figure 6A; Pelusi et al., 2017). Thereby the *KIT* D816V mutation is positioned within its normal environment of human KIT sequences, which is important when targeting mutant-specific tyrosine kinase activity. In the SCL-GFP-KIT D816V mouse model tamoxifen induces GFP-KIT D816V expression in hematopoietic stem cells and their progeny, and all GFP^+^ cells express KIT D816V (Figure 6A).

**Figure 6.**
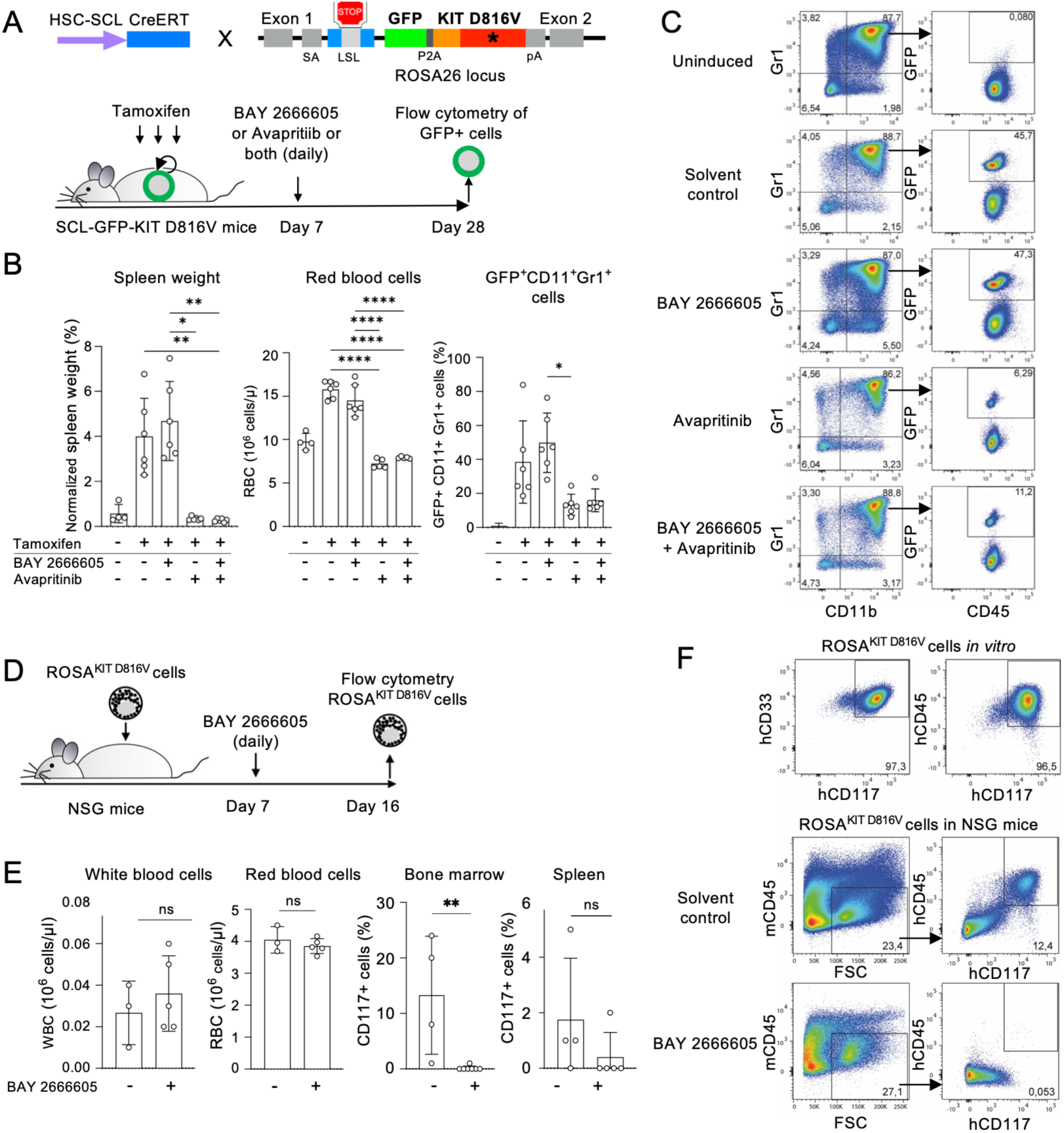
Impact of BAY 2666605 and Avapritinib on KIT D816V cells in SCL-GFP-KIT D816V mice and on ROSA^KIT^ ^D816V^ cells in NSG mice. (A) Schematic representation of HSC-SCL-CreERT GFP-KIT D816V mouse model (short SCL-GFP-KIT D816V mice). HSC-SCL-CreERT mice were bred with R26-LSL-GFP-KIT D816V mice containing a *knockin* of GFP-2A-KIT D816V in the ROSA26 locus. KIT D816V represents a chimeric KIT receptor with extracellular mouse KIT (orange) and intracellular human KIT sequences (red) with KIT D816V mutation (star). SA, splice acceptor; LSL, loxP-Stop-loxP transcriptional STOP cassette; P2A, 2A self-cleaving peptide; pA, poly A signal. Tamoxifen treatment removes the STOP cassette and induces GFP-KIT D816V expression (green). BAY 2666605 and Avapritinib treatment was from Day 7 onwards daily until Day 28 and GFP-KIT D816V^+^ cells were analyzed by flow cytometry (n=6 mice per group; for uninduced control: n=4 mice). (B) Spleen weight in relation to whole body weight per mouse and total red blood cells (RBC) in peripheral blood of SCL-GFP-KIT D816V mice at Day 28 after treatment with BAY 2666605, Avapritinib or both, or left untreated as in (A). Frequencies of myeloid GFP^+^CD11b^+^Gr1^+^ cells after treatment of SCL-GFP-KIT D816V mice analyzed by flow cytometry (right panel). (C) Representative flow cytometry dot plots of myeloid GFP^+^CD11b^+^Gr1^+^ cells in SCL-GFP-KIT D816V mice after treatment with compounds of (A) and (B) at Day 28. (D) ROSA^KIT^ ^D816V^ cells were intravenously transplanted in sublethally irradiated NSG mice. Mice were treated with BAY 2666605 from Day 7 to Day 16 (BAY 2666605, n=8 mice; solvent control, n=6 mice; one mouse of the treatment group and 3 mice of the control group were lost - most likely due to bone marrow aplasia caused by the combination of irradiation and tumor cell engraftment). ROSA^KIT^ ^D816V^ cells were analyzed by flow cytometry. (E) Total white and red blood cell counts (WBC and RBC, respectively) in peripheral blood and frequencies of hCD45^+^hCD117^+^ ROSA^KIT^ ^D816V^ cells in bone marrow and spleen at Day 16 after BAY 2666605 treatment as in (D). (F) Flow cytometry analysis of ROSA^KIT^ ^D816V^ cells in *in vitro* culture prior to transfer into NSG mice (top panel) and hCD45^+^hCD117^+^ ROSA^KIT^ ^D816V^ cells in bone marrow of NSG mice at Day 16 of treatment as in (D) (lower panel). Statistics: Brown–Forsythe and Welch one-way ANOVA followed by Dunnett T3 post-hoc tests; for non-normally distributed data: Mann–Whitney test and Kruskal–Wallis test with Dunn’s multiple comparisons: non-indicated statistics, not significant; * p <0.05; ** p <0.01; *** p <0.001; **** p <0.0001; ns, not significant.

Given that the BAY 2666605 pharmacological dynamics is well studied (Aquilanti et al., 2024; Papadopoulos et al., 2024), we focused on BAY 2666605 and applied BAY 2666605, TKI Avapritinib, or both to tamoxifen-induced SCL-GFP-KIT D816V mice (Figure 6A). Tamoxifen-induced KIT D816V expression increased spleen weight and size, red blood cell counts and GFP^+^ cells (Figure 6B and C, Supplementary Figure 13A) as expected from our previous work (Pelusi et al., 2017). Importantly, TKI Avapritinib alone or in combination with BAY 2666605 normalized spleen weight and size, and red blood cell counts to levels of uninduced mice. However, BAY 2666605 alone did not impact on spleen weight and size, red blood cell counts and GFP^+^ induced myeloid cells (Figure 6B and C).

We then continued to carefully analyze a large panel of hematopoietic cell types, including hematopoietic stem cells, erythroid, myeloid and lymphoid cells, mast cells and megakaryocytes in bone marrow and spleen (Supplementary Figure 13B-F). The effect of Avapritinib was particularly pronounced on erythroid precursor cells leading to a differentiation block and an accumulation of the CD71^+^Ter119^+^ S0 population (Supplementary Figure 13B, C and F). No impact of Avapritinib, BAY 2666605, or their combination was observed on FcεRIα^+^CD117^+^ or GFP^+^ mast cells (Supplementary Figure 13D). There was some impact of Avapritinib on GFP^+^CD150^+^CD41^+^ megakaryocytes, although not statistically significant (Supplementary Figure 13E).

The lack of BAY 2666605 activity in the SCL-GFP-KIT D816V mouse model might be due to the high specificity of PDE3A-SLFN12 molecular glues and their strict requirement for SLFN12, which has no direct orthologue in mice (Jo & Pommier, 2022). We thus proceeded to determine BAY 2666605 activity in human cells and adoptively transferred human ROSA^KIT^ ^D816V^ mast cells in NSG mice (Figure 6D). BAY 2666605 was applied daily from day 7 after ROSA^KIT^ ^D816V^ cell transplantation onwards and at day 16 differential blood counts were assessed and peripheral blood, bone marrow and spleen were analyzed by flow cytometry. BAY 2666605 had no effect on murine white and red blood cells and platelets (Figure 6E and Supplementary Figure 13G) due to the absence of SLFN12 in mice. In contrast, BAY 2666605 effectively reduced frequencies of human CD117^+^ ROSA^KIT^ ^D816V^ mast cells in bone marrow (Figure 6E and F, Supplementary Figure 13H), and to some extent also in spleen (Figure 6E).

Together, the PDE3A-SLFN12 molecular glue BAY 2666605 effectively targets human KIT D816V cells *in vivo* in mice.

## Discussion

More than 85% of patients with advanced SM carry the *KIT* D816V mutation and 75% of those patients respond clinically and molecularly to TKI Avapritinib with a reduction in *KIT* D816V variant allele frequency (VAF). Responses to Avapritinib are impressive for advanced SM and MCL (Gotlib et al., 2026) and even deep molecular responses do occur, but these are rare and in most cases do not persist, thereby resulting in a significant unmet need for these diseases.

Here we used patient KIT D816V iPS cells for drug discovery employing an open and unbiased phenotypic approach to identify non-TKI compounds that target KIT D816V cells and potentially synergize with TKIs. Screening of a library of FDA approved and experimental drugs for specific killing of KIT D816V cells identified the novel compound LDC 3416. LDC 3416 acts as a PDE3A modulator and our finding uncovered the molecular glue PDE3A-SLFN12 signaling pathway as a novel approach for the targeted elimination of malignant KIT D816V cells.

We demonstrate that cells derived from patient KIT D816V iPS cells express PDE3A-SLFN12, components of the molecular glue complex, rendering them susceptible to LDC 3416-induced killing. We extend these findings to further PDE3A-SLFN12 molecular glues, including Anagrelide, BRD 9500 and BAY 2666605. All these compounds engage in similar interactions within the tetrameric PDE3A-SLFN12 molecular glue complex as shown by molecular docking studies. Accordingly, all these compounds induce selective killing of KIT D816V cells, albeit with varying levels of activity. In various assays LDC 3416, BRD 9500 and BAY 2666605 exhibited similar activities, while Anagrelide consistently showed lower activity. Mast cell lines, such as HMC-1.1^KIT^ ^V560G^ and HMC-1.2^KIT^ ^V560G^ ^D816V^, which do not express components of the PDE3A-SLFN12 complex, proved resistant to cell-selective killing. Conversely, ROSA^KIT^ ^D816V^ cells, a human mast cell line engineered by lentivirus transduction to express high levels of KIT D816V, acquired high sensitivity to killing by PDE3A-SLFN12 molecular glues.

Furthermore, using scRNA-seq analysis, we found that mast cells from KIT D816V patients with indolent or advanced SM express PDE3A and SLFN12. This suggests the need for clinical trials to evaluate the efficacy of PDE3A-SLFN12 molecular glues in the treatment of SM. BAY 2666605 was tested in a first-in-human dose escalation study in patients with solid tumors expressing PDE3A and SLFN12 (NCT04809805). However, the study was discontinued due to thrombocytopenia before the therapeutic window could be reached (Papadopoulos et al., 2024). In cancer therapy, combinations of two or more therapeutic agents are frequently employed to enhance the activity of individual drugs or to compensate for loss of efficacy should the dosage of a specific drug need to be reduced due to adverse side effects.

In this context, we found that PDE3A-SLFN12 molecular glues act synergistically with KIT D816V selective TKIs, such as Avapritinib and Nintedanib, to kill KIT D816V cells. This is consistent with PDE3A-SLFN12 molecular glues and TKI targeting independent signaling pathways. Interestingly, the different modes of action - induction of RNAse activity vs TKI - also have conceptual consequences, and PDE3A-SLFN12 molecular glues and TKI are expected to be highly complementary when applied simultaneously. We found that high KIT D816V expression e.g. in ROSA^KIT D816V^ cells requires high TKI concentrations to effectively inhibit KIT phosphorylation but, importantly, makes these cells particularly sensitive to PDE3A-SLFN12 molecular glue action. Consequently, low KIT D816V activity might be well controlled by TKIs and high KIT D816V activity by PDE3A-SLFN12 molecular glues. Avapritinib, which is approved for SM therapy, shows an overall response rate of 75% but also leaves patients refractory to TKI treatment (DeAngelo et al., 2021; Gotlib et al., 2021). Thus, the PDE3A-SLFN12 molecular glues identified here could offer novel therapy options in cases where TKI treatment has failed or a reduction in TKI dosage is required.

In mouse models of MPN, mutated malignant cells skewed nonmutated cells in BM towards a malignant MPN phenotype, suggesting that both mutated and nonmutated cells contribute to the malignant phenotype (Kleppe et al., 2015; Bonal et al., 2025; Zenke and Koschmieder, 2025). Similarly, KIT D816V cells, which exhibit an inflammatory phenotype and an interferon gene signature (Toledo et al., 2023), can be expected to impact on cells without KIT D816V mutation. Consequently, both mutated and nonmutated cells would contribute to the malignant phenotype. Our observation that KIT D816V cells express PDEA3 and SLFN12 and that IFNα induces PDEA3 and SLFN12 in cells without KIT D816V mutation is consistent with this premise. Importantly, while TKIs will only target KIT D816V mutant cells, the molecular glues identified in this study will target both: KIT D816V mutated cells and nonmutated cells, which have been skewed under the influence of the KIT D816V mutated cells towards expression of PDEA3 and SLFN12.

Patients with advanced-stage KIT D816V SM and/or with associated hematological neoplasms (SM-AHN) may exhibit an increased number of megakaryocytes, an observation that has hitherto been underestimated in the context of mast cell malignancies. Pathognomonic skewed megakaryocytes are implicated in myelofibrosis in MPN (Psaila et al., 2020) and SM disease progression can likewise lead to myelofibrosis (Sotlar et al., 2008; Chiu et al., 2009; Xu and Xu, 2018; Naumann et al., 2021). Consequently, the PDE3A-SLFN12 molecular glues described here could provide a simultaneous targeted approach against multiple malignant KIT D816V cell types: KIT D816V stem/progenitor cells, mast cells and megakaryocytes. To substantiate this concept, we demonstrate using the KIT D816V iPS cell model that LDC 3416 and BAY 2666605 effectively inhibited megakaryocyte differentiation, although predominantly through conventional PDE3A inhibition rather than PDE3A-SLFN12 molecular glue activity.

The discovery of LDC 3416 and PDE3A-SLFN12 molecular glues targeting KIT D816V cells has implications extending beyond mast cell malignancies, affecting other diseases, such as gastrointestinal tumors (GIST) and AML. This notion aligns well with studies by Takaki et al. 2024, who identified a non-TKI compound with PDE3A-SLFN12 molecular glue properties that was effective on GIST cell lines and in GIST xenotransplantation models.

Finally, we observed that the PDE3A-SLFN12 molecular glue BAY 2666605 was effective *in vivo* in mice exclusively on human KIT D816V cells after xenotransplantation. In conventional KIT D816V mouse models, BAY 2666605 proved ineffective, even when utilizing a humanized KIT D816V *knockin* construct. This result is attributable to the strict requirement of SLFN12, which is absent in mice. The finding underscores the importance of employing human-centric NAMs for drug discovery, such as cells derived from disease and patient specific iPS cells that most faithfully recapitulate the complexity of human disease phenotypes and drug responses.

Taken together, the data reported here open the prospect for clinical use of PDE3A-SLFN12 molecular glues, either as monotherapy or in combination with TKIs, to simultaneously target multiple malignant KIT D816V cell types in mast cell malignancies and in other KIT D816V associated diseases.

## Acknowledgements

The authors thank J. Butterfield for HMC-1.1 and HMC-1.2 cell lines. We also thank all patients who kindly donated samples. We also would like to thank M. Grasshoff for imaging processing. The experiments were supported by the Interdisciplinary Center for Clinical Research (IZKF) Aachen Flow Cytometry, Genomics and Immunohistochemistry Facilities, Medical Faculty, RWTH Aachen University, Aachen, Germany. Biomaterial samples were obtained from the RWTH centralized Biomaterial Bank Aachen (RWTH cBMB, Aachen, Germany) in accordance with biomaterial bank regulations and approval of the ethics committee of the Medical Faculty, RWTH Aachen University, Aachen, Germany (EK206/09). The work was supported in parts by funds from the German Research Foundation (Deutsche Forschungsgemeinschaft; DFG) to MZ (428858617), IGC (428857561), and SK (428858786) within the Clinical Research Unit CRU344. Parts of the work were co-funded by the Ministry of Culture and Science of the State of North Rhine-Westphalia and the European Regional Development Fund (EFRE) (StemCellFactory) to BMK and MZ. GR and AP acknowledge the support by the Federal Ministry of Education and Research (BMBF) and the Federal State of North Rhine-Westphalia (NRW) as part of the NHR Program. LDC acknowledges Merck for provision of the Merck Biopharma Mini Library, which was used as a subcollection of the FDA-approved compound library employed in this study. Furthermore, LDC thanks KHAN-I Technology Transfer Fund GmbH & Co KG for funding the LDC-related project activities. XF was funded by China Scholarship Council (CSC). AK and MZ received support by Aachener Krebs- und Leukämiehilfe. AK received support by the Stiftung Universitätsmedizin Aachen. MAST was funded by CAPES-Alexander von Humboldt postdoctoral fellowship (99999.001703/2014-05). MAST and PC were funded by donation by U. Lehmann.

