## Supplementary Materials and Methods and Supplementary Figures for "PDE3A-SLFN12 Molecular Glues Target Multiple KIT D816V Cell Types in Preclinical Models of Mast Cell Malignancies"

**Culture of iPS Cells and their Differentiation into Hematopoietic Stem/Progenitor Cells, Mast Cells and Megakaryocytes**

Patient-specific iPS cells with *KIT* D816V mutation and without mutation, referred to as KIT D816V iPS cells and control iPS cells, respectively, were from ASM patient 1 and MCL patient 3 as described (Toledo et al., 2021) here referred to as patient 1 and 2, respectively. KIT D816V iPS cells captured the mutational landscape of the patients, including further mutations, such as *ASXL1, ABL1*, *NFE2*, *NRAS*, *RUNX1*, *SRSF2*, *TET2* and *TP53* (Toledo et al., 2021). Routinely, iPS cells were cultured on Matrigel (Corning, Corning, NY) in StemMACS iPS-Brew XF medium (Miltenyi Biotec, Bergisch Gladbach, Germany) as described (Flosdorf et al., 2024; Chawla et al., 2026). iPS cells were passaged with Accutase (PAN-Biotech, Aidenbach, Germany) to obtain single cells and seeded in 6-well tissue culture plates (Greiner BIO-ONE, Frickenhausen, Germany) on Matrigel with 0.1 x $10^6$ cells/well and cultured at 37°C and 5% $CO_2$. Medium was supplemented with Rho kinase (ROCK) inhibitor Y-27632 (Abcam, Cambridge, UK) or CEPT cocktail (Tristan et al., 2023; Tocris Bioscience, Bristol, UK) for the first day after passaging. Alternatively, iPS cells were cultured on mouse embryonic fibroblasts (MEF) in Knockout DMEM, 20% Knockout serum replacement, 0.1 mM non-essential amino acids, 2 mM GlutaMAX, 0.1 mM β-mercaptoethanol, 100 U/ml penicillin and 100 µg/ml streptomycin (all Thermo Fisher Scientific) and 10 ng/ml bFGF (Peprotech, Cranbury, NJ, USA). iPS cells on MEF feeder were passaged with Collagenase IV (Thermo Fisher Scientific) as described (Chawla et al., 2026).

Hematopoietic differentiation of iPS cells was induced by spin embryoid body (EB) formation and sequential addition of specific cytokines essentially as described before (Toledo et al., 2021; Flosdorf et al., 2024; Chawla et al., 2026; Supplementary Figure 1A and B). Briefly, single cell suspension of iPS cells were seeded in 96-well U-bottom suspension culture plates (Greiner BIO-ONE; 5000 cells/well) in serum free medium (SFM) with 10 ng/ml BMP-4 (Miltenyi Biotec), 10 ng/ml bFGF, 10 µM Y-27632 or CEPT cocktail, and centrifuged at 380 g for 5 minutes for EB formation and cultured at 37°C and 5% $CO_2$. SFM was a 1:1 mixture of IMDM and F12 medium (both Thermo Fisher Scientific) containing 0.5% BSA (Sigma-Aldrich), 2 mM GlutaMAX, 1% chemically defined lipid concentrate (both Thermo Fisher Scientific) and 0.4 mM 1-thioglycerol, 50 µg/ml L-ascorbic acid (L-AA) and 6 µg/ml human holo-transferrin (all Sigma-Aldrich).

Starting from day 2, EB received 50% fresh SFM medium plus cytokines every day: Day 2-8, 10 ng/ml BMP-4, 10 ng/ml bFGF, 50 ng/ml SCF and 10 ng/ml VEGF-A (both Miltenyi Biotec). Frequently, EB started to produce hematopoietic progenitors at about Day 8-10, and then BMP-4 and VEGF-A treatment was terminated and 30 ng/ml IL-3 (Miltenyi Biotec) were added from Day 8-14 to assist progenitor production.

At about Day 14, progenitors were harvested from 96-well plates and cultured for another 10-14 days in 10 cm dishes (Greiner BIO-ONE) with 1-2 x $10^6$ cells/ml in expansion medium containing 50 ng/ml SCF, Flt3 ligand (Flt3L, 1% of supernatant of Flt3L producing B16 cells), 30 ng/ml IL-3, 12.5 ng/ml IL-6/soluble IL-6 receptor (hyper-IL-6; Fischer et al., 1997), 100 U/ml penicillin and 100 µg/ml streptomycin (Thermo Fisher Scientific). Progenitors were harvested, passed through 40 µm cell strainer (Falcon) and subjected MACS selection with CD117/KIT MicroBeads (Miltenyi Biotec).

KIT D816V iPS cells were differentiated into mast cells as described (Toledo et al., 2021 and 2023). Briefly, Day 14 progenitors were harvested and cultured in SFM with 50 ng/ml SCF and 10 ng/ml hyper-IL-6 for up to 60-70 days. The frequency of mast cells was determined by flow cytometry as $CD45^+$ $CD117/KIT^{high}$ cells.

KIT D816V iPS cells were differentiated into megakaryocytes as described (Chawla et al., 2026).

Briefly, Day 10 progenitors were cultured in SFM with SCF (50 ng/ml) and TPO (20 ng/ml; both Miltenyi Biotec) for four days (Supplementary Figure 12A). On Day 14, cells were harvested and treated with either LDC 3416, BAY 2666605, Trequinsin or combinations of compounds in SFM with SCF and TPO. Day 18-21 megakaryocyte cultures were then subjected to downstream flow cytometry analysis.

Culture and differentiation of human HES-3 ES cells (ES03) and of CRISPR/Cas9 engineered KIT D816V ES cells derived thereof was performed as in Toledo et al., 2021 and essentially as described above for iPS cells.

**ROSA and HMC-1 Cell Culture**

ROSA$^{KIT\ WT}$ cells (RRID: CVCL_5G49) are a human mast cell line derived from CD34$^{+}$ cord blood cells (Saleh et al., 2014). ROSA$^{KIT\ WT}$ cells are SCF-dependent and lentivirus transduction with *KIT* D816V converted ROSA cells into a SCF-independent cell line, expressing high levels of KIT D816V, referred to as ROSA$^{KIT\ D816V}$ cells (RRID: CVCL_5G50) (Saleh et al., 2014). ROSA$^{KIT\ WT}$ and ROSA$^{KIT\ D816V}$ cells were cultured in IMDM medium (Thermo Fisher Scientific) with fetal calf serum (FCS) and SCF or without SCF, respectively, and 1 mM sodium pyruvate, MEM vitamins and amino acids, 2 mM L-glutamine, insulin, transferrin and selenium supplement, 100 U/ml penicillin and 100 µg/ml streptomycin (all Thermo Fisher Scientific) at 1-2×$10^{6}$ cells/ml as modified from Saleh et al., 2014 and Wilhelm et al., 2023.

HMC-1.2 and HMC-1.1 (RRID: CVCL_H205 and CVC_H206, respectively) cells are immortalized mast cell lines derived from a MCL patient (Butterfield et al., 1988). HMC-1.2 cells contain KIT D816V and V560G mutations, while HMC-1.1 represents the respective control cell line without KIT D816V mutation, containing the V560G mutation only. HMC-1 cells were cultured in the RPMI 1640 medium, 10% FCS, 2 mM L-glutamine, 100 U/ml penicillin and 100 µg/ml streptomycin (all Thermo Fisher Scientific; Kaiser et al., 2026). Passaging was performed every 2-3 days, and fresh culture medium was used for expansion.

**Flow Cytometry Analysis and Magnetic Activated Cell Sorting**

Cells were analyzed by flow cytometry with BD FACS Canto II or LSR Fortessa device (BD Bioscience, Franklin Lakes, NJ) as described (Toledo et al., 2021, Flosdorf et al., 2024; Chawla et al., 2026). Antibodies are listed in Supplementary Table 4. Briefly, cells were harvested, passed through a 40 µm cell strainer (Falcon) and stained with antibodies for 30 minutes at 4°C in FACS buffer (PBS, 2% FCS or 0.5% BSA, 2 mM EDTA, all Gibco). iPS cell-derived hematopoietic progenitors were cultured in RPMI 1640, 2 mM L-glutamine, 100 U/ml penicillin and 100 µg/ml streptomycin (all Thermo Fisher Scientific) for 1.5-2 hours without SCF prior to CD117/KIT staining, to achieve maximal KIT surface expression. Cells were washed with 1 ml FACS buffer to remove unbound antibodies, resuspended in 350 µl FACS buffer and subjected to flow cytometry analysis. Data analysis was with FlowJo software (BD Bioscience) and GraphPad Prism.

CD117/KIT$^{+}$ iPS cell-derived hematopoietic progenitors (referred to as KIT D816V cells and KIT control cells) were obtained by short culture without SCF (1.5-2 hours as above) followed by MACS selection with CD117/KIT MicroBeads and LS MACS columns (both Miltenyi Biotec) according to the manufacturer's instructions. KIT$^{+}$ cells were then used for compound testing, RNA and protein analysis.

**Compound Library Screening and Testing by CellTiter-Glo and MTT Assay**

A compound library of 4330 FDA-approved and experimental drugs, containing also 80 compounds of the Merck Biopharma Mini Library, was screened at 10 µM final concentration with KIT D816V hematopoietic stem/progenitor cells derived from patient iPS cells (patient 1 and 2) and KIT D816V human ES cells (referred to as KIT D816V cells). Culture was in RPMI 1640, 10% FCS, 2 mM L-glutamine, 100 U/ml penicillin and 100 µg/ml streptomycin (all Thermo Fisher Scientific) in 384- and 1536-well microtiter plates (48 hours, 37°C and 5% $CO_2$) and cell viability was assessed by CellTiter-Glo Luminescent Cell Viability Assay (Promega, Madison, WI, USA). Cells derived from isogenic iPS cells and human ES cells (HES-3) without KIT D816V mutation were used as controls (KIT control cells).

First, to ensure assay linearity and optimal signal intensity, increasing cell numbers of KIT D816V cells or KIT control cells (25, 50, 100, 200, 500 and 1000 cells/well) were seeded in 1536-well plates and evaluated in CellTiter-Glo Luminescent Cell Viability Assays (48 hours, 37°C and 5% $CO_2$, Supplementary Figure 1D). Assay quality of Z'>0.6; S/B˜10-14; S/N˜10-15 for 400 cells/well of both cell types was acceptable for screening. Second, screening of the 4330 compound library was done in 1536-well format with KIT D816V cells and KIT control cells of patient 1 (400 cells/well, 48 hours and 10 µM compound concentration). This identified 335 compounds with a bioluminescence signal of less than 50% for KIT D816V cells relative to vehicle control (Supplementary Figure 1E). Third, clustering of compounds based on chemical structure to select prototype compounds and flagging of reported artefact compounds led to a selection of 234 compounds that were further investigated in dose-response curves (Supplementary Figure 1F and Supplementary Table 1).

Hit validation and dose-response curves (0.003 to 10 µM) to determine IC50 values were then done with KIT D816V cells from patient 1 and 2 iPS cells and from KIT D816V human ES cells, and with KIT control cells of the respective isogenic iPS cells and human ES cells (HES-3) without *KIT* D816V mutation (n=2 with low day to day variation). Therefore, compounds and vehicle control DMSO were added at different concentrations by Echo Liquid Handling Technology to achieve final concentrations from 0.003 to 10 µM**.** KIT D816V cells or KIT control cells were seeded in 1536-well plates at a density of 400 cells/well in 4 µl RPMI 1640 medium, 10% FCS, 2 mM L-glutamine, 100 U/ml penicillin and 100 µg/ml streptomycin (all Thermo Fisher Scientific), cultured for 48 hours, 37°C and 5% $CO_2$ and cell viability was evaluated in CellTiter-Glo Luminescent Cell Viability Assays. Hit verification rate for KIT D816V cells was 88% and 35 compounds showed a prominent inhibition of KIT D816V cell growth vs unmutated control (Figure 1A; Supplementary Figure 1F, Supplementary Table 1). The lead compound LDC 3416 was resynthesized and found to exhibit the same activity as LDC 3416 used for library screening, hit validation and initial dose-response curves.

All other compounds were purchased from commercial chemical vendors (MedChem Express, Monmouth Junction, NJ, USA and Selleckchem, Houston, TX, USA) and dissolved as 10 mM stock solutions in DMSO (Sigma-Aldrich) in 5 µl aliquots per PCR tube for storage at -80°C and single use. Serial dilutions of compound (1 nM to 10 µM) were placed in white flat-bottom 96-well plates (Greiner BIO-ONE) in 90 µl RPMI 1640 medium supplemented with 10% FCS, 2 mM L-glutamine, 100 U/ml penicillin and 100 µg/ml streptomycin. DMSO was used as a vehicle control. Cells were added in 10 µl aliquots (20 x$10^3$ cells/well) and incubated at 37°C and 5% $CO_2$ for 66-72 hours (Toledo et al., 2021). Cell viability was determined by Cell Titer-Glo Luminescent Cell Viability Assay as above.

MTT assay was used for determining the impact of compounds on cell viability of ROSA$^{KIT\ WT}$, ROSA$^{KIT\ D816V}$, HMC-1.1$^{KIT\ V560G}$ and HMC-1.2$^{KIT\ V560G\ D816V}$ cells. Briefly, cells were cultured for 66-72 hours in flat-bottom 96-well plates (Greiner BIO-ONE; 20 x$10^3$ cells/well) in the respective culture medium with or without compounds as above. Ten µl MTT solution (Sigma-Aldrich; 5 mg/ml in PBS) were added per well and cells were incubated for another 4 hours at 37°C. To dissolve crystals, 96-well plates were centrifuged at 2000 rpm for 4 minutes, medium was removed and 100 µl DMSO/well were added, followed by incubation at room temperature for 10 minutes and the absorbance at 490 nm was recorded.

All the experiments were repeated at least three times using three-well replicates per individual experiment. Data analysis and heatmap plots were done with GraphPad Prism and Morpheus https://software.broadinstitute.org/morpheus/.

Co-treatment of compounds was performed by titration of molecular glues (LDC 3416, Anagrelide, BRD 9500 and BAY 2666605) and specific concentrations of TKIs (Nintedanib and Avapritinib) or Trequinsin. Titrations of single compounds served as controls. For example, titration of LDC 3416 plus 100 nM Nintedanib was one group, and single treatments of LDC 3416 or Nintedanib were used as controls. Data analysis to determine synergistic or antagonistic effects between compounds was performed with Combenefit (https://sourceforge.net/projects/combenefit/). Results were then processed and visualized in R with packages synergyfinder (v3.16.0), ggplot2 (v3.5.1), ggsci (v3.2.0), patchwork (v1.3.0), purrr (v1.1.0) and dplyr (v1.1.4).

**scRNA-Seq Data Analysis**

Data are from BM mast cells of ISM patients enriched for KIT/CD117 and FcεRI expression (Söderlund et al., 2023; GSE222830) and peripheral blood mononuclear cells of SM-AHN patients and healthy controls (Huang et al., 2024; GSE249445).

GSE222830 scRNA-Seq data analysis was done on preprocessed data (Söderlund et al., 2022) containing 10,914 cells and 18,500 genes in Python (v3.10.20) using Scanpy (v1.11.5). Leiden-clustered cells were annotated into six hematopoietic populations (mast cells, T cells, erythroid cells, granulocytes/monocytes, B cells/CD34+ progenitors, and basophils) based on canonical lineage markers (e.g., *KIT*, *TPSAB1*, *CPA3* for mast cells; *CD3D*, *CD3E*, *TRAC* for T cells; *HBB*, *GATA1* for erythroid cells) and visualized on the precomputed UMAP embedding (Supplementary Figure 4A and B). Cluster identity was confirmed with a marker-gene dot plot showing scaled mean expression per cluster. Statistical enrichment of *PDE3A* and *SLFN12* across these clusters was assessed with the Wilcoxon rank-sum test (one cluster vs. all others) and summarized as a dot plot in which color encoded the Wilcoxon z-score and size encoded statistical significance (−log10 adjusted p-value). Within the mast cell cluster, cells were classified by binarized expression (raw/normalized count >0) of *PDE3A* and *SLFN12*, defining double-positive (*PDE3A+SLFN12+*), *PDE3A*-only, *SLFN12*-only, and double-negative subpopulations, which were visualized on the UMAP embedding (Supplementary Figure 4C). Differential gene expression between double-positive and double-negative mast cells was tested with the Wilcoxon rank-sum test, using double-negative cells as the reference group; genes were considered significantly differentially expressed at a Benjamini-Hochberg-adjusted $p < 0.05$ and log2 fold-change >1. Results were visualized as a volcano plot (log2 fold-change vs. −log10 adjusted p-value) built in Matplotlib (v3.10.8), and as a dot plot of curated signature genes distinguishing the two subpopulations (e.g., *VAV3*, *BTK*, *KIT* enriched in *PDE3A+*/*SLFN12+* cells; *NFKB1*, *NFKBIA*, *TNFAIP3* enriched in *PDE3A-*/*SLFN12-* cells), with dot color representing log2 fold-change and dot size representing statistical significance (Supplementary Figure 4D and E). Differential expression results were processed and filtered using pandas (v2.3.3) and numpy (v2.0.2).

GSE249445 scRNA-Seq data were preprocessed with CellRanger (version 3.0.2, 10x Genomics). Genes were filtered if present in less than 3 cells using the Seurat R package (v5.2.1; Hao et al., 2024). We filtered out cells with more than 5000 genes detected and more than 15% mitochondrial transcript counts. We focused on 4 SM-AHN patients and 3 healthy controls and data were normalized with “SCTransform”. Clustering was performed with the standard Seurat pipeline, “RunPCA” on the 2000 most variable genes, followed by “FindNeighbors” on the first 30 principal components (PC) and finally using “FindClusters” with a resolution of 0.3, resulting in 18 different clusters.

For visualization, a UMAP was created using the first 30 PCs from PCA (Supplementary Figure 3A). During annotation, cluster 12 was identified as having subclusters. Cluster 12 was therefore subclustered with “FindSubCluster” and resolution 0.3 into 4 different clusters, resulting in a total of 21 clusters (Supplementary Figure 3A). Cells were subsequently annotated in relation to these clusters. Cell type annotation was performed manually by a specialist based on the most abundantly and specifically expressed genes in each cluster (Supplementary Figure 3B). For visualization of gene expression, MAGIC imputation was performed (v2.0.3.999; van Dijk et al., 2018). To better visualize and understand populations of interest, we further subset and re-clustered cells annotated as mast cells, mesenchymal stroma cells (MSC), platelets, and B cells. The procedure was as described above for the entire dataset with the exception of the resolution parameter during clustering set to 0.5.

**Gene Expression Analysis by RT-qPCR**

RNA was isolated with NucleoSpin RNA kit (Macherey-Nagel, Düren, Germany) according to the manual provided by the manufacturer. Total RNA was eluted with RNase-free water and the RNA concentration was determined by Nanodrop spectrophotometer ND-2000. RNA was reverse transcribed into cDNA using random primers and MultiScribe Reverse Transcriptase (Invitrogen). qPCR was done with gene specific primers (Eurofins Genomics, Ebersberg, Germany; Supplementary Table 3) and FAST SYBR Green Master Mix on the StepOnePlus Real-Time PCR system (both Applied Biosystems). Gene expression was normalized to GAPDH or β-actin expression and 2-dCt values were plotted as bar diagrams or in heatmap format.

**Molecular Docking**

Molecular docking studies were performed using the Schrödinger 2023-1 suite (Glide, Schrödinger, 2023). The target model was the PDE3A-SLFN12 complex, with PDE3A derived from an X-ray crystallographic structure (PDB ID 7KWE) and SLFN12 from a cryo-EM structure (PDB ID 7LRD) (both Garvie et al., 2021).

Ligands LDC 3416, BRD 9500, BAY 2666605 and Trequinsin were structurally preprocessed using the LigPrep tool, generating two stereoisomeric forms (R and S) at pH 7.2 for LDC 3416. Docking calculations were carried out using Glide (Friesner et al., 2004). The receptor grid was centered on the co-crystallized ligand.

Docking was performed on all ligands (LDC 3416, BRD 9500 and BAY 2666605) and on Trequinsin, which was used as a negative control. For LDC 3416, two stereoisomers (R and S) were evaluated. Each binding pose was ranked based on the Glide Score to assess potential binding affinities.

**Western Blotting and Immunofluorescence Analysis**

Cell pellets were collected and treated with RIPA lysis solution, NaF and $Na_3VO_4$, incubated on ice for about 1 hour, and centrifuged 10 minutes, 10,000xg at 4°C. Samples were KIT D816V cells and KIT control cells (3 x$10^6$ cells) and ROSA$^{KIT\ D816V}$ cells (5 x$10^6$ cells) treated with 100 nM Nintedanib and/or 1 µM LDC 3416 for 4 hours or left untreated (DMSO, vehicle control). Samples were subjected SDS-PAGE and blotted onto polyvinylidene difluoride (PVDF) membranes (Thermo Fisher Scientific). Membranes were blocked with 5% non-fat milk (Sigma Aldrich) in Tris-buffered saline (TBS) and incubated overnight with polyclonal rabbit anti-PDE3A

(1:200) or monoclonal rabbit anti-SLFN12 antibodies (1:500, both Abcam), polyclonal rabbit anti-KIT or polyclonal rabbit anti-p-KIT antibodies (both 1:1000, Cell Signaling Technology, Danvers, MA, USA; Supplementary Table 4). Membranes were then incubated with horse radish peroxidase (HRP) conjugated goat anti-rabbit (1:10,000, Thermo Fisher Scientific) at room temperature for 1 hour. Proteins were detected with Fusion SL device (Vilber, Marne-la-Vallee, France). β-actin protein, detected by mouse monoclonal antibody 8H10D10 (Cell Signaling Technology), was used as housekeeping control.

BM samples of SM patients with KIT D816V mutation were formalin-fixed paraffin-embedded tissue sections on coverslips. Paraffin was removed with xylene and ethanol followed by antigen retrieval in antigen unmasking solution (Vector Laboratories, Newark, CA, USA) at 90°C for 20 minutes. Samples were blocked with 5% normal goat serum and 0.25% Triton X-100 in PBS for 30 minutes at room temperature. Samples were then incubated with monoclonal mouse anti-mast cell tryptase antibody (Agilent/Dako, Carpinteria, CA) and polyclonal rabbit anti-PDE3A antibodies (Abcam; Supplementary Table 4) for detection of mast cells and PDE3A, respectively, in 1% BSA and 1% normal goat serum in PBS at 4°C overnight. Secondary antibodies were goat anti-rabbit Alexa Fluor 488 and goat anti-mouse Alexa Fluor 594 (both Invitrogen) and reaction was for 60 minutes. Nuclei were stained with DAPI and coverslips were mounted with Fluorescence Mounting Medium (Agilent/Dako). Image acquisition and processing were done with EVOS M7000 Imaging System (Invitrogen).

**Apoptosis Assay**

The apoptosis assay followed the Apoptosis Assay Kit manual (BioLegend, San Diego, CA, USA). KIT D816V progenitor cells were treated with LDC 3416 and/or Nintedanib and with DMSO vehicle control for 66 hours in 12-well plates with 0.5 x $10^6$ cells/well and put on ice. Cells were transferred to FACS tubes and resuspended in Annexin V Binding Buffer. Progenitor cells without Annexin staining were prepared for compensation. 5 µl of APC Annexin V was added to detect early apoptosis, and 5 µl of 7-AAD solution was added to detect late apoptosis. Samples were subjected to flow cytometry with BD FACS Canto II and analyzed with FlowJo software.

**CFU Assay**

Primary BM samples of MCL patients and patient controls (no MCL or other hematological malignancies; Supplementary Table 2) were subjected to Colony Forming Unit (CFU) assay with and without compounds (modified from Kalmer et al., 2025). Briefly, methylcellulose in IMDM medium (MethoCult H4230, StemCell Technologies, Vancouver, BC, Canada) was supplemented with 10 ng/ml IL-3, 14 ng/ml EPO, 10 ng/ml GM-CSF and 50 ng/ml SCF (all Immunotools, Friesoythe, Germany) and is referred to as CFU medium. Cells were seeded in CFU medium at a density of 1.5 x $10^4$ cells/ml in 24-well format with 1 ml/well in duplicates and cultured for 14 days at 37°C and 5% $CO_2$. Vehicle control DMSO, 100 nM LDC 3416 or 100 nM BAY 2666605, 100 nM Avapritinib or 100 nM Nintedanib and combinations of thereof were directly put in the CFU medium on Day 0. Colony numbers per well were determined on Day 14 by microscopy and cells were harvested and subjected to flow cytometry analysis. For flow cytometry analysis, the following panels and antibodies were used: for monocytes CD14 and CD16, for mast cells and progenitors CD45 and CD117, and for granulocytes CD66b and for erythroid cells CD235a (glycophorin) and CD71 (transferrin receptor; Supplementary Table 4).

**Animal Experiments**

All animal experiments were approved by the local authorities (Landesamt für Verbraucherschutz und Ernährung NRW, LAVE NRW, license 2024-178) and conducted in accordance with institutional guidelines. SCL-GFP-KIT D816V mice were bred in-house as previously described (Pelusi et al., 2017). Non-obese diabetic severe combined immunodeficiency gamma (NSG) mice were purchased from Charles River (Germany) (six weeks old). Detailed information on each *in vivo* experiment is depicted in the experimental scheme in Figure 6A and D.

In brief, HSC-SCL-CreERT mice were bred with R26-LSL-GFP-KIT D816V mice containing a *knockin* of GFP-2A-KIT D816V in the ROSA26 locus as previously described (Pelusi et al., 2017). KIT D816V represents a chimeric KIT receptor with extracellular mouse KIT and intracellular human KIT sequences with *KIT* D816V mutation. Tamoxifen treatment for three consecutive days removes the STOP cassette and induces GFP-KIT D816V expression. On Day 7, BAY 2666605 (20-40 mg/kg body weight), Avapritinib (30 mg/kg body weight) or their combination was administered by oral gavage until Day 28. On Day 28 mice were analyzed for the impact of treatment by assessing differential peripheral blood counts, spleen weight and size, and by flow cytometry analysis of various tissues (BM, bone marrow; PB, peripheral blood; liver; spleen).

For adoptive transfer of human $ROSA^{KIT\ D816V}$ mast cells into NSG mice, mice were sublethally irradiated with 240 cGy in MultiRad 225 Precision X-Ray device (Madison, CT, USA). Sulfamethoxazol/Trimethoprim (Ratiopharm, Ulm, Germany) was added to the drinking water in a final concentration of 100 µg/ml. Five hours post-irradiation, $5x10^6$ $ROSA^{KIT\ D816V}$ cells were transplanted intravenously as described previously (Bibi et al., 2016; Kaiser et al., 2026). From day 7 to day 16, mice were treated daily with BAY 2666605 (20-40 mg/kg body weight) or solvent control (90% PEG BioUltra 400, Sigma-Aldrich, 10% Ethanol, Merck). At terminal analysis, BM, PB, and spleen were examined by flow cytometry analysis. Long-bones were flushed with FACS buffer (PBS, 2% FCS, 2 mM EDTA, all Gibco), organs were crushed and strained (40 µm cell strainer) to get a single cell solution for antibody staining (Supplementary Table 4).

### Statistical Analysis

All data are expressed as the mean ± SD of at least three independent experiments or displayed according to specific criteria as described. Statistical analysis was performed using the Welch's t-test or Welch ANOVA using GraphPad Prism 11 software. p values ≤0.05 considered statistically significant are indicated by *; ** p <0.01; *** p <0.001; **** p <0.0001. IC50 values were calculated by analyzing nonlinear regression with GraphPad Prism 11. For the animal experiments: Brown–Forsythe and Welch one-way ANOVA followed by Dunnett T3 post-hoc tests; for non-normally distributed data: Mann–Whitney test and Kruskal–Wallis test with Dunn's multiple comparisons: not significant, all non-indicated statistics; * p <0.05; ** p <0.01; *** p <0.001; **** p <0.0001 using GraphPad Prism 11 software.

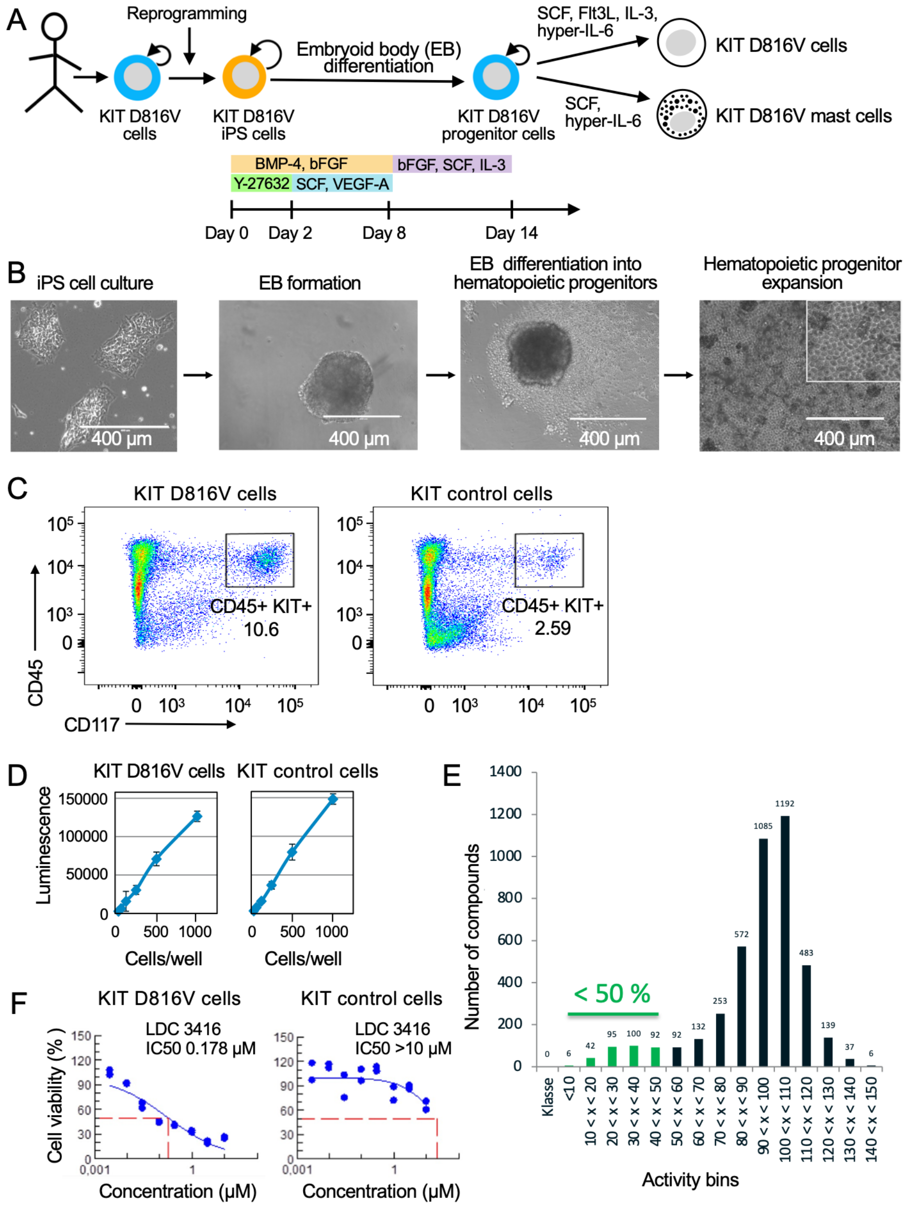


**Supplementary Figure 1. KIT D816V iPS cell differentiation into KIT D816V cells and KIT D816V mast cells and compound screening and hit validation.**

(A) Schematic representation of KIT D816V iPS cell differentiation into KIT D816V hematopoietic stem/progenitor cells (referred to as KIT D816V cells) and KIT D816V mast cells. Isogenic iPS cells without *KIT* D816V mutation were used as controls. Bent arrows indicated self-renewal. iPS cell differentiation is induced by spin embryoid body (EB) formation in 96-well format and sequential addition of compounds and specific cytokines as indicated and expansion of KIT D816V cells (SCF, Flt3L, IL-3, hyper-IL-6) and mast cells (SCF, hyper-IL-6).

(B) iPS cell culture, EB formation, EB differentiation and KIT D816V cell expansion as in (A) (scale bar, 400 µm).

(C) Representative flow cytometry analysis for CD45 and CD117/KIT expression of KIT D816V cells and KIT control cells obtained from iPS cells of patient 1. Note the higher frequency of CD117/KIT$^{+}$ cells in the KIT D816V culture compared to KIT control culture (n=3).

(D) Bioluminescence assay demonstrates linear response to cell numbers (25, 50, 100, 200, 500 and 1000 cells/well) of KIT D816V cells and KIT control cells in 1536-well format for compound screening (48 hours, 37°C and 5% $CO_2$).

(E) Screening of a library of 4330 compounds identified 335 compounds with a bioluminescence signal of less than 50% for KIT D816V cells relative to vehicle control.

(F) Representative dose response curve of LDC 3416 on KIT D816V cells and KIT control cells (1 nM to 10 µM). Cell viability is shown in percent of vehicle control (DMSO).

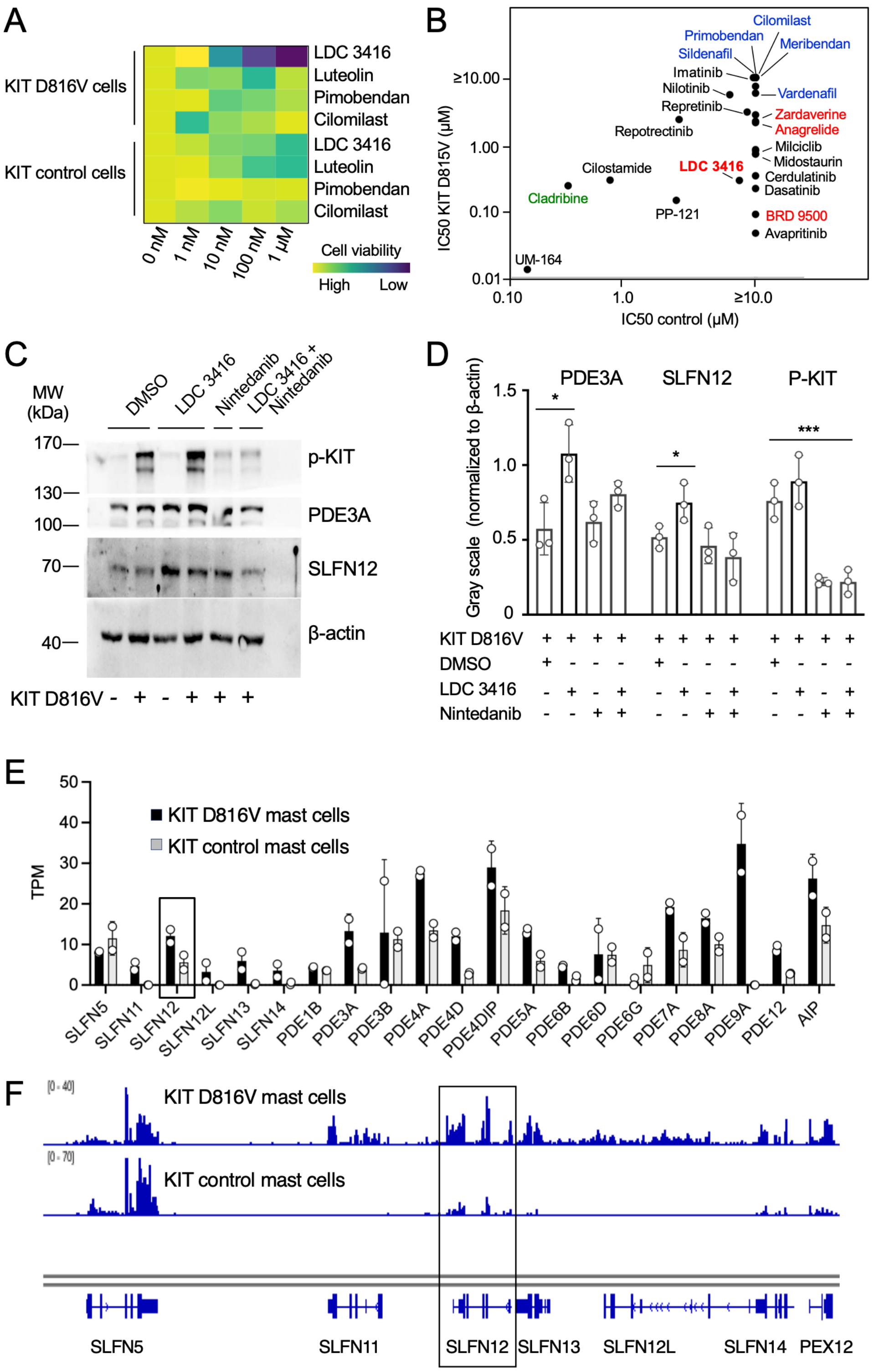


**Supplementary Figure 2. PDE compound screening and PDE3A/SLFN12 expression in KIT D816V cells.**

(A) Conventional PDE inhibitors Luteolin, Primobendan and Cilomilast do not impact cell viability of KIT D816V cells (0-1 µM; n=3). LDC 3416 activity on the respective KIT D816V cells is shown as control. Cell viability is depicted in heatmap format (yellow, high cell viability; blue, low cell viability). DMSO, vehicle control.

(B) KIT D816V cells and KIT control cells (patient 1) were subjected to screening with PDE3, PDE4, PDE5, PDE7, PDE9 and PDE10 inhibitors (40 compounds) and kinase inhibitors (15 compounds) as controls. IC50 values upon drug treatment are shown and IC50 values equal to or above 10 µM were not resolved (≥10.0). Most PDE inhibitors had IC50 values ≥10 and representative PDE inhibitors are indicated in blue. PDE inhibitors targeting the PDE3A-SLFN12 complex and LDC 3416 are in red. Representative TKI are shown in black. Cladribine, chemotherapy drug, green.

(C) PDE3A and SLFN12 protein expression in KIT D816V cells and KIT control cells (KIT D816V + and -, respectively) by Western blotting (n=3). A representative image is shown and positions of molecular weight (MW) markers are indicated. p-KIT, phosphorylated KIT; β-actin, loading control.

(D) PDE3A and SLFN12 protein expression and KIT phosphorylation (p-KIT) by Western blotting and quantification of gray scale analysis in KIT D816V cells (n=3). DMSO, vehicle control. LDC 3416 and Nintedanib treatments are indicated (+). Impact of LDC 3416 treatment (1 μM, 4 hours) on PDE3A and SLFN12 protein expression (two-way ANOVA, * $p=0.02$ and * $p=0.04$, respectively) and inhibition of p-KIT by Nintedanib (100 nM, 4 hours, *** $p<0.001$) are shown. The data are represented as the mean ± SD of 3 independent experiments.

(E) Expression of SLFN and PDE1-12 family members and of aryl hydrocarbon receptor interacting protein (AIP) in KIT D816V mast cells and isogenic KIT control cells without mutation (black and open bars, respectively; n=2). Data are from our RNA-seq data (GSE223883, Toledo et al., 2023). SLFN12 expression, boxed; TPM, transcripts per million reads.

(F) RNA-seq profile of human SLFN5-SLFN14 gene cluster of KIT D816V mast cells and isogenic control of GSE223883 in (E). SLFN12 expression, boxed.

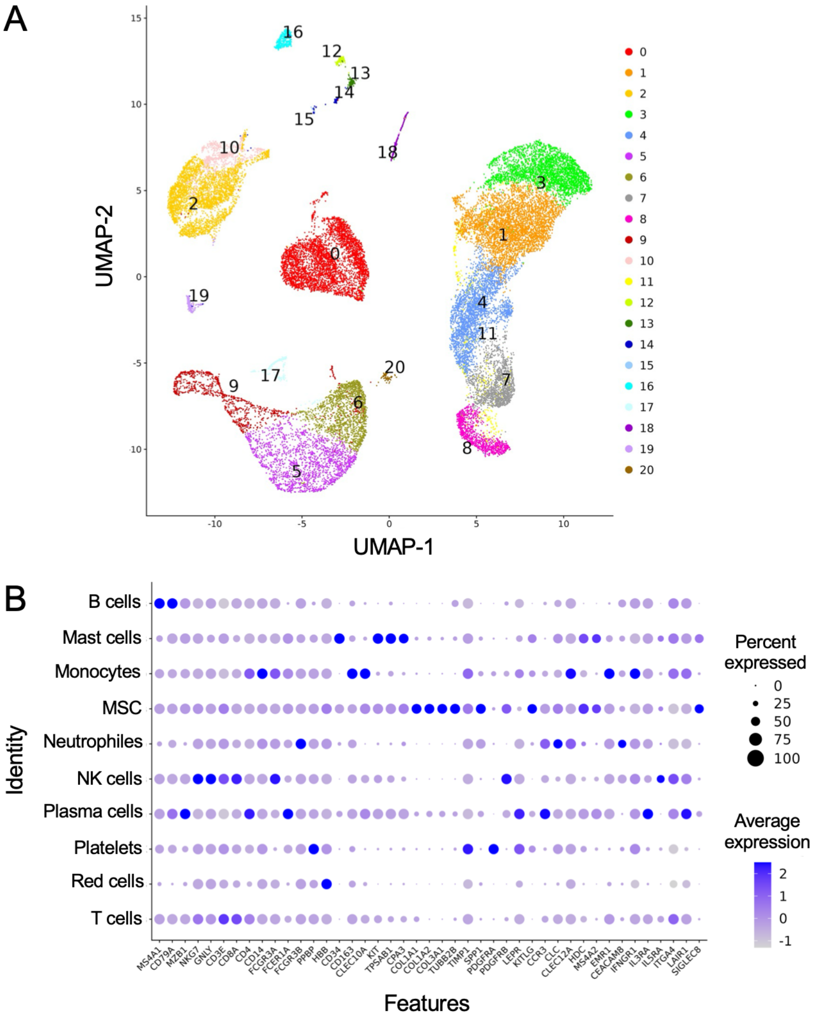


**Supplementary Figure 3. UMAP visualization of SM-AHN patient scRNA-seq data.**

(A) UMAP visualization of clustering analysis of scRNA-seq data of human SM-AHN patients and healthy controls described by Huang et al., 2024 (GSE249445) resulting in 21 different clusters.

(B) Dot plot depicting gene expression across relevant genes for individual cell types. Cell type annotation was by a manual and iterative process based on gene expression data of each cluster. The final annotation consisted of 10 different cell types (see Figure 1F).

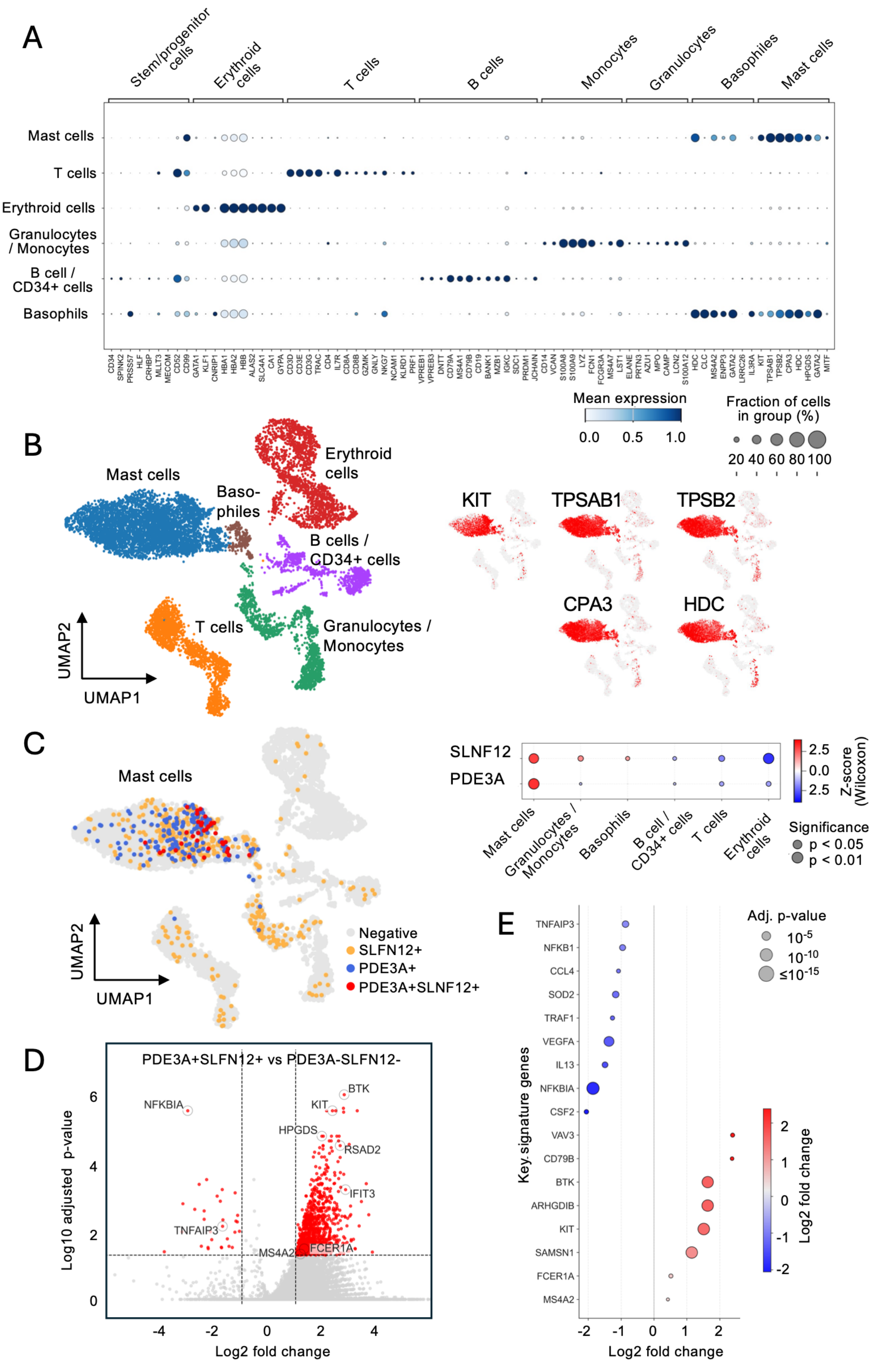

**Supplementary Figure 4. PDE3A and SLFN12 expression in KIT/CD117⁺ FcεR⁺ mast cells of ISM patients by scRNA-seq analysis.**

(A) scRNA-seq data of KIT/CD117⁺ FcεRI⁺-enriched BM mast cells from 3 ISM patients (Söderlund et al., 2022; GSE222830) were subjected to marker gene expression analysis for cell type annotation by reference database queries (CellMarker and PanglaoDB) coupled with iterative, manual curation of cluster specific expression profiles.

(B) UMAP visualization of scRNA-seq data of KIT/CD117⁺ FcεRI⁺ enriched BM mast cells of (A) and of mast cell marker expression *CD117/KIT*, mast cell tryptase (*TPSAB1*), mast cell tryptase beta II (*TPSB2*), carboxypeptidase A3 (*CPA3*) and histidine decarboxylase (*HDC*) in red.

(C) *PDE3A* and *SLFN12* gene expression in mast cells of (B). *PDE3A* and *SLFN12* double-positive cells (*PDE3A+SLFN12+*), *PDE3A*-only, *SLFN12*-only, and double-negative cells (*PDE3A+, SLFN12+*- and Negative, respectively) are color coded and shown.

(D) Volcano plot showing differential gene expression between *PDE3A+SLFN12+* and *PDE3A- SLFN12-* mast cell subpopulations of (C).

(E) DGE analysis of *PDE3A+SLFN12+* versus *PDE3A-SLFN12-* mast cells, demonstrating upregulation of *KIT*, *BTK*, and *FCERA1* and downregulation of the NF-kB inhibitor alpha (*NFKBIA*) expression.

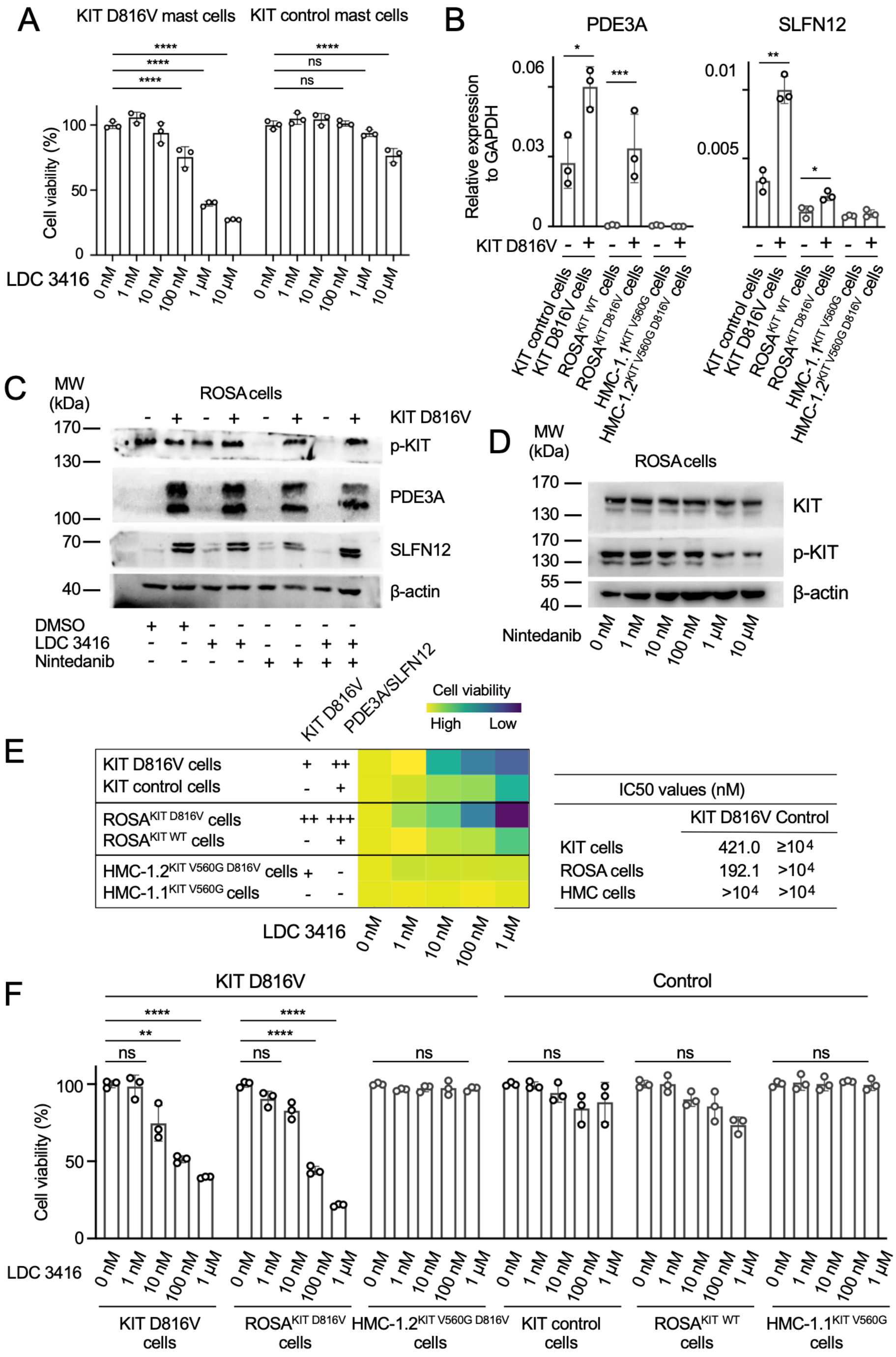


**Supplementary Figure 5. LDC 3416 kills KIT D816V mast cells and PDE3A/SLFN12 expression in KIT D816V cells and ROSA$^{KIT\ D816V}$ cells and response to LDC 3416.**

(A) Drug response curves for LDC 3416 on KIT D816V mast cells of patient 1 (n=3-5). KIT control cells, isogenic cells of patient 1 without *KIT* D816V mutation. DMSO, vehicle control (0 nM). Data are represented as mean ± SD of triplicates. Two-way ANOVA, **** p<0.0001 in comparison to no compound treatment (0 nM); ns, not significant.

(B) KIT D816V cells from iPS cells and ROSA$^{KIT\ D816V}$ cells express more PDE3A and SLFN12 mRNA by quantitative RT-PCR compared to KIT control cells without *KIT* D816V mutation and ROSA$^{KIT\ WT}$ cells without KIT D816V overexpression (KIT D816V + and -, respectively). HMC-1.1$^{KIT\ V560G}$ and HMC1.2$^{KIT\ V560G\ D816V}$ cell lines show no or low expression of PDE3A and SLFN12 mRNA, respectively (n=3). Welch's t-test, KIT D816V cells from iPS cells, PDE3A, * p=0.017 and SLFN12, ** p=0.001; ROSA cells, PDE3A, *** p<0.001 and SLFN12, * p=0.02.

(C) ROSA$^{KIT\ D816V}$ cells and ROSA$^{KIT\ WT}$ cells (KIT D816V + and -, respectively) were analysed for PDE3A, SLFN12 and phosphorylated KIT (p-KIT) protein expression in response to LDC 3416 (1 µM, 4 hours) and/or 100 nM Nintedanib (100 nM, 4 hours) by Western blotting (n=3). DMSO, vehicle control. Positions of molecular weight markers are indicated.

(D) Inhibition of p-KIT in ROSA$^{\text{KIT D816V}}$ cells upon Nintedanib treatment (1 nM to 10 µM) by Western blotting as in (C).

(E) LDC 3416 impacts on viability of KIT D816V cells and ROSA$^{\text{KIT D816V}}$ cells. KIT D816V and PDE3A/SLFN12 expression are indicated. Data are shown in heatmap format (yellow, high cell viability; blue, low cell viability). Controls, isogenic KIT control cells and ROSA$^{\text{KIT WT}}$ cells. HMC-1 cells not expressing PDE3A/SLFN12 are unaffected by LDC 3416 treatment (n=6, 3 independent experiments). DMSO, vehicle control (0 nM). IC50 values of LDC 3416 on KIT D816V cells, ROSA$^{\text{KIT D816V}}$ cells and the respective controls without *KIT* D816V mutation and of HMC1.2$^{\text{KIT V560G D816V}}$ and HMC-1.1$^{\text{KIT V560G}}$ cells are shown.

(F) Quantification of LDC 3416 activity on cell viability of KIT D816V cells, ROSA$^{\text{KIT D816V}}$ cells and HMC-1.2$^{\text{KIT V560G D816V}}$ cells and on the respective cells without mutation (KIT control cells, ROSA$^{\text{KIT WT}}$ cells and HMC1-1$^{\text{KIT V560G}}$, respectively). HMC-1.2$^{\text{KIT V560G D816V}}$ and HMC1-1$^{\text{KIT V560G}}$ cells not expressing PDE3A/SLFN12 are unaffected by LDC 3416 treatment (n=6, 3 independent experiments). Two-way ANOVA, KIT D816V cells, 100 nM LDC 3416, ** p=0.009; 1 µM LDC 3416, *** p=0.0002; ROSA$^{\text{KIT D816V}}$ cells, 100 nM LDC 3416, *** p=0.0004) and 1 µM LDC 3416 (**** p<0.0001). ns, not significant.

A

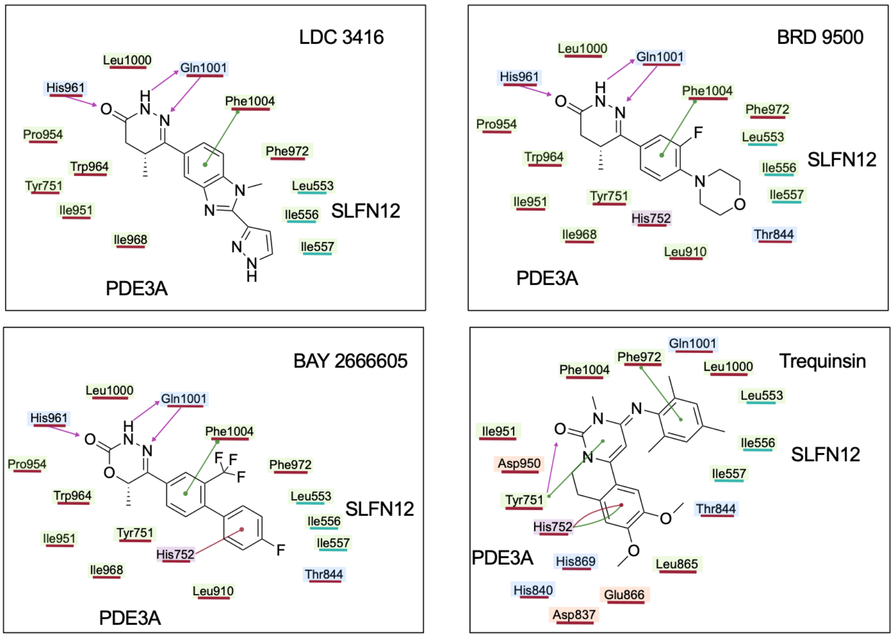


B

| Compound | Docking score (kcal/mol) |
|---|---|
| LDC 3416 | -10.6 |
| BRD 9500 | -11.3 |
| BAY 2666605 | -11.5 |
| Trequinsin | -7.5 |

C

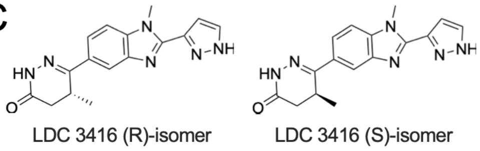


D

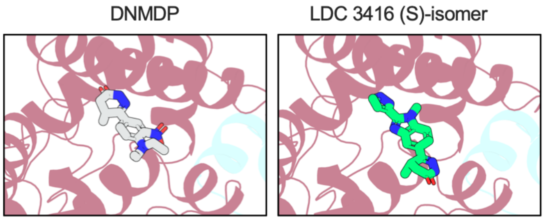


**Supplementary Figure 6. Molecular docking of LDC 3416, BRD 9500, BAY 2666605 and Trequinsin in PDE3A-SLFN12 complex.**

(A) Protein-ligand interaction pattern for LDC 3416 (R)-isomer, BRD 9500, BAY 2666605 and Trequinsin within the PDE3A-SLFN12 complex are shown in 2D representation. Amino acid residues in PDE3A and SLFN12 within 3 Å from the ligand in the binding pocket are shown with residue name and number and are labeled for PDE3A and SLFN12 (magenta bar and turquoise bar, respectively). The residue outlines are color-coded based on the interaction type: light blue for polar interactions, green for hydrophobic interactions, violet for positively charged residues and orange for negatively charged residues. Green lines indicate π-π stacking interactions, red lines represent π-cation interactions and magenta arrows represent hydrogen bonds.

(B) Glide docking scores for the best binding pose of LDC 3416 (R)-isomer, BRD 9500, BAY 2666605 and Trequinsin in PDE3A-SLFN12 complex.

(C) Chemical structure of LDC 3416 enantiomers: LDC 3416 (R)-isomer and LDC 3416 (S)-isomer, respectively.

(D) Ribbon representation of the PDE3A-SLFN12 complex model with a zoomed-in view of the binding site with co-crystalized ligand DNMDP (in licorice, grey) from PDB ID 7KWE (left). Ribbon representation of the PDE3A-SLFN12 complex model with a zoomed-in view of the binding pose with the LDC 3416 (S)-isomer (in licorice, green, right). In the docking binding pose LDC 3416 (S)-isomer adopts an orientation that is flipped upside down compared to DNMDP in the structure binding pose. Specifically, the positioning of key functional groups is inverted, leading to a distinct interaction pattern with the binding site.

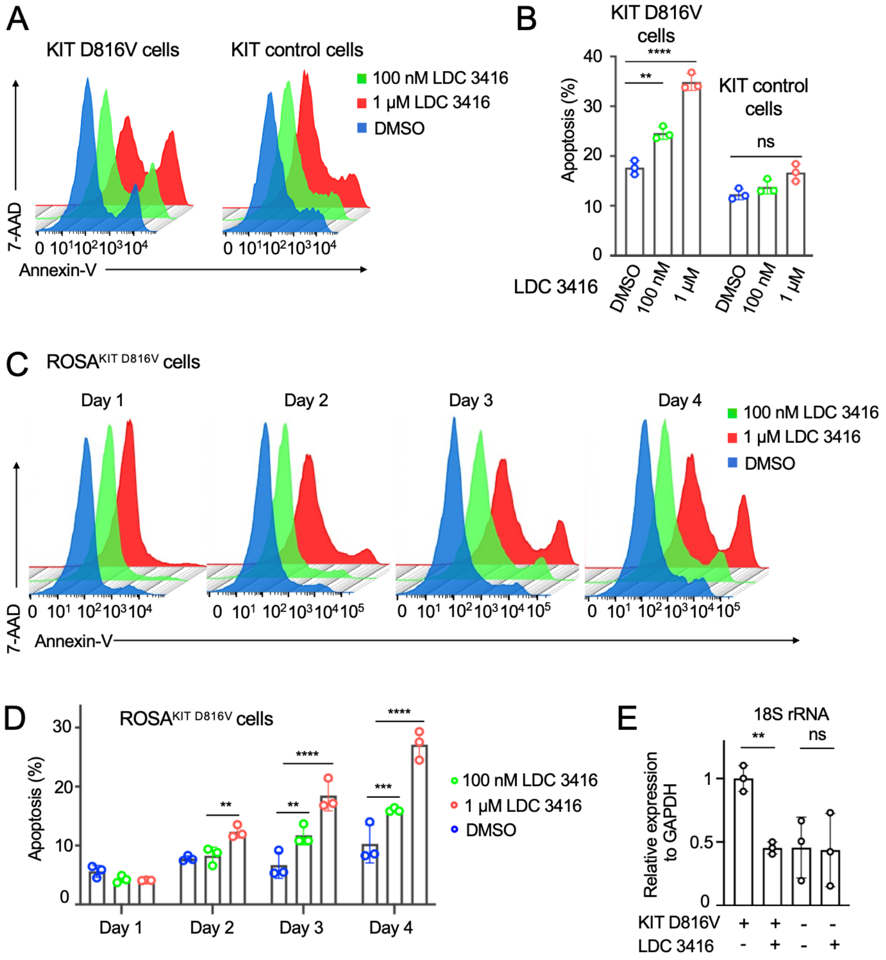


**Supplementary Figure 7. LDC 3416 induces apoptosis in KIT D816V cells and ROSA$^{KIT\ D816V}$ cells.**

(A) LDC 3416 induces apoptosis in KIT D816V cells measured by Annexin V/7-AAD staining and flow cytometry (patient 1; n=3). KIT control cells, isogenic control without mutation. DMSO, vehicle control.

(B) Quantification of apoptosis analysis of (A). Data are in percent of total cells. Two-way ANOVA, 100 nM LDC 3416, ** p=0.003 and 1 µM LDC 3416, **** p<0.0001; ns, not significant.

(C) LDC 3416 induces apoptosis in ROSA$^{KIT\ D816V}$ cells measured by Annexin V/7-AAD staining and flow cytometry as in (A) (n=3). DMSO, vehicle control.

(D) Kinetics of LDC 3416 treatment and analysis for apoptosis as in (C). Data are in percent of total cell numbers. Two-way ANOVA, ** p<0.01, *** p<0.001, and **** p<0.0001.

(E) LDC 3416 treatment (4 hours) reduces 18S rRNA expression in KIT D816V cells (KIT D816V+) by quantitative RT-PCR. 18S rRNA levels were unaffected by LDC 3416 treatment in KIT control cells without mutation (KIT D816V -) (n=3). Welch's t-test, ** p=0.0008, ns, not significant.

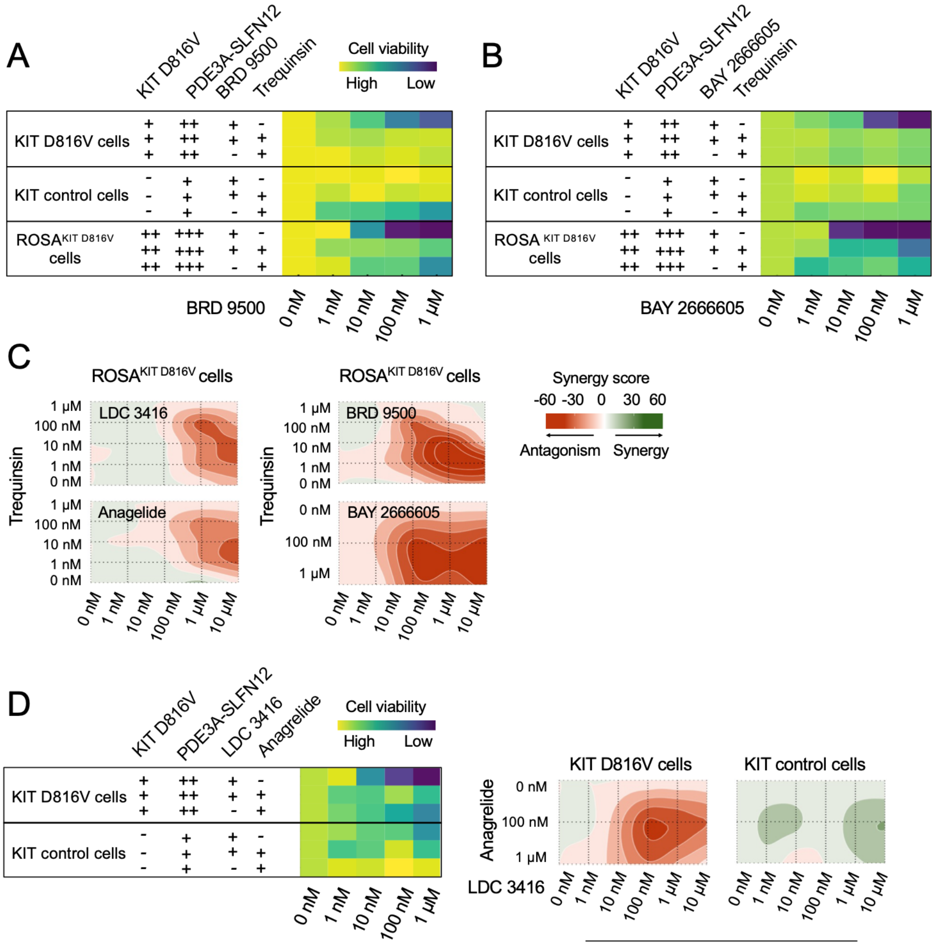


| IC50 values (nM) | KIT D816V cells | KIT control cells |
|---|---|---|
| LDC 3416 | 438.2 | ≥10^4 |
| Anagrelide | 2163.2 | >10^4 |
| LDC 3416 + Anagrelide | >10^4 | >10^4 |
| BRD 9500 | 408.5 | >10^4 |
| LDC 3416 + BRD 9500 | 6776.0 | >10^4 |

**Supplementary Figure 8. PDE3A inhibitor Trequinsin antagonizes molecular glue BRD 9500 and BAY 2666605 killing activity in KIT D816V cells.**

(A and B) PDE3A inhibitor Trequinsin antagonizes molecular glue BRD 9500 and BAY 2666605 activity in KIT D816V cells and ROSA$^{KIT\ D816V}$ cells (yellow, high cell viability; blue, low cell viability). KIT D816V and PDE3A-SLFN12 expression and compound treatments are indicated. Cells were treated with BRD 9500 and BAY 2666605 (0-1µM), respectively, and/or Trequinsin (1µM) for 66 hours and cell viability was determined. DMSO, vehicle control (0 nM).

(C) Representation of Trequinsin antagonism of the molecular glues LDC 3416, BRD 9500, BAY 2666605, and Anagrelide in ROSA$^{KIT\ D816V}$ cells determined by Combenefit analysis. Concentrations of Trequinsin, LDC 3416, BRD 9500 and Anagrelide were 1 nM - 1 µM. For BAY 2666605 / Trequinsin co-treatment concentrations of Trequinsin were 1 nM -1 µM, and BAY 2666605 concentrations were 100 nM and 1 µM. The red color code shows antagonism (n=3).

(D) Anagrelide antagonizes LDC 3416 activity in KIT D816V cells but not in KIT control cells without *KIT* D816V mutation. Data are shown in heatmap format as in (A) and in antagonism representation as in (C) (n=3).

(E) BRD 9500 compound antagonizes LDC 3416 activity in killing KIT D816V cells, which however was less prominent than the antagonism by Trequinsin (Figure 3B) and Angelide (panel D). Data are in heatmap format as in (A) (n=3).

(F) IC50 values of Anagrelide and BRD 9500 antagonizes LDC 3416 activity in KIT D816V cells of panels (D) and (E).

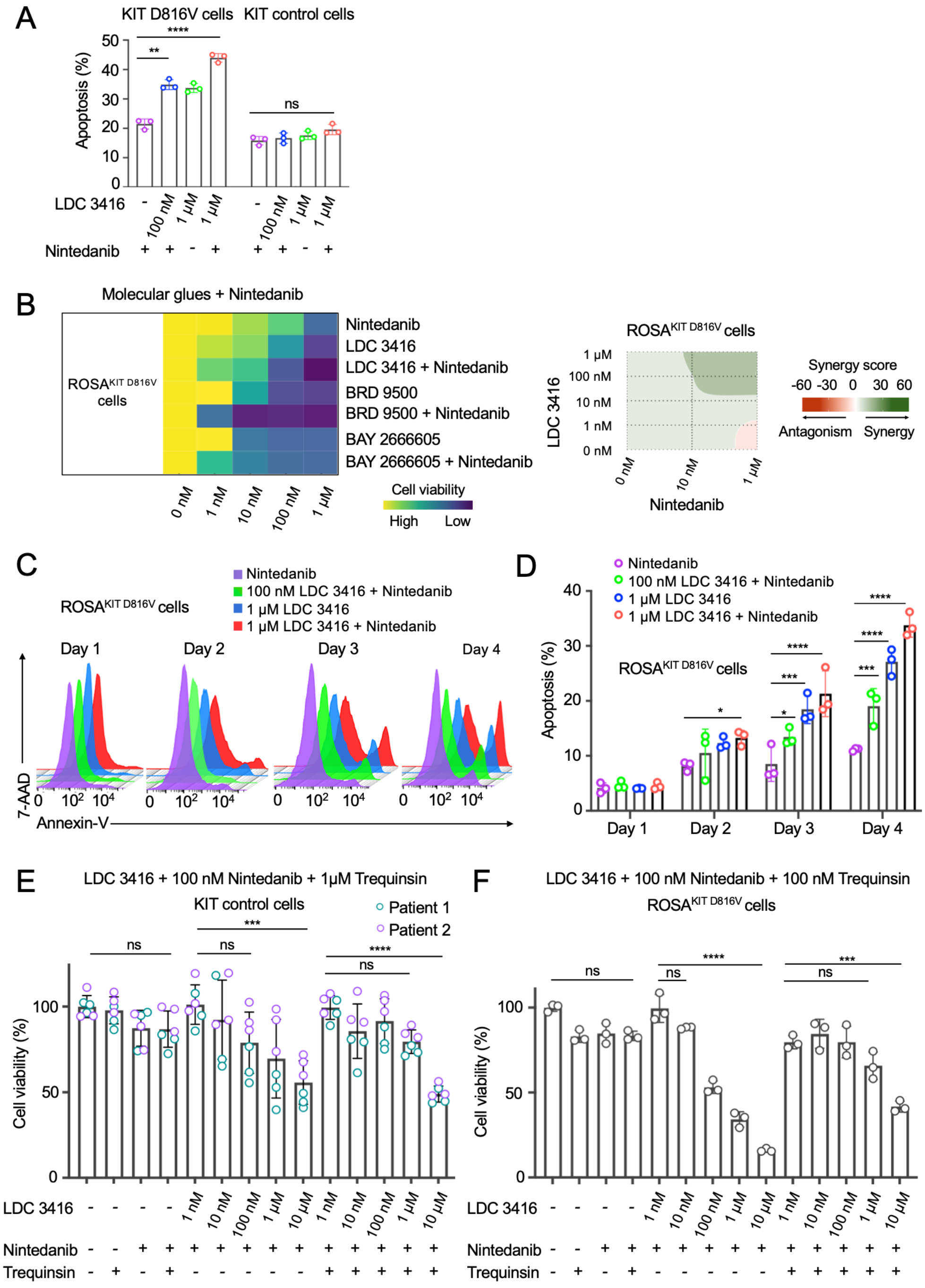


**Supplementary Figure 9. Synergy of molecular glues LDC 3416, BRD 9500 and BAY 2666605 with Nintedanib in killing and apoptosis of ROSA$^{KIT\ D816V}$ cells.**

(A) Quantification of synergy of LDC 3416 (100 nM and 1 µM) with TKI Nintedanib (100 nM) in inducing apoptosis in KIT D816V cells of Figure 4C (patient 1; n=3). Control, KIT control cells without mutation. Two-way ANOVA, ** p=0.0027, **** p<0.0001; ns, not significant.

(B) Co-treatment of LDC 3416, BRD 9500 and BAY 2666605 with TKI Nintedanib (100 nM) on viability of ROSA$^{KIT\ D816V}$ cells. Cell viability by MTT assay is depicted in heatmap format (yellow, high cell viability; blue, low cell viability; n=3). Synergy representation by Combenefit analysis of LDC 3416 and Nintedanib. Green color code shows synergy.

(C and D) Co-treatment of LDC 3416 (100 nM and 1 µM) and Nintedanib (100 nM) in induction of apoptosis of ROSA$^{KIT\ D816V}$ cells. (C) Representative Annexin V/7-AAD staining and analysis by flow cytometry at day 1 to 4. (D) Quantification of data of (E) in percent of total cells (n=3). Two-way ANOVA, * p≤0.05, *** p<0.001, and **** p<0.0001.

(E) KIT control cells of Trequinsin antagonism (1 µM) on LDC 3416 (1 nM to 10 µM) of Figure 4D. (n=3). Two-way ANOVA, *** p<0.001, **** p<0.0001; ns, not significant.

(F) Trequinsin (100 nM) antagonizes LDC 3416 synergy with Nintedanib (100 nM) and rescues growth of ROSA$^{KIT\ D816V}$ cells (n=3). Two-way ANOVA as in (E).

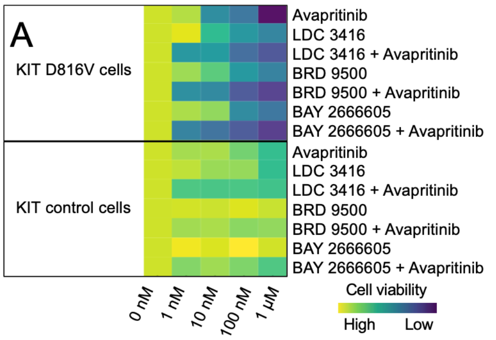


| IC50 values (nM) | | |
|---|---|---|
| | KIT D816V cells | KIT control cells |
| Avapritinib | 83.4 | 2902.1 |
| LDC 3416 | 403.0 | ≥$10^4$ |
| LDC 3416 + Avapritinib | 18.3 | >$10^4$ |
| BRD 9500 | 407.4 | >$10^4$ |
| BRD 9500 + Avapritinib | 2.6 | >$10^4$ |
| BAY 2666605 | 374.5 | >$10^4$ |
| BAY 2666605 + Avapritinib | 0.53 | >$10^4$ |

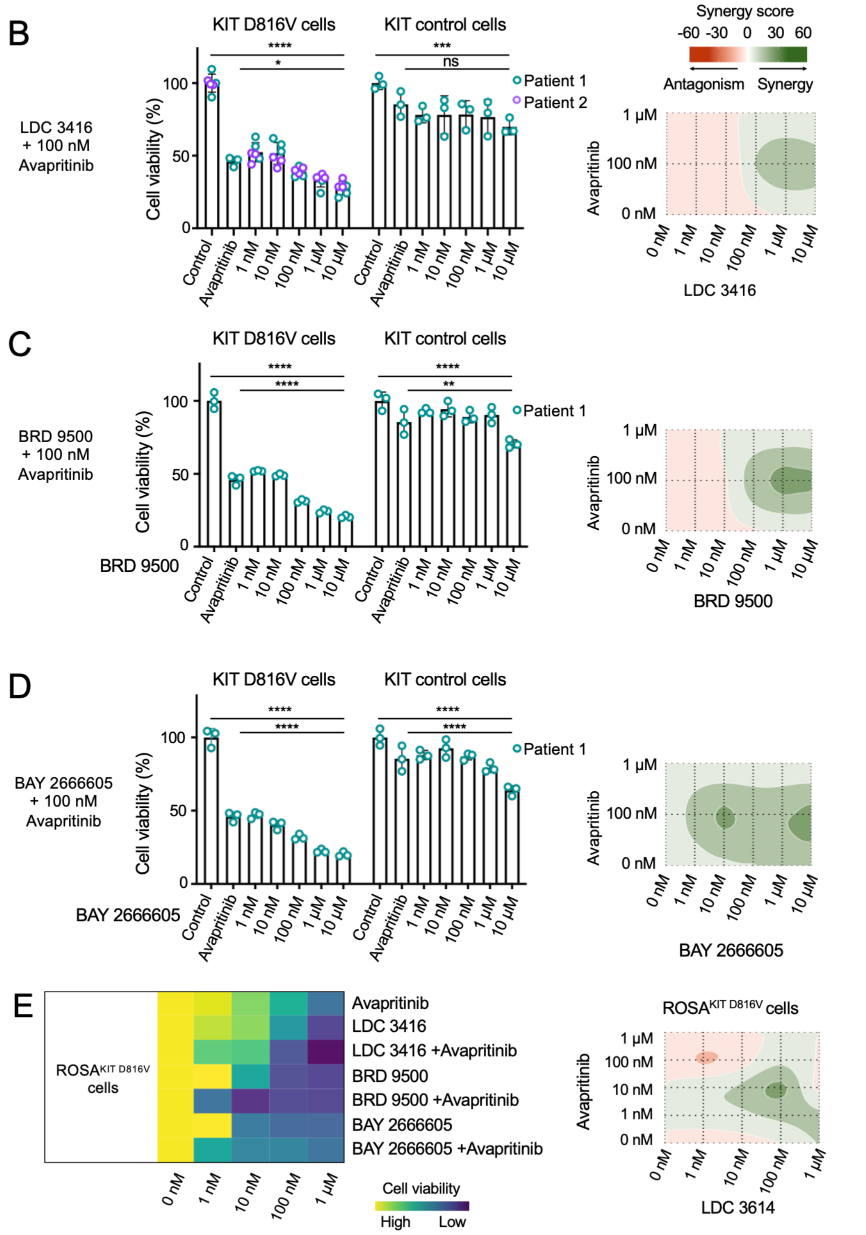


**Supplementary Figure 10. Synergy of molecular glues LDC 3416, BRD 9500 and BAY 2666605 and TKI Avapritinib in killing KIT D816V cells.**

(A) The molecular glues LDC 3416, BRD 9500 and BAY 2666605 synergizes with TKI Avapritinib (100 nM) in killing KIT D816V cells. Growth inhibition is depicted in heatmap format (yellow, high cell viability; blue, low cell viability). There is no or very little growth inhibition by compounds in KIT control cells (yellow to light green color code). IC50 values of synergy of molecular glues with TKI Avapritinib in KIT D816V cells are shown (n=3).

(B-D) Synergy of the molecular glues LDC 3416, BRD 9500 and BAY 2666605 with TKI Avapritinib of (A) is shown in dose response curves (left); Control, untreated; Avapritinib, 100 nM. Synergy representation by Combenefit analysis with Avapritinib concentrations 100 nM and 1 µM (right); green color code shows synergy (n=3-6, 3 independent experiments). Two-way ANOVA, * p≤0.05, ** p<0.01, *** p<0.001, and **** p<0.0001; ns, not significant.

(E) Synergy of molecular glues LDC 3416, BRD 9500 and BAY 2666605 (1 nM to 1 µM) and TKI Avapritinib (100 nM) in $ROSA^{KIT\ D816V}$ cells by MTT assay. Cell viability is shown in heatmap format (left; yellow, high cell viability; blue, low cell viability; n=3). Synergy of LDC 3416 with TKI Avapritinib in $ROSA^{KIT\ D816V}$ cells is shown by Combenefit analysis and color coded (right; green for synergy).

A

| | | KIT D816V | Treatment |
|---|---|---|---|
| Patient 47 | MCL + CMML-0 | 4% (BM) | Avapritinib |
| Patient 50 | MCL | 21% (BM) | Avapritinib |
| Patient 59 | MCL + MDS IB1 | 25% (BM) | Avapritinib |
| Patient control 39 | na | na | na |
| Patient control 43 | na | na | na |

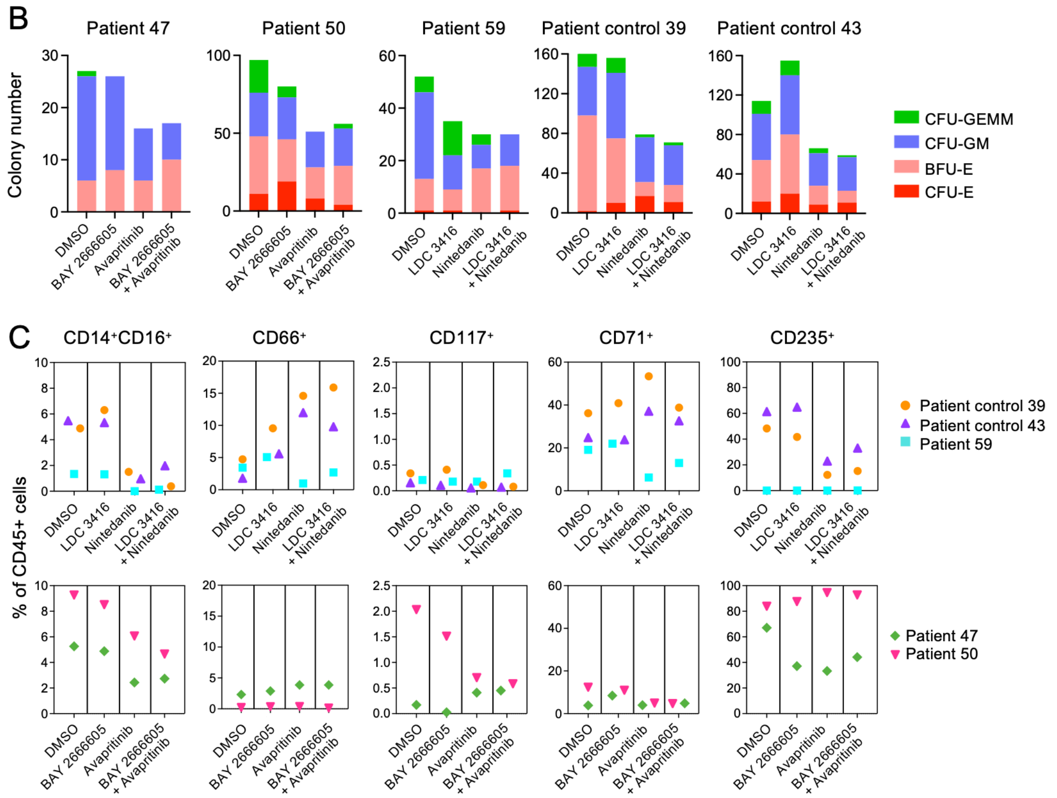


**Supplementary Figure 11. Molecular glue and TKI treatment of KIT D816V MCL patient BM samples.**

(A) MCL patient and patient control BM samples. Disease status, KIT D816V allele burden and treatment are indicated. MCL – CMML-0 refers to MCL with chronic myelomonocytic leukemia (CMML) stage 0. MCL – MDS IB1 refers to MCL with myelodysplastic syndrome (MDS) with increase blasts (IB) type 1. na, not applicable.

(B) SM patient bone marrow samples were treated with molecular glues LDC 3416 and BAY 2666605 and TKI Avapritinib and Nintedanib, and combinations thereof as indicated in colony assays and the number of CFU-GEMM, CFU-GM, BFU-E and CFU-E were assessed. Vehicle control, DMSO. CFU-GEMM, colony forming unit – granulocyte, erythrocyte, monocyte, megakaryocyte; CFU-GM, colony forming unit – granulocyte, macrophage; BFU-E, burst-forming unit – erythroid; CFU-E, colony-forming unit – erythroid.

(C) Phenotyping of CFU colonies of (B) by flow cytometry. Myeloid cells, $CD14^{+}CD16^{+}$; granulocytes, $CD66^{+}$; progenitors and mast cells, $CD117^{+}$; erythroid cells, $CD71^{+}$ (transferrin receptor) and $CD235a^{+}$ (glycphorin A).

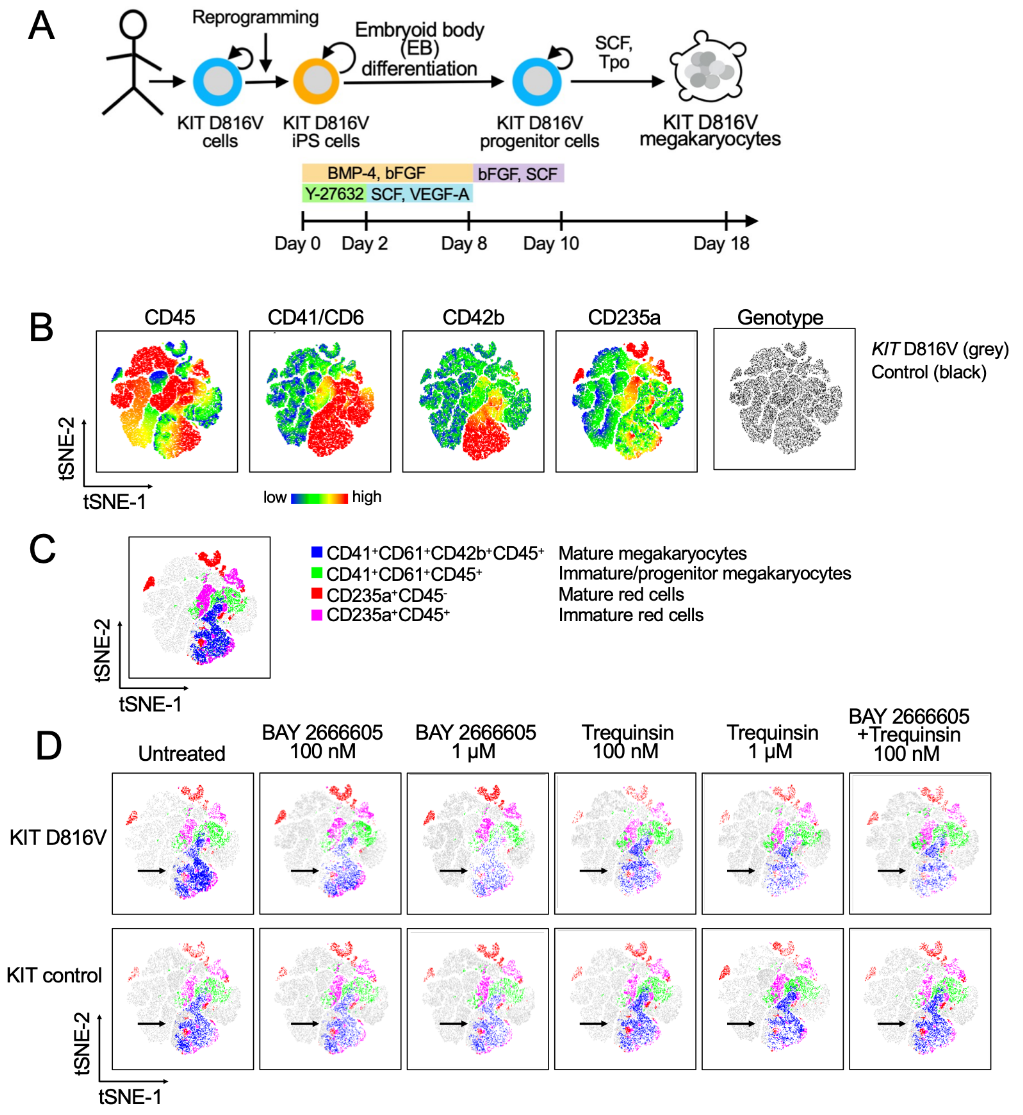


**Supplementary Figure 12. BAY 2666605 targets KIT D816V megakaryocytes.**

(A) Schematic representation of KIT D816V iPS cell differentiation into KIT D816V hematopoietic stem/progenitor cells and further into KIT D816V megakaryocytes. Isogenic iPS cells without *KIT* D816V mutation were used as controls. Bent arrows indicated self-renewal. iPS cell differentiation is induced by spin embryoid body (EB) formation in 96-well format and sequential addition of compounds and specific cytokines as indicated for generation and expansion of KIT D816V progenitor cells and their differentiation into KIT D816V megakaryocytes (SCF, Tpo).

(B) t-SNE plots of KIT D816V cells and KIT control cells upon megakaryocyte differentiation depicting expression of surface markers CD45, CD41, CD61, CD42b, and CD235a by flow cytometry. Color scale indicates relative expression levels (low to high). Genotype distribution is shown for KIT D816V cells and KIT control cells (grey and black, respectively, right panel).

(C) t-SNE clustering and annotation of cell populations: CD41$^+$CD61$^+$CD42b$^+$CD45$^+$ mature megakaryocytes (blue), CD41$^+$CD61$^+$CD45$^+$ immature/progenitor megakaryocytes (green), CD235a$^+$CD45$^-$ mature red cells (red), and CD235a$^+$CD45$^+$ immature erythroid cells (magenta).

(D) t-SNE plots of KIT D816V cells and KIT control cells (top and bottom rows, respectively) upon treatment with BAY 2666605 (100 nM, 1 μM), Trequinsin (100 nM, 1 μM), and a combination thereof, and untreated control. Arrows indicate shifts in the megakaryocyte population (blue).

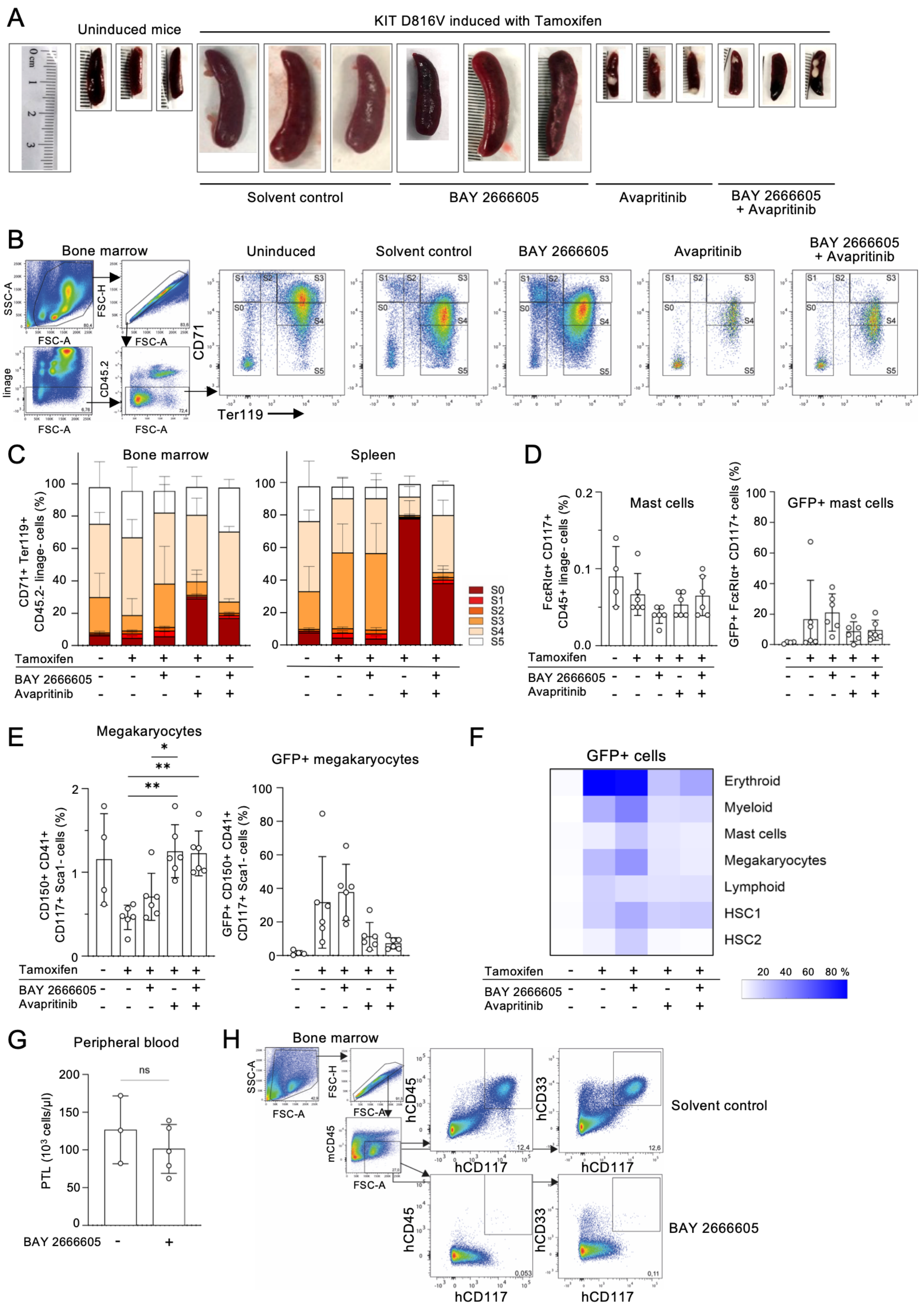


**Supplementary Figure 13. Activity of BAY 2666605 and Avapritinib on KIT D816V cells in SCL-GFP-KIT D816V mice and on ROSA$^{KIT\ D816V}$ cells in NSG mice.**

(A) Spleen size of SCL-GFP-KIT D816V mice (ruler in cm) without and with Tamoxifen induction and treatment with BAY 2666605, Avapritinib or both was analysed at Day 28 as in Figure 6A. Uninduced mice refers to spleen size of healthy control.

(B) Gating strategy for erythroid precursors in bone marrow of SCL-GFP-KIT D816V mice by flow cytometry and representative image of the impact of BAY 2666605, Avapritinib or both on CD71$^{+}$Ter119$^{+}$ erythroid cells at Day 28. S0-S5 represent different stages of erythroid development as indicated.

(C) Impact of BAY 2666605, Avapritinib or both on CD71$^{+}$Ter119$^{+}$ erythroid cells of different stages of development. S0-S5 as in (B) in bone marrow and spleen.

(D) Frequency of total mast cells (FcεRIα$^{+}$CD117$^{+}$ linage$^{-}$ CD45$^{+}$ cells) and GFP$^{+}$ mast cells in bone marrow after treatment as in (C).

(E) Frequency of total megakaryocytes (CD150$^{+}$CD41$^{+}$CD117$^{+}$Sca1$^{-}$ cells) and GFP$^{+}$ megakaryocytes in bone marrow after treatment as in (C).

(F) Heat map shows the impact of BAY 2666605, Avapritinib or both on distinct GFP$^{+}$ cell subsets in bone marrow (erythroid, myeloid and lymphoid cells, hematopoietic stem cells, HSC1 and HSC2, mast cells and megakaryocytes as defined in Pelusi et al., 2017).

(G) Total platelets (PTL) in peripheral blood after treatment with BAY 2666605 in NSG mice engrafted with ROSA$^{KIT\ D816V}$ cells.

(H) Gating strategy for ROSA$^{KIT\ D816V}$ cells in NSG mice by staining for hCD33, hCD45 and hCD117 and flow cytometry. Impact of BAY 2666605 on ROSA$^{KIT\ D816V}$ cells in NSG mice at Day 16 is shown as in Figure 6D.

Statistics: Brown–Forsythe and Welch one-way ANOVA followed by Dunnett T3 post-hoc tests; for non-normally distributed data: Mann–Whitney test and Kruskal–Wallis test with Dunn's multiple comparisons: not significant: all non-indicated statistics; * p <0.05; ** p <0.01; ns, not significant.

| Commercial name or compound ID | Patient 1 KIT D816V | Patient 1 KIT control | Patient 2 KIT D816V | Patient 2 KIT control | HES cells KIT D816V | HES cells KIT control |
|---|---|---|---|---|---|---|
| | IC50 (µM) | IC50 (µM) | IC50 (µM) | IC50 (µM) | IC50 (µM) | IC50 (µM) |
| Dasatinib | 0,39 | >10 | 0,09 | >10 | 0,85 | >10 |
| **LDC193416** | 0,30 | 7,40 | 0,72 | >10 | 1,88 | >10 |
| Cerdulatinib | 0,56 | >10 | 0,09 | 0,11 | 0,47 | >10 |
| Disulfiram | 0,18 | 3,04 | 0,17 | 5,93 | 0,16 | >10 |
| Milciclib | 0,63 | >10 | 0,19 | 0,35 | 0,65 | >10 |
| AZD7762 | 0,05 | 0,63 | 0,01 | 0,92 | 0,10 | >10 |
| LDC043400 | 0,64 | 7,33 | 1,56 | >10 | ND | >10 |
| PP121 | 0,16 | 1,83 | 0,06 | 0,59 | 0,20 | 2,72 |
| R406 | 0,52 | 5,35 | 0,18 | 0,47 | 0,66 | 3,00 |
| Gusacitinib | 1,13 | >10 | 0,11 | 0,10 | 0,59 | >10 |
| TWS119 | 1,39 | >10 | 0,30 | >10 | >10 | >10 |
| Delanzomib | 0,14 | 0,97 | 0,11 | 0,64 | 0,01 | >10 |
| ML291 | 1,44 | 9,77 | 0,79 | 4,73 | 0,66 | >10 |
| Avapritinib | 1,32 | 8,57 | 0,28 | >10 | 0,61 | >10 |
| LDC001816 | 0,95 | 6,00 | 0,45 | 1,74 | 1,08 | 6,23 |
| LDC001527 | 1,72 | >10 | 0,31 | >10 | 1,76 | >10 |
| BETd-246 | 0,01 | 0,05 | 0,01 | 0,05 | 0,01 | >10 |
| ARV-825 | 0,02 | 0,08 | 0,01 | 0,15 | 0,02 | >10 |
| LDC001386 | 2,07 | >10 | 1,98 | >10 | 1,09 | >10 |
| SU5612 | 2,26 | >10 | 0,76 | 6,91 | >10 | >10 |
| dBET6 | 0,02 | 0,07 | 0,25 | >10 | 0,01 | >10 |
| Staurosporine | 0,06 | 0,25 | 0,13 | 2,66 | 0,06 | 1,98 |
| LOC14 | 1,52 | 6,28 | 1,25 | 6,28 | 0,82 | 8,07 |
| LDC001813 | 1,35 | 5,46 | 0,48 | >10 | 0,40 | >10 |
| Abemaciclib | 1,85 | 7,27 | 1,12 | >10 | 2,46 | 3,92 |
| Momelotinib | 2,59 | >10 | 0,83 | 1,74 | 2,96 | >10 |
| CZC-25146 | 2,58 | 9,36 | 0,86 | 5,52 | 3,66 | 7,96 |
| BET Degrader-1 | 0,03 | 0,12 | 0,02 | 0,68 | 0,02 | >10 |
| A-674563 | 2,99 | >10 | 1,74 | >10 | ND | >10 |
| Vactosertib | 3,04 | >10 | 0,81 | 8,97 | 3,03 | >10 |
| Bx-795 | 3,12 | >10 | 1,61 | 8,91 | 8,64 | >10 |
| GLPG0634 analoge | 3,13 | >10 | 0,26 | >10 | 0,43 | >10 |
| Ixazomib | 0,22 | 0,68 | 0,21 | 0,42 | 0,48 | >10 |
| LDC001719 | 1,33 | 4,02 | 0,28 | 0,43 | 4,33 | >10 |
| R-268712 | 3,31 | >10 | 0,94 | >10 | 6,09 | >10 |
| Bms-38703 | 0,26 | 0,76 | 0,04 | >10 | 0,23 | >10 |
| LDC208887 | 1,01 | 2,98 | 0,72 | 4,78 | 0,34 | 1,42 |
| PF-03814735 | 1,63 | 4,76 | 0,25 | 0,38 | 2,20 | >10 |
| LDC001443 | 3,59 | >10 | 4,69 | >10 | 1,68 | 0,45 |
| Prodigiosine | 1,11 | 3,06 | 4,34 | 9,63 | 7,36 | >10 |
| LDC001634 | 0,44 | 1,22 | 0,27 | 0,30 | 1,05 | >10 |
| Onx-914 | 0,06 | 0,15 | 0,07 | 0,35 | 0,06 | >10 |
| LDC001449 | 3,79 | >10 | 2,24 | >10 | 1,19 | >10 |
| LDC205477 | 2,62 | 6,58 | >10 | >10 | >10 | >10 |
| LDC001546 | 0,37 | 0,93 | 0,14 | >10 | 0,17 | >10 |
| LDC001569 | 0,35 | 0,86 | 0,14 | 2,83 | 1,70 | >10 |
| Bms-214662 | 0,04 | 0,10 | 0,08 | 0,19 | 0,04 | 0,17 |
| Bortezomib | 0,01 | 0,02 | 0,01 | 0,02 | 0,01 | >10 |
| LDC001555 | 1,67 | 3,72 | 1,28 | 0,57 | 1,29 | >10 |
| LDC001393 | 4,53 | >10 | 1,59 | 5,82 | 8,76 | 8,88 |
| A-485 | 4,60 | >10 | 1,30 | >10 | >10 | >10 |
| LDC000267 | 1,67 | 3,60 | 1,36 | 1,48 | 1,31 | >10 |
| LDC001095 | 2,38 | 5,12 | 3,49 | 3,14 | 3,98 | >10 |
| LDC001538 | 0,92 | 1,99 | 0,46 | >10 | 0,57 | >10 |
| LDC001620 | 1,97 | 4,20 | 1,08 | >10 | 1,00 | >10 |
| MG-132 | 0,29 | 0,60 | 0,28 | 0,50 | 0,38 | 1,03 |
| LDC001808 | 1,21 | 2,50 | 0,60 | 3,14 | 1,68 | 9,47 |
| Rocaglaol | 0,04 | 0,09 | 0,01 | 0,02 | 0,25 | >10 |
| ARV-771 | 0,17 | 0,35 | 0,07 | 0,30 | 0,13 | >10 |
| Ixazomib | 0,12 | 0,25 | 0,10 | 0,19 | 0,49 | >10 |
| LDC001608 | 1,46 | 2,84 | 0,65 | >10 | 0,76 | >10 |
| R547 | 2,26 | 4,38 | 1,37 | 1,21 | 1,74 | >10 |
| PF-477736. | 0,34 | 0,66 | 0,11 | 0,72 | 2,07 | >10 |
| LDC001793 | 3,90 | 7,20 | 3,35 | 2,86 | 3,04 | >10 |
| LDC001600 | 2,10 | 3,87 | 1,54 | 2,57 | 4,09 | >10 |
| Anisomycin | 0,10 | 0,18 | 0,07 | 0,11 | 0,10 | 1,17 |

| | | | | | | |
|---|---|---|---|---|---|---|
| BETd-260 | 0,00 | 0,01 | 0,00 | 0,01 | 0,00 | >10 |
| CZC54252 | 0,96 | 1,72 | 0,33 | 1,98 | 1,90 | 5,03 |
| Homoharringtonine | 0,04 | 0,08 | 0,01 | ND | 0,06 | >10 |
| Midostaurin | 0,59 | 1,00 | 2,25 | >10 | 0,55 | 1,00 |
| Flavopiridol | 0,14 | 0,24 | 0,09 | 0,21 | 0,09 | >10 |
| LDC046529 | 0,75 | 1,27 | 0,42 | 1,57 | 0,84 | 3,04 |
| LDC001740 | 4,39 | 7,22 | 2,91 | 2,52 | 3,98 | >10 |
| LDC001456 | 6,16 | >10 | 7,12 | >10 | >10 | >10 |
| LDC211813 | 0,79 | 1,26 | 0,66 | 1,14 | 0,41 | 7,25 |
| Carfilzomib | 0,03 | 0,05 | 0,02 | 0,19 | 0,00 | >10 |
| SB1317 | 0,12 | 0,18 | 0,04 | 7,45 | 0,04 | >10 |
| SCH-1473759 | 0,17 | 0,26 | 0,04 | 0,05 | 0,18 | 8,31 |
| MZ 1 | 0,63 | 0,98 | 0,20 | 0,48 | 0,34 | >10 |
| LDC001806 | 0,27 | 0,41 | 0,10 | >10 | 0,08 | >10 |
| 5-Nonyloxytryptamine | 2,70 | 4,09 | 2,22 | 8,46 | ND | 8,89 |
| LDC001562 | 3,82 | 5,76 | 2,05 | >10 | 1,92 | >10 |
| LDC207258 | 0,97 | 1,46 | 0,43 | 0,36 | 0,52 | >10 |
| LDC245352 | 6,74 | >10 | 2,28 | 3,05 | >10 | >10 |
| BX-912 | 0,81 | 1,19 | 0,32 | 1,24 | 1,70 | 3,51 |
| LDC001776 | 6,84 | >10 | 2,91 | 0,47 | >10 | >10 |
| LDC001694 | 2,71 | 3,93 | 1,96 | 2,19 | 1,72 | >10 |
| LDC001794 | 2,35 | 3,35 | 1,55 | >10 | 1,39 | >10 |
| Staurosporine | 1,78 | 2,49 | 0,74 | 1,85 | 2,10 | 5,47 |
| PD 407824 | 2,44 | 3,34 | 1,12 | 1,57 | 2,77 | >10 |
| MZP-54 | 0,60 | 0,82 | 0,18 | 0,42 | 0,39 | >10 |
| LDC001831 | 5,28 | 7,17 | 5,99 | 5,58 | ND | >10 |
| AZD-5438 | 3,19 | 4,32 | 1,21 | >10 | 1,86 | >10 |
| LDC000526 | 1,96 | 2,63 | 0,91 | >10 | 1,02 | >10 |
| LDC001724 | 2,10 | 2,81 | 1,12 | >10 | 1,17 | >10 |
| LDC000928 | 3,29 | 4,33 | 1,46 | 2,99 | 4,27 | >10 |
| THZ531 | 0,11 | 0,14 | 0,07 | 7,23 | 0,07 | ND |
| MG-115 | 0,46 | 0,58 | 0,27 | 0,88 | 0,35 | >10 |
| LDC208431 | 0,01 | 0,01 | 0,00 | 0,00 | 0,12 | >10 |
| LDC001593 | 1,78 | 2,26 | 0,99 | >10 | 1,17 | >10 |
| LDC001769 | 4,96 | 6,31 | 2,17 | >10 | 3,34 | >10 |
| LDC001454 | 3,20 | 4,07 | 1,94 | >10 | 2,43 | 2,84 |
| Berzosertib | 3,36 | 4,20 | 1,50 | 1,88 | 2,46 | 9,35 |
| LDC001130 | 7,99 | >10 | 2,34 | >10 | 1,46 | 0,87 |
| Triptolide | 0,01 | 0,02 | 0,01 | ND | 0,01 | >10 |
| LDC001408 | 8,11 | >10 | 2,43 | 5,86 | >10 | >10 |
| LDC001717 | 5,83 | 7,15 | 3,27 | 8,52 | 6,31 | 9,52 |
| Plicamycin. | 0,95 | 1,15 | 1,22 | 1,37 | 0,22 | >10 |
| Luminespib | 0,00 | 0,00 | 0,00 | 0,01 | 0,00 | ND |
| CI-387785 | 2,10 | 2,46 | 1,73 | 1,77 | 1,80 | 1,81 |
| LDC001632 | 2,96 | 3,44 | 2,09 | 2,95 | 0,15 | >10 |
| LDC001536 | 6,07 | 7,05 | 2,83 | >10 | 2,96 | >10 |
| CGP60474 | 0,07 | 0,08 | 0,03 | >10 | ND | >10 |
| LDC001702 | 8,69 | >10 | 3,30 | >10 | >10 | >10 |
| Tesevatinib | 2,20 | 2,48 | 1,79 | 3,12 | 2,73 | 5,57 |
| LDC001560 | 2,49 | 2,79 | 1,65 | >10 | 0,93 | >10 |
| LDC001288 | 9,23 | >10 | 1,62 | >10 | >10 | >10 |
| BAY 11-7821 | 0,81 | 0,86 | 0,97 | 2,14 | 0,32 | 0,91 |
| LDC001644 | 2,02 | 2,15 | 0,80 | 2,64 | 0,40 | 2,67 |
| LDC212037 | >10 | >10 | 7,01 | 2,48 | 1,39 | 6,96 |
| Capivasertib | 1,27 | 1,33 | 0,37 | 4,68 | 1,77 | 1,24 |
| LDC000003 | 3,11 | 3,12 | 1,45 | 1,01 | 1,09 | >10 |
| Lestaurtinib | 0,16 | 0,16 | 0,02 | 0,09 | 0,18 | >10 |
| Desloratadine | >10 | >10 | >10 | >10 | >10 | >10 |
| LDC001175 | >10 | >10 | >10 | >10 | >10 | >10 |
| LDC001181 | >10 | >10 | >10 | >10 | 1,71 | >10 |
| LDC001198 | >10 | >10 | >10 | >10 | 2,56 | 3,09 |
| LDC001255 | >10 | >10 | >10 | 6,14 | 3,89 | 9,01 |
| LDC001458 | >10 | >10 | >10 | >10 | >10 | >10 |
| LDC001799 | >10 | >10 | >10 | >10 | 2,19 | >10 |
| LDC037359 | >10 | >10 | >10 | >10 | >10 | >10 |
| PIK-75 | >10 | >10 | >10 | >10 | >10 | >10 |
| Prochlorperazine Edisy | >10 | >10 | >10 | >10 | ND | >10 |
| Itopride | >10 | >10 | >10 | >10 | >10 | >10 |
| Apomorphine | >10 | >10 | 7,91 | >10 | >10 | >10 |

| | | | | | | |
|---|---|---|---|---|---|---|
| Selamectin | >10 | >10 | >10 | >10 | >10 | >10 |
| JNK-IN-7 | >10 | >10 | >10 | >10 | 1,07 | >10 |
| Haloxyfop-P-Methyl | >10 | >10 | >10 | >10 | >10 | >10 |
| LDC206648 | >10 | >10 | >10 | >10 | >10 | 1,45 |
| LDC209902 | >10 | >10 | >10 | >10 | >10 | >10 |
| GSK503 | >10 | >10 | >10 | >10 | >10 | >10 |
| Tazemetostat | >10 | >10 | >10 | >10 | >10 | >10 |
| 10-Hydroxycamptothecin | 0,49 | 0,47 | 0,21 | 0,09 | 0,87 | 1,51 |
| LDC204014 | 2,52 | 2,39 | 1,65 | 0,84 | 2,38 | >10 |
| LDC001725 | 2,43 | 2,25 | 1,55 | >10 | 1,06 | >10 |
| LDC001734 | >10 | >10 | 7,27 | 5,97 | >10 | >10 |
| LDC245344 | >10 | 9,19 | >10 | >10 | >10 | >10 |
| LDC000887 | 0,44 | 0,40 | 0,23 | 0,43 | 0,37 | 1,64 |
| Flavopiridol | 0,67 | 0,61 | 0,23 | 0,57 | 0,28 | >10 |
| HDACs/mTOR Inhibitor 1 | 0,02 | 0,02 | 0,01 | 0,01 | 0,01 | >10 |
| LDC000920 | 5,12 | 4,50 | 2,07 | 7,61 | 6,46 | >10 |
| LDC001572 | 2,85 | 2,49 | 1,34 | >10 | 1,77 | 8,06 |
| LDC001539 | >10 | 8,57 | 2,04 | >10 | ND | >10 |
| LDC001585 | >10 | >10 | 9,54 | 8,82 | 1,25 | >10 |
| LDC208628 | 2,37 | 2,00 | 1,23 | 2,01 | 1,94 | 8,31 |
| Torkinib | 0,47 | 0,39 | 0,14 | 0,58 | 0,63 | 1,63 |
| LDC001827 | 0,77 | 0,63 | 0,24 | 0,48 | 0,56 | >10 |
| LDC193458 | 2,48 | 2,00 | 2,46 | 4,11 | 1,83 | 4,02 |
| Cucurbitacin | 0,09 | 0,07 | 0,02 | 0,12 | 0,04 | >10 |
| Auranofin | 0,30 | 0,23 | 0,16 | 1,49 | 0,23 | ND |
| LDC001805 | 3,56 | 2,72 | 1,63 | >10 | 0,31 | >10 |
| Brigatinib | 3,10 | 2,36 | 0,81 | 2,27 | 4,30 | 2,28 |
| Atuveciclib | 4,48 | 3,32 | 1,00 | >10 | 2,33 | >10 |
| WZ8040 | 1,54 | 1,13 | 1,06 | 1,09 | 1,21 | 1,97 |
| LDC001824 | >10 | >10 | 7,41 | 4,51 | ND | >10 |
| LDC001571 | 5,81 | 4,15 | 2,10 | >10 | 1,76 | >10 |
| LDC193464 | 1,93 | 1,37 | 1,62 | 2,54 | 2,36 | 1,81 |
| Oprozomib | 0,42 | 0,30 | 0,19 | 1,63 | 0,01 | >10 |
| LDC198434 | 1,81 | 1,24 | 1,44 | 1,62 | 1,68 | >10 |
| CHZ868 | 1,08 | 0,73 | 2,23 | 0,21 | 1,43 | 5,94 |
| Corin | 0,43 | 0,29 | 0,29 | 0,43 | 0,36 | 9,15 |
| Ganetespib | 0,01 | 0,01 | 0,00 | 0,01 | 0,01 | ND |
| LDC001701 | 2,33 | 1,54 | 1,05 | 0,63 | ND | >10 |
| Apilimod | 2,07 | 1,36 | 2,62 | 2,11 | 1,52 | 4,53 |
| Puromycin | 1,39 | 0,90 | 0,96 | 1,90 | 0,35 | >10 |
| PR-825 | 0,09 | 0,06 | 0,06 | 0,13 | 0,06 | >10 |
| LDC206735 | 1,39 | 0,85 | 0,45 | 1,06 | 1,65 | >10 |
| CHIR-265 | 2,04 | 1,12 | 1,99 | 2,60 | 4,90 | 1,17 |
| LDC001671 | 3,96 | 2,16 | 1,34 | >10 | ND | >10 |
| JNJ-26481585 | 0,02 | 0,01 | 0,01 | 0,01 | 0,03 | >10 |
| Ponatinib | 1,41 | 0,75 | 0,65 | 2,03 | 1,24 | >10 |
| Givinostat | 0,34 | 0,18 | 0,27 | 0,45 | 0,32 | >10 |
| Dactinomycin | 0,01 | 0,01 | 0,01 | ND | 0,02 | >10 |
| (R)-CR8 | 0,25 | 0,13 | 0,11 | 0,43 | 0,08 | 1,90 |
| LDC210284 | 0,02 | 0,01 | 0,01 | 0,01 | 0,02 | >10 |
| Romidepsin | 0,02 | 0,01 | 0,16 | 0,21 | 0,01 | ND |
| GNE-272 | >10 | 4,86 | 6,44 | 6,06 | >10 | >10 |
| TG101209 | 1,30 | 0,63 | 0,60 | 0,49 | 1,22 | 1,93 |
| LDC200345 | 4,22 | 2,05 | 3,44 | 2,10 | 3,12 | 2,82 |
| LDC193472 | >10 | 4,85 | >10 | 0,66 | >10 | >10 |
| CP-681301 | 1,55 | 0,73 | 1,41 | 1,05 | 0,39 | >10 |
| Trichostatin-A (TSA) | 0,12 | 0,06 | 0,10 | 0,09 | 0,09 | >10 |
| Sunitinib | 1,46 | 0,67 | 0,84 | 1,43 | 1,55 | 3,37 |
| BAG 956 | 0,48 | 0,20 | 0,18 | 0,37 | 1,00 | 0,33 |
| Dovitinib | 0,87 | 0,36 | 0,52 | 5,21 | 2,80 | 2,31 |
| CA3 | 1,07 | 0,44 | 3,18 | 0,61 | 0,83 | 2,24 |
| Fimepinostat | 0,01 | 0,00 | 0,00 | ND | 0,01 | 7,17 |
| M344 | 0,93 | 0,37 | 0,59 | 0,84 | 0,79 | >10 |
| HDAC-IN-3 | 0,48 | 0,19 | 0,31 | 0,25 | 0,37 | >10 |
| CCT251545 analogue | 1,71 | 0,69 | 1,84 | 2,05 | 2,15 | 3,45 |
| LDC001788 | 0,30 | 0,12 | 0,10 | >10 | 0,07 | >10 |

| | | | | | | |
|---|---|---|---|---|---|---|
| Sapanisertib | 0,20 | 0,08 | 0,03 | 0,30 | 0,13 | 8,17 |
| Pracinostat. | 0,43 | 0,16 | 0,37 | 0,35 | 0,45 | 0,91 |
| LDC211625 | 1,00 | 0,36 | 0,51 | 0,25 | 0,48 | 0,57 |
| Dacinostat | 0,25 | 0,09 | 0,20 | 0,38 | 0,31 | >10 |
| SB-743921 | 1,73 | 0,54 | 1,07 | 1,92 | 1,32 | 2,13 |
| Vorinostat | 0,68 | 0,21 | 0,55 | 0,26 | 0,79 | >10 |
| LDC207256 | 0,04 | 0,01 | 0,01 | 0,04 | 0,04 | 0,03 |
| Monensin | 0,79 | 0,24 | 0,85 | 1,57 | 0,03 | 0,33 |
| IKK-2 Inhibitor VI | 1,82 | 0,52 | 0,41 | 0,25 | 1,06 | 3,91 |
| LDC245350 | 8,21 | 2,29 | 1,53 | 2,91 | 5,28 | 9,02 |
| CAY10603 | 0,83 | 0,22 | 0,43 | 0,42 | 0,56 | >10 |
| Oxamflatin | 0,54 | 0,14 | 0,31 | 0,74 | 0,48 | >10 |
| AZD-8055 | 0,40 | 0,10 | 0,06 | 0,09 | 0,40 | 0,79 |
| Camptothecin | 11,91 | 2,78 | 4,01 | 1,33 | >10 | >10 |
| CUDC-101 | 0,33 | 0,08 | 0,18 | 0,06 | 0,18 | >10 |
| Clofarabine | 9,36 | 2,11 | 1,70 | 2,35 | 6,97 | >10 |
| ABT-737 | 1,14 | 0,22 | 0,47 | 0,59 | 0,62 | 0,57 |
| Vistusertib | 1,53 | 0,27 | 0,19 | 0,61 | 0,72 | 4,00 |
| Torin 2 | 0,08 | 0,01 | 0,03 | 0,05 | 0,06 | ND |
| Belinostat | 0,75 | 0,12 | 0,67 | 0,42 | 0,51 | >10 |
| Apitolisib | 0,76 | 0,11 | 0,16 | 0,45 | 0,61 | 0,21 |
| LDC206769 | 6,48 | 0,91 | 1,49 | 1,27 | 3,81 | 1,41 |
| LDC193452 | 0,40 | 0,03 | 0,07 | 0,06 | 0,22 | 0,06 |
| ABBV-744 | 4,12 | 0,19 | 0,34 | 0,21 | 1,38 | 4,16 |
| Hesperadin | 5,39 | 0,23 | 0,57 | 0,05 | >10 | >10 |
| Brefeldin A | 2,53 | 0,09 | 0,42 | 0,18 | 0,15 | >10 |
| LDC206786 | 0,14 | 0,00 | 0,01 | 0,01 | 0,08 | ND |
| LDC001477 | 2,59 | ND | 1,73 | >10 | ND | >10 |
| LDC001617 | 6,98 | ND | 0,76 | ND | 2,80 | >10 |
| LDC202293 | >10 | ND | 0,63 | >10 | 0,69 | >10 |
| LDC203955 | 2,44 | ND | 1,50 | 5,09 | 1,17 | 5,97 |
| Dinaciclib | ND | ND | 0,01 | >10 | 0,01 | >10 |
| THZ1 | ND | 0,18 | 0,05 | >10 | 0,06 | 0,28 |

**Supplementary Table 1. IC50 values of 234 compounds subjected to KIT D816V cell hit validation.**

KIT D816V cells and isogenic KIT control cells derived from iPS cells (patient 1 and 2) and CRISPR/Cas9 engineered KIT D816V human ES cells and unmutated KIT control (HES cells; Toledo et al., 2021) were treated with compounds in dose response curves (0.003 to 10 µM) and IC50 values upon drug treatment are shown. IC50 values above 10 µM were not resolved (>10). LDC 3416 is highlighted. Commercial compound names if applicable or compound identifications (ID) are shown. Compounds effective in reducing of cell viability are color coded in red. Compounds, which were essentially ineffective, are in green. ND, not determined.

**Patient 39**. A 68-year-old woman was referred by the Department of Allergology due to multiple antibiotic allergies and an elevated serum tryptase level of 26 ng/ml. A BM biopsy excluded an underlying SM. *KIT* D816V and also *ASXL1* and *RUNX1* mutations were negative. Testing for hereditary alpha-tryptasemia (HαT) was not performed. BM biopsy did not show any overt hematological malignancies and the BM sample was classified as Patient control 39.

**Patient 43.** A 55-year-old man presented to the Department of Allergology following an anaphylactic reaction to a wasp sting. Further assessment revealed an elevated serum tryptase level of 14 ng/ml and the patient was referred to our Department of Hematology, Oncology and Stem Cell Transplantation for further diagnostic work up, as underlying SM was suspected. SM was ruled out by BM biopsy and molecular testing on *KIT* D816V mutation. There were no obvious hematological malignancies and the BM sample was classified as Patient control 43.

**Patient 47.** A 75-year-old man was diagnosed with SM-AHN. The SM part was diagnosed as MCL and AHN part as CMML-0. Initial serum tryptase level was 259 ng/ml. Molecular diagnostic revealed *KIT* D816V, *ASXL1* and *SRSF2* mutations. Midostaurin was not well tolerated and therefore therapy was switched to Avapritinib. BM sample for analysis was taken after 6 months of Avapritinib treatment (dose 100 mg/day). Patient's SM reached a complete remission with normalized tryptase levels and absence of *KIT* D816V mutation after two years of treatment. One year later the patient showed progress of AHN to MDS IB II with 14% blasts in BM smears, but still no relapse of SM. *KIT* D816V was still negative. Now, additional to *ASXL1* and *SRSF2* mutations also *IDH2*, *MPL* and *TET2* mutations were detected. Treatment with Avapritinib was stopped and the patient received therapy with azacitidine. The patient died one year after the MDS diagnosis.

**Patient 50.** A 76-year-old man was diagnosed with *KIT* D816V positive ISM for 12 years and then progressed to MCL. The patient presented with splenomegaly, weight loss and transfusion-dependent anemia due to recurrent gastrointestinal (GI) bleedings based on histological confirmed infiltration of SM in the gastrointestinal tract. The patient was treated with Midostaurin for two years and one year with Ripretinib, both without sufficient response. Treatment was then switched to Avapritinib. The patient was under Avapritinib treatment (dose 100 mg/day) for 6 months when the BM sample was taken and subjected to analysis. Serum tryptase level was 25.8 ng/ml. Eventually the patient reached a partial response (IWG MRT ECNM criteria and pure pathological remission criteria).

**Patient 59.** A 65-year-old man presented with weight loss, night sweat, fever and unexplained splenomegaly. He further developed hepatomegaly and ascites and eventually BM biopsy revealed the diagnosis of *KIT* D86V positive MCL-AHN. AHN was classified as MDS IB 1. BM analysis identified additional mutations in *SRSF2*, *RUNX1* and *TET2*. During treatment with Midostaurin the patient showed further disease progression and treatment was switched to Avapritinib. BM sample was taken and subjected to analysis when the patient was under treatment with Avapritinib (dose 100 mg/day) for 7 months. Serum tryptase level was 89 ng/ml. During Avapritinib treatment a partial remission was achieved with complete remission of C findings, and the patient could proceed with allogeneic stem cell transplantation (alloSCT). After alloSCT the SM part of the disease is currently still in remission, but the AHN part showed relapse and developed into an acute myeloid leukemia, which requires cytoreductive therapy.

**Supplementary Table 2. Information on BM samples for CFU assay.**

Patients disease history and information on the BM samples taken and analyzed. All patients provided informed consent to the use of samples for the study (Medical Faculty of RWTH Aachen University, Aachen, Germany, Ethics board reference number EK206/09).

| Gene | Primer sequence (5'-3') | |
|---|---|---|
| GAPDH | Forward | GAAGGTGAAGGTCGGAGTC |
| | Reverse | GAAGATGGTGATGGGATTTC |
| PDE3A | Forward | AAGCCCAGAGTGAATCCCG |
| | Reverse | GAAACTCGTCTCAACAAGCCAG |
| SLFN12 | Forward | GTGTTTGCTAAAGAGCCTGATTC |
| | Reverse | AGTGATGTCTCTGCCGTTGC |

**Supplementary Table 3. RT-qPCR primers.**

Nucleotide sequences of RT-qPCR primers used in the study are shown

| Antibody (Human) | Fluorochrome | Clone | Dilution | Cat # | RRID | Company |
|---|---|---|---|---|---|---|
| CD11b | BV510 | M1/70 | 1:100 | 101245 | AB_2561390 | Biolegend |
| CD14 | AF700 | M5E2 | 1:100 | 561029 | AB_396944 | BD Bioscience |
| CD16 | APC | HI16a | 1:100 | 2181016 | AB_2726150 | ImmunoTools |
| CD16 | FITC | REA423 | 1:100 | 130-113-392 | | Miltenyi Biotec |
| CD34 | PE | 581 | 1:100 | 343506 | AB_1731862 | Biolegend |
| CD41/CD61 | FITC | REA607 | 1:100 | 130-124-887 | AB_2819707 | Miltenyi Biotec |
| CD42b | APC | HIP1 | 1:100 | 303912 | AB_2113770 | Biolegend |
| CD45 | APC-Vio770 | REA747 | 1:200 | 130-110-773 | AB_2658250 | Miltenyi Biotec |
| CD66b | PE | 6/40c | 1:100 | 21609664 | | ImmunoTools |
| CD71 | FITC | CY1G4 | 1:100 | 334103 | AB_1236432 | Biolegend |
| CD117 | FITC | 104D2 | 1:100 | 11-1178-42 | AB_2572472 | eBioscience |
| CD235a | PE | REA175 | 1:100 | 130-120-613 | AB_2801777 | Miltenyi Biotec |
| **Antibody (Mouse)** | **Fluorochrome** | **Clone** | **Dilution** | **Cat #** | **RRID** | **Company** |
| B220 | PB | RA3-6B2 | 1:100 | 103230 | AB_492877 | Biolegend |
| B220 | Pe-Cy5 | RA3-6B2 | 1:100 | 103210 | AB_312995 | Biolegend |
| CD3 | APC | 17A2 | 1:100 | 100236 | AB_2561456 | Biolegend |
| CD3 | Pe-Cy5 | 17A2 | 1:100 | 555276 | AB_395700 | BD Pharmingen |
| CD4 | Pe-Cy5 | RM4-5 | 1:100 | 553050 | AB_394586 | BD Pharmingen |
| CD8a | Pe-Cy5 | 53-6.7 | 1:100 | 15-0081 82 | AB_468706 | Invitrogen |
| CD11b | PB | M1/70 | 1:100 | 101224 | AB_755986 | Biolegend |
| CD11b | PE cy 7 | M1/70 | 1:100 | 101216 | AB_312799 | Biolegend |
| CD11b | Pe-Cy5 | M1/70 | 1:100 | 101210 | AB_312793 | Biolegend |
| CD16/CD32 | APC | 93 | 1:100 | 17-0161-81 | AB_469356 | Invitrogen |
| CD33 | PerCp-Cy5.5 | P67.6 | 1:100 | 366615 | AB_2566417 | Biolegend |
| CD34 | AF450/PB | RAM34 | 1:100 | 48-0341-82 | AB_2043837 | Invitrogen |
| CD41 | PB | MWReg30 | 1:100 | 133932 | AB_2750526 | Biolegend |
| CD45 | APC-Cy7-A | 5B1 | 1:100 | 130-133-115 | AB_2905022 | Miltenyi Biotec |
| CD45.2 | PB | 104 | 1:100 | 109820 | AB_492872 | Biolegend |
| CD45.2 | PE | 104 | 1:100 | 109808 | AB_313445 | Biolegend |
| CD48 | PB | HM48-1 | 1:100 | 103418 | AB_756140 | Biolegend |
| CD71 | PE | RI7217 | 1:100 | 113808 | AB_313569 | Biolegend |
| CD117 | APC cy7 | 2B8 | 1:100 | 105826 | AB_1626278 | Biolegend |

| | | | | | | |
|---|---|---|---|---|---|---|
| CD117 | PeCy7 | 104D2 | 1:100 | 25-1178-42 | AB_10718535 | Invitrogen |
| CD150 | APC | TC15-12F12.2 | 1:100 | 115910 | AB_493460 | Biolegend |
| FcER1a | APC | MAR-1 | 1:100 | 134316 | AB_10640121 | Biolegend |
| Gr1 | APC | RB6-8C5 | 1:100 | 108412 | AB_313377 | Biolegend |
| Gr1 | Pe-Cy5 | RB6-8C5 | 1:100 | 108410 | AB_313375 | Biolegend |
| Sca1 | PE cy 7 | E13-161.7 | 1:100 | 122514 | AB_756199 | Biolegend |
| Ter119 | APC | TER-119 | 1:100 | 116212 | AB_313713 | Biolegend |
| Ter119 | Pe-Cy5 | TER-119 | 1:100 | 116210 | AB_313711 | Biolegend |
| | **Application** | | | | | |
| PDE3A | WB, IF | Rabbit polyclonal | 1:500 | ab244337 | | Abcam |
| SLFN12 | WB | EPR20904-32 | 1:100 | ab234418 | | Abcam |
| c-KIT | WB | D13A2 | 1:1000 | 3074 | AB_1147633 | Cell Signaling Techologies |
| p-KIT | WB | Rabbit polyclonal | 1:1000 | 3391 | AB_2131153 | Cell Signaling Technologies |
| Tryptase | IF | AA1 | 1:200 | M7052 | AB_2206478 | Agilent/Dako |
| | **Fluorochrome** | | | | | |
| Goat anti-rabbit | AlexaFluor 594 | Goat polyclonal | 1:200 | A11012 | AB_2534079 | Thermo Fisher Scientific |
| Goat anti-mouse | AlexaFluor 488 | Goat polyclonal | 1:200 | A32723 | AB_2633275 | Thermo Fisher Scientific |
| | **Application** | | | | | |
| Goat anti-rabbit | WB | Goat polyclonal | 1:200 | P0448 | AB_2617138 | Agilent/Dako |
| Goat anti-mouse | WB | Goat polyclonal | 1:200 | P0447 | AB_2617137 | Agilent/Dako |

**Supplementary Table 4. Antibodies used for flow cytometry, Western blotting and immunofluorescence.**

The antibodies used in this study with the respective fluorophores for flow cytometry, clone, catalogue and RRID numbers and suppliers are shown. WB, Western blotting; IF, immunofluorescence.
